# Discovery of a wide variety of α-1,6-cyclized β-1,2-glucan synthases: a new entrance for host-microbe interactions

**DOI:** 10.1101/2025.04.21.649730

**Authors:** Sei Motouchi, Inhyuk Kim, Ai Ishiguro, Kenshiro Niu, Wakana Tachibana, Shiro Komba, Toma Kashima, Yoshinao Kitano, Hiroyuki Nakai, Kenji Kai, Masahiro Nakajima

## Abstract

α-1,6-Cyclized β-1,2-glucans (CβGα) are established virulence factors in *Xanthomonas*, yet functional breadth and distribution of CβGαs in nature have remained unclear. Here, we biochemically identified enzymes synthesizing CβGαs with specific chain lengths, exhibiting potential natural occurrence of various CβGαs. Structural analyses identify subtle variations in a loop, named Loop X, as a key determinant of product size, leading to understanding of the whole picture of the mechanism that controls the sizes. We further demonstrated that CβGα composed of 13 glucose units contributes to plant-virulence in *Ralstonia pseudosolanacearum*, expanding the functional scope of virulence-associated cyclic glucans. Overall, it provides the potential targets for regulating various plant-microbe interactions and also serves as a vital lead to discovering unknown host-microbe interactions with significant potential for agricultural applications.

## Introduction

α-1,6-Cyclized β-1,2-glucan (CβGα) is a glycan synthesized by such as *Xanthomonas campestris* pv. *campestris* (Xcc, phytopathogen), *Ralstonia pseudosolanacearum* (phytopathogen) and *Cereibacter sphaeroides* (photosynthetic bacteria)^1,2^. Xcc, *R. pseudosolanacearum* and *C. sphaeroides* synthesize CβGα with 16 glucose moieties (CβG16α), CβG13α and CβG18α, respectively. Although the physiological function of CβG13α and CβG18α are not identified, CβG16α is known to be essential for pathogenicity but not for survival. CβG16α is responsible for inhibition of PR-1 protein and callose accumulation in plants^3^.

No enzyme responsible for synthesis of CβGα had been identified until the author’s group found that the enzyme from Xcc (XccOpgD), which belongs to the glycoside hydrolase family GH186, converts linear β-1,2-glucan directly to CβG16α^4^. Considering conservation of substrate recognition residues, it is suggested that GH186 homologs from almost all of *Xanthomonas*, which cause disease in approximately 400 crop species worldwide^5^, are synthases of CβG16α^4^.

One of the remarkable features of XccOpgD is that the enzyme employs a very unique reaction mechanism, anomer-inverting transglycosylation for cyclization of β-1,2-glucan by α-1,6-glycosidic linkage. The amino acid residues critical for this reaction mechanism are highly conserved within the GH186 family. In contrast, substrate recognition residues that seem to be responsible for producing a cyclic glucan with a particular chain length are not conserved at all within GH186, implying the diversity of chain lengths of products^4^.

In this perspective, we performed biochemical analyses toward various GH186 homologs from bacteria including *R. pseudosolanacearum, Bradyrhizobium diazoefficiens* (a root-nodule bacterium), and *C. sphaeroides*. We identified eight kinds of unique synthases producing new CβGαs in terms of degrees of polymerization and/or side chains. Furthermore, structural analyses of cyclic tridecaose (13), octadecaose (18), henicosaose (21) and nonacosaose (29) synthases explain how DPs of CβGαs as products by the GH186 enzymes are diversified. This study clearly indicated various bacteria have the potential to produce various CβGαs, leading to a new and wide perspective of symbiosis and pathogenicity of various proteobacteria.

## Result

### Reaction products of GH186 homologs

We focused on the twelve homologs [OpgDs from *Methylomagnum ishizawai* (MiOpgD2, MiOpgD3 and MiOpgD4), *Ralstonia pseudosolanacearum* (RpOpgD2, RpOpgD3), *Cereibacter sphaeroides* (CsOpgD2), *Novosphingobium pentaromativorans* (NpOpgD), *Bradyrhizobium diazoefficiens* (BdOpgD), *Pseudoaltermonas espejiana* (PeOpgD, PeOpgD2), *Shewanwlla oneidensis* (SoOpgD2) *Roseobacter litoralis* (RlOpgD)] based on phylogenetic analysis in the previous study^4^. These homologs were expected to be transglycosylases with different functions from XccOpgD (16, the clarified DP are written in parenthesis if the homologs have specificity for DP of transglycosylation products) because the residues recognizing a moiety released as a cyclic glucan in a substrate are diversified within these homologs. Consequently, these homologs cover most of major clusters classified by Sequence similarity network (SSN) analysis (Figure 1).

**Figure 1.**
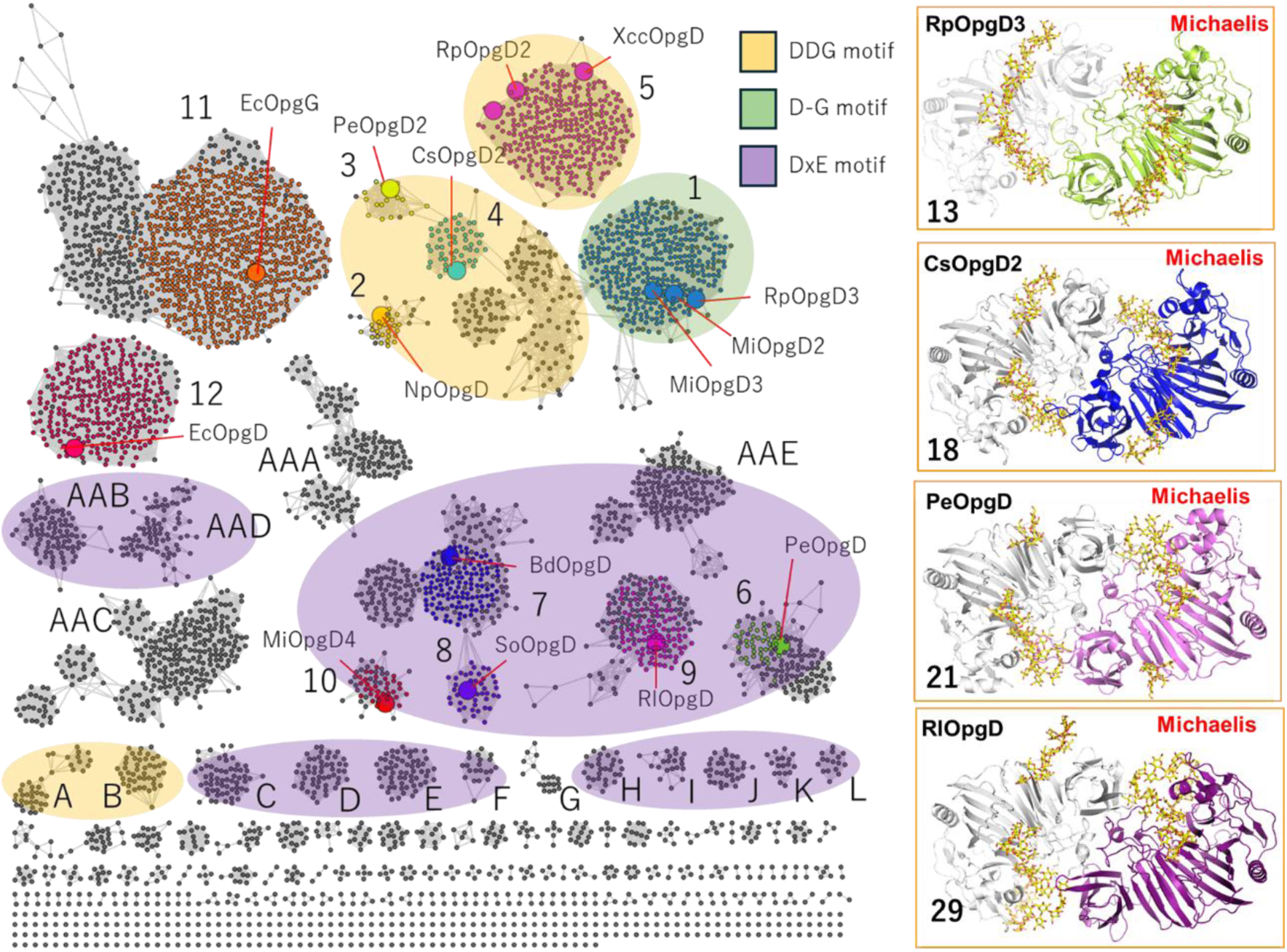
SSN analysis for GH186 family and overall Michaelis complexes. (middle, SSN) The clusters containing proteins which have D-G, DDG or DxE motif at Loop X are shown by semi-transparent colors; green, orange and purple, respectively. This color coding is applied to the clusters with 10 or more nodes. Clusters 1–10 represent the groups which contain the homologs analyzed in this study. Clusters AAA–AAD represent large groups with unknown biochemical function. Clusters A–L represent the medium clusters (which have 10 or more nodes) with unknown biochemical function. (left and right, Michaelis complexes) Chains A of RpOpgD3, CsOpgD2, PeOpgD and RlOpgD are shown as light green, blue, purple and deep purple cartoons, respectively. Each ligand (linear β-1,2-glucan) in the protein is shown as a yellow stick.

First, these homologs were analyzed biochemically. They converted linear β-1,2-glucans to products whose DPs appear to be specific or smeared products on TLC plates. ESI-MS analyses showed that molecular mass of the main products released by all the homologs were approximately 18 less than those of linear β-1,2-glucans (Experiment 0) (Figure 2a, Supplementary Figures 1–12). These results suggest that all the main products were cyclic forms. To elucidate the detailed structures of the products, the products were treated with BtBGL^6^, an exo β-glucosidase from *Bacteroides thetaiotaomicron* which prefers β-1,2-glucooigosaccharides as substrates (Experiment 1) to degrade β-1,2-glucans with only linear form, or both BtBGL and CpSGL^7^, an endo β-1,2-glucanase (Experiment 2) to degrade β-1,2-glucans with both linear and cyclic forms. TLC analyses of the reactants by Experiment 1 showed that the spots of the main products unchanged as they are in many of the enzymes while the smeared products released by some enzymes in Experiment 0 were migrated at one specific spots (Figure 2a). ESI-MS analyses for the reactants by Experiment 1 showed that molecular mass of these specific oligosaccharides were still approximately 18 less than those of linear glucans (Supplementary Figures 1–12), indicating that the products released by Experiment 0 contain cyclic glucan moieties, as suggested above. The products of MiOpgD4 (21 in Experiment 1), RpOpgD2 (16 in Experiment 1), and NpOpgD (16 in Experiment 1) have different mass spectra between Experiment 0 and 1, suggesting that several glucose moieties at the non-reducing end of a substrate remain as a side chain in cyclic products and the side chains are removed by BtBGL (Figure 2b, c, Supplementary Figures 3, 4, 7). It is because BtBGL is an exo β-glucosidase acting on “the non-reducing end”. In contrast, the same molecular mass spectra for the main products in Experiment 0 and 1 of MiOpgD2 (12), MiOpgD3 (13), MiOpgD4 (21), RpOpgD3 (13), CsOpgD2 (18), BdOpgD (21), PeOpgD2 (16-18), PeOpgD (21), SoOpgD2 (21), RlOpgD (29) suggested that these homologs mainly produced cyclic glucans without side chain (Figure 2bc, Supplementary Figure 1-2, 5-6, 8-12). Subsequently, NMR analysis for the cyclic hexadecaose from Experiment 1 of RpOpgD2 was conducted to examine whether BtBGL can degrade a side chain completely or not. Consequently, the cyclic hexadecaose produced by Experiment 1 of RpOpgD2 was demonstrated to be CβG16α (Supplementary Table 1), indicating that BtBGL can remove a side chain in a CβGα completely without being sterically hindered. Thus, a DP of a main product after Experiment 1 represents a DP of a cyclic moiety.

**Figure 2.**
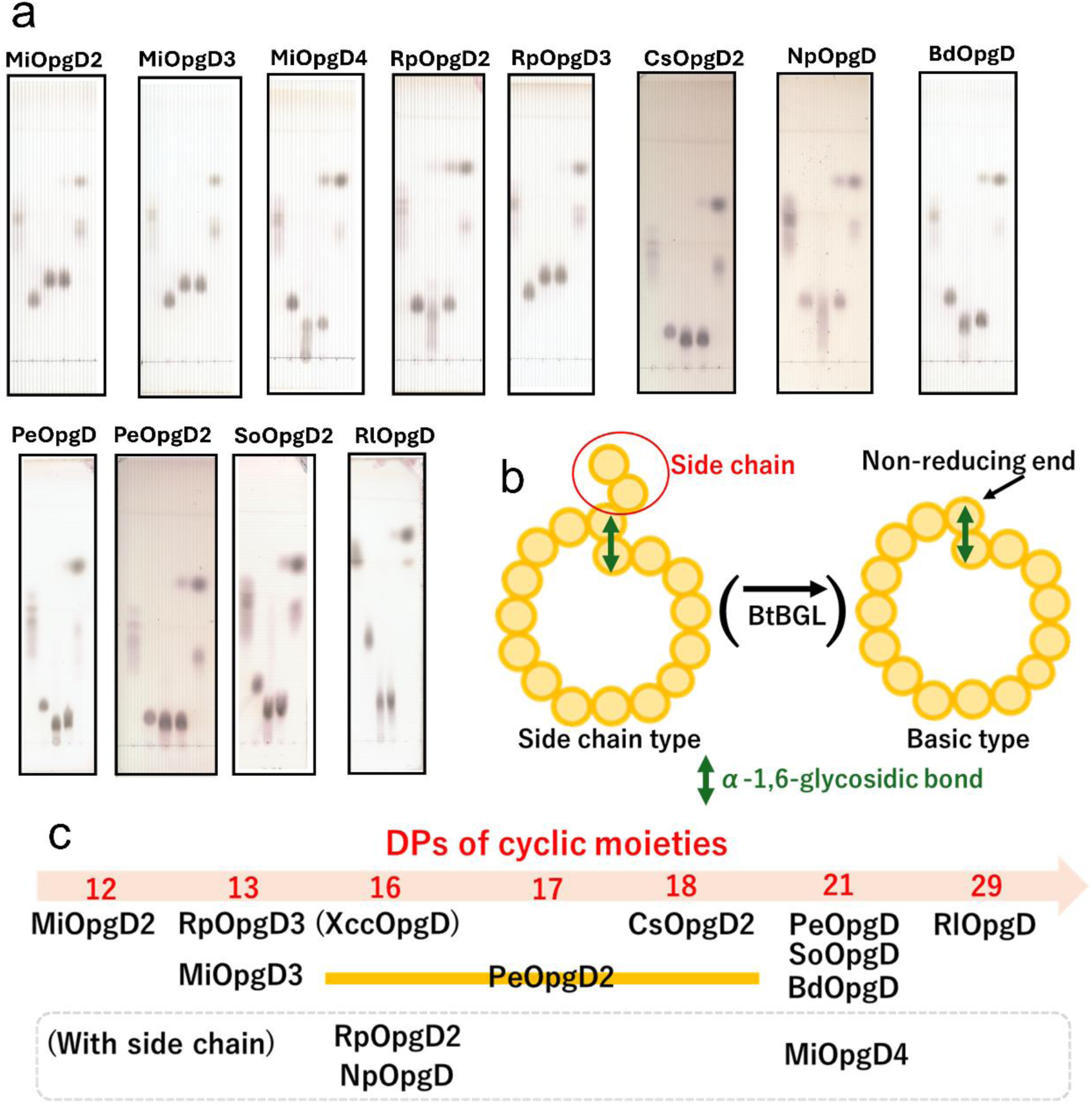
Reaction products from linear β-1,2-glucan by GH186 homologs. (a) TLC analysis of reaction products. In each TLC plate, lanes from left to right are as follows; 1st, β-1,2-glucooligosaccharide marker (DP3–7); 2nd, CβG16α (1%); 3rd, reaction products produced from linear β-1,2-glucans (average DP of 121, 1%) as substrates; 4th, samples of the 3rd lane treated with BtBGL; 5th, samples of the 3rd lane treated with BtBGL and CpSGL. (b) Schematic representation of basic type and side chain type of CβG16α. (c) Summary of the main products of the GH186 homologs analyzed. See Supplementary Figs. 1–12 for detail of each homolog.

The previous study of XccOpgD showed that an oligosaccharide detected by TLC after Experiment 2 was a linear pentaose containing one α-1,6-glycosidic linkage^4^. Experiment 2 for all the enzymes in this study showed the same patterns as that of XccOpgD (Figure 2a, Supplementary Figure 1-12c). Therefore, all the cyclic moieties of products are suggested to be CβGα. To clarify the structure of the products by MiOpgD2, RpOpgD3, CsOpgD2 and PeOpgD, NMR was conducted. Consequently, the main compound produced by each homolog were demonstrated to be CβG12α, CβG13α, CβG18α and CβG21α, respectively (Supplementary Figure 1, 4-6, 9g, h).

Overall, MiOpgD2, MiOpgD3, MiOpgD4, RpOpgD3, CsOpgD2, BdOpgD, PeOpgD, SoOpgD2, and RlOpgD synthesized cyclic dodecaose (12), tridecaose (13), henicosaose (21), tridecaose (13), octadecaose (18), henicosaose (21), henicosaose (21), henicosaose (21) and nonacosaose (29), respectively (Figure 2bc). PeOpgD2 synthesized cyclic hexadecaose and heptadecaose mainly and cyclic octadecaose as a minor product. MiOpgD4, RpOpgD2 and NpOpgD synthesized cyclic henicosaose with a side chain, cyclic hexadecaose with a side chain, cyclic hexadecaose with a side chain, respectively (Figure 2bc).

In addition, SSN analysis for GH186 showed that groups classified by SSN tend to be clustered depending on DP specificities of products (Figure 1). Detailed categories were mentioned based on structural analyses described below.

### Enzymatic profiles

Optimal pH and temperature were examined for each homolog by TLC analysis (Supplementary Figures 1–12d, e). Substrate specificity for each homolog was investigated under the optimal condition that adequate transglycosylation activity was detected. As a result, all homologs in this study were specific to linear β-1,2-glucan among tested polysaccharides (Supplementary Figures 1–12f). Thus, specific activities of RpOpgD3, CsOpgD2 and RlOpgD were analyzed using linear β-1,2-glucan (average DP of 121). The values at substrate concentration, 0.39 % (w/v), are 1.27, 0.11 and 0.018 U/mg, respectively. Considering that the specific activity of XccOpgD at same substrate concentration is 0.29 U/mg^4^, it appears that as the DP of the synthesized CβGα increases, the specific activity tends to decrease.

### Michaelis complexes

D379 in XccOpgD, the general acid catalyst in the anomer-inverting transglycosylation^4^, is conserved among all homologs analyzed in this study (Supplementary Figure 13). Therefore, we assumed that general acids are common in GH186 transglycosylases, and mutants of potential general acid residues were used to obtain Michaelis complexes of several homologs in this study. Consequently, substrate-enzyme complex structures of RpOpgD3 (13), CsOpgD2 (18), PeOpgD (21) and RlOpgD (29) were obtained at 1.57, 2.40, 2.20, 1.83 Å, respectively (Figure 1, Supplementary Table 2). Each asymmetric unit of CsOpgD2 (18) and RlOpgD (29), contains two molecules with almost the same conformation, and a molecule is observed in an asymmetric unit of RpOpgD3 (13) and PeOpgD (21). Chains A were used for discussing the catalytic clefts of the homologs with a dimer in an asymmetric unit [CsOpgD2 (18) and RlOpgD (29)].

In the Michaelis complex of XccOpgD (16), the closed conformation of the catalytic cleft is achieved by α-helix 3 approaching the substrate in the cleft^4^. RpOpgD3 (13) and CsOpgD2 (18), PeOpgD, RlOpgD (29) also seem to form closed conformations in the presence of a substrate, based on superposition with the Michaelis complex of XccOpgD (Supplementary Figure 14), implying that the closure motion of α-helix 3 is shared in GH186.

### Substrate binding modes

The electron densities of linear β-1,2-glucans were clearly observed in the complex structures of RpOpgD3 (13), CsOpgD2 (18), PeOpgD (21) and RlOpgD (29) (Supplementary Figure 15a). Each substrate in the complex structure is rounded on the side of the non-reducing end. The linearly arranged moieties in the substrates are firmly grabbed by the enzymes, while the other rounded moieties look pinched softly (Supplementary Figure 15b).

First, we defined subsite positions of the newly obtained complex structures (subsite is the nomenclature used for substrate binding sites; see https://www.cazypedia.org/index.php/Sub-site_nomenclature for details). Based on the subsite positions of XccOpgD, subsites −3 to +4 and the sequential three subsites from the non-reducing ends^4^ are well superimposed among all the complex structures. In particular, the distorted conformations (^1^*S*_3_) of the Glc moieties in the middle of the catalytic pockets are well superimposed (Supplementary Figure 16ab). Such distortion indicates that these moieties are subsites −1, namely cleavage sites, because the distortion generally causes anomeric position to be pseudoaxial, which is suitable for nucleophilic attack to an anomeric carbon atom^8^. Thus, the origins of subsite positions (−1 and +1) of the four newly solved structures are the same as XccOpgD.

According to the defined subsite positions, the rounded moieties in the substrates are at subsites −1 to −13, −18, −21 and −29 in RpOpgD3 (13), CsOpgD2 (18), PeOpgD (21) and RlOpgD (29), respectively (Supplementary Figure 15a). This observation is consistent with the DP specificities of the reaction products. 6-Hydroxy groups at the non-reducing end glucose moieties in the complex structures are located at appropriate positions for nucleophilic attack to the anomeric carbon atoms (Supplementary Figure 17), suggesting that the reaction mechanism of 4 homologs is fundamentally the same as that of XccOpgD^4^ (See Supplementary note 1).

The shapes of the bound substrates are clearly different from each other depending on the DP specificities. The key subsite is −5, which is classified into three types (Supplementary Figure 18); <1> CsOpgD2 (18) has a distorted glucose moiety as well as XccOpgD, <2> RpOpgD3 (13) has a glucose moiety with a usual (chair, ^4^*C*_1_) conformation although dihedral angles around the glycosidic bond with subsite −4 are unusual, according to Perić-Hassler, L. et al.^9^, <3> In PeOpgD (21) and RlOpgD (29), both the conformations and the dihedral angles are within usual ranges. Details of differences in substrate binding modes are discussed below.

### The principal determinant for DP specificity of the reaction products

One of the remarkable feathers of the GH186 enzymes in this study is that the enzymes produced the cyclic glucans with specific range of DPs in the cyclic moieties. Combination of structural observation and sequence analysis unveiled a key loop for DP specificity of the products. Hereafter this loop is called Loop X (342 a.a–344 a.a. in XccOpgD). Loop X is located on the opposite side of the catalytic center in the cyclic moiety of the ligand (Figure 3bc). In Loop X, a motif composed of two or three residues is considered to be the principal determinant. This motif is classified into three types; D–G, DDG and DxE, which corresponds to the classification based on subsite −5 described above. The first Asp conserved in the three types of Loop X commonly interact with the third glucose moieties from the non-reducing end, which are important for stabilizing catalytic pathway. The Asp residues and the third glucose moieties are well superimposed with each other (Figure 3a). Therefore, differences in DP specificities of the products mainly depend on the second and third residues of loop X.

**Figure 3.**
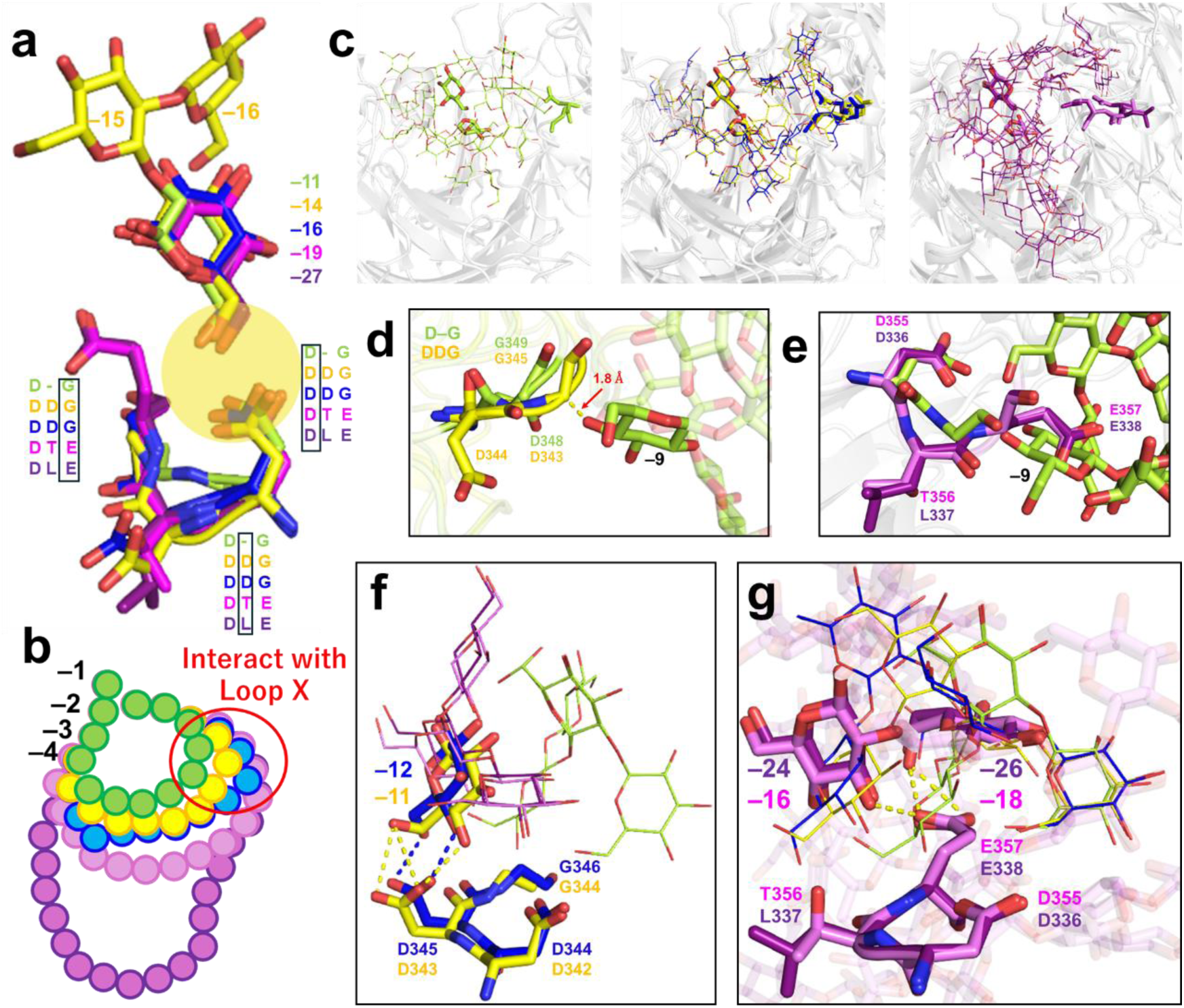
Superposition of Loop X among the Michaelis complexes of XccOpgD and the homologs in this study. The Michaelis complexes of XccOpgD (chain B) and the GH186 homologs in this study (chains A) are superimposed based on residues in the catalytic centers. Complex structures of XccOpgD, RpOpgD3, CsOpgD2, PeOpgD and RlOpgD are distinguished by color; yellow, light green, blue, purple and deep purple, respectively. Subsite numbers are labeled appropriately as needed. Dotted lines without labels represent hydrogen bonds with appropriate distances. (a) Third glucose moieties from the non-reducing end and Loop X of XccOpgD, RpOpgD3, CsOpgD2, PeOpgD, RlOpgD are shown as sticks. The first and second glucose moieties from the non-reducing end are also shown as sticks in XccOpgD. These glucose moieties are labelled with subsite numbers in the complexes. Two or three residues in Loop X are shown as sticks. The residues in the aligned sequences are boxed to indicate corresponding residues. (b) The schematic diagram of superposition of linear β-1,2-glucans in the Michaelis complexes. Glucose moieties at plus subsites are omitted. (c) Overall view of the position of loop X. Protein moieties in the complexes are shown as grey semi-transparent cartoons. The residues of Loop X and the glucose moieties at subsite −1 and the non-reducing end in the complexes are shown as sticks. The other part of the linear β-1,2-glucans are shown as lines. (left) RpOpgD3, (middle) XccOpgD and CsOpgD2 are superimposed. (right) PeOpgD and RlOpgD are superimposed. (d) Steric hindrance between the glucose moiety at subsite −9 of RpOpgD3 and Loop X of XccOpgD. (e) Steric hindrance between the glucose moiety at subsite −9 of RpOpgD3 and Loops X of PeOpgD and RlOpgD. (f) Specific glucose moieties recognized by the second Asp of DDG-motif in XccOpgD and CsOpgD2. The glucose moieties in the five complexes are shown as lines. Only glucose moieties which interact with the second Asp on Loop X in XccOpgD and CsOpgD2 complexes are changed to stick representation. The residues of the DDG-motifs are shown as sticks. (g) Specific localization of glucose moieties of PeOpgD and RlOpgD complexes by the interactions with third Glu of DxE-motif in Loop X. The glucose moieties at subsites −16 and −18 in PeOpgD and subsites −24 and −26 in RlOpgD, and Loop X are shown as sticks. The other glucose moieties in the complexes of PeOpgD and RlOpgD are shown as semi-transparent sticks. The glucose moieties from the common subsites (subsites –19 of PeOpgD) to the subsites near the DxE-motif in Loop X of RpOpgD3, XccOpgD and CsOpgD2 are shown as lines.

In the D–G motif of RpOpgD3 (13), the second residue is missing. which makes the Loop X short. Such shortness makes it possible to avoid steric hindrance with the glucose moiety at subsite −9. In contrast, it is clear that this glucose moiety collides with E357 in the DxE region of PeOpgD (21) potentially (Figure 3e). Gly344 in the DDG region of XccOpgD (16) is too close to the glucose moiety as well (Figure 3d). This observation suggests that deletion of the second residue is needed to be specific to DP13 of the product. The D–G motif is conserved in cluster 1 including MiOpgD2 (12) (Supplementary Figure 19). Overall, the D–G region is an indicator for small cyclic glucan synthases (DPs of 12–13).

In XccOpgD and CsOpgD2 (18), DDG motif is conserved at Loop X, and the second Asp residues form two hydrogen bonds with subsites −11 (XccOpgD) and −12 (CsOpgD2). In contrast, no glucose moiety is observed at the corresponding positions in the enzymes producing CβGαs with DPs of 13, 21 and 29 (Figure 3f). Incompatibility with DP of 13 is due to the potential steric hindrance described in the previous paragraph. Potential collision with CβGα with DPs 21 and 29 is not observed. It is probably because most of the DxE motif has Leu (major) or Thr (minor) as the second residue, which can provide weaker interactions than Asp potentially. Overall, the second Asp is suggested to be a critical mark indicating DP specificity to 16–18 and not to 21 or higher. The DDG motif is highly conserved among cluster 2, 3, 4, 5, A and B (Figure 1, Supplementary Figure 19).

The loop X of PeOpgD (21) and RlOpgD (29) have the motif DxE. Superimposition between the structures of PeOpgD and RlOpgD revealed that the Glu residues in the motif interact with subsite −18 (PeOpgD) and −26 (RlOpgD) in each homolog (Figure 3g). These interactions are starting points of the significantly different flow of substrate toward the reducing end compared with the other enzymes with D–G or DDG motifs (Figure 3g). Sequence analysis showed that all the clusters which contain the synthases of henicosaose or larger glucans (BdOpgD, MiOpgD4, SoOpgD2, PeOpgD, RlOpgD) have this motif (Figure 1, Supplementary Figure 19). These results suggest that the DxE motif is an indicator for predicting large cyclic glucan (larger than henicosaose) synthase.

In summary, the motif in Loop X is an indicator of DP ranges of the products in the cyclic moiety (12–13, 16–18, and 21 or higher). Other elements are needed for the GH186 enzymes to be specific to particular DPs, which is discussed below.

### Co-determinant for specificity to DPs 12–13

There is a highly conserved Gly residue [Gly77 in RpOpgD3 (13)] that should be noted among the cluster of the D–G motif (Figure 4a-c, Supplementary Figure 20). This is a unique feature that is not found in DDG and DxE motif enzymes. The Gly residue is located in α-helix 3, and the helices are well superimposed among complex structures except for XccOpgD (Figure 4a-c). Because the Gly residue is located vicinity of a glucose moiety at subsite −6, substitution with the other residues is not acceptable due to the potential collision with the substrate (Figure 4a-c). Altogether, it is considered that the Gly residue in α-helix 3 is introduced with the D–G motif by co-evolution to be specific to small DPs of the products.

**Figure 4.**
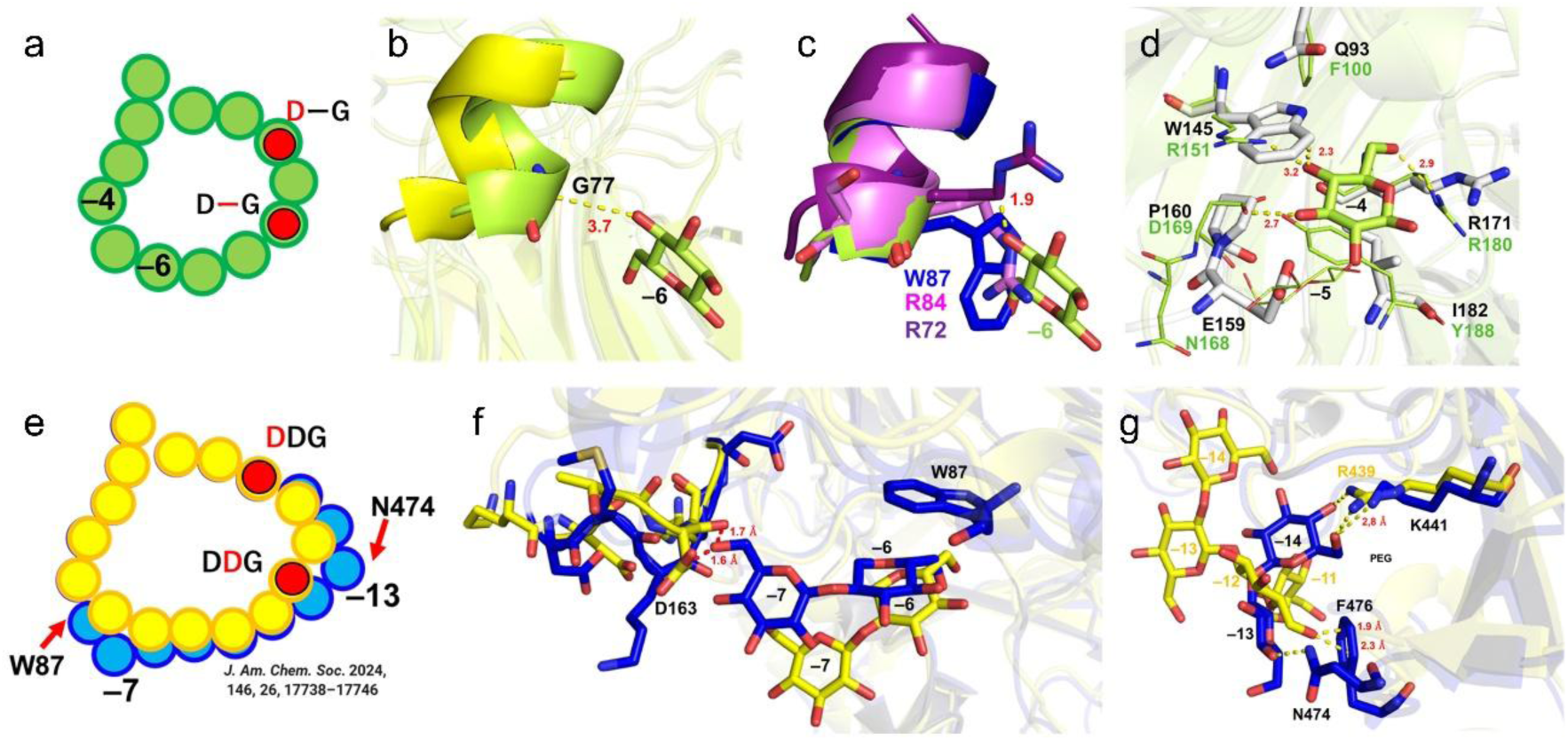
The features of catalytic clefts and substrate binding mode of D-G-motif and DDG-motif GH186 homologs. Michaelis complexes of XccOpgD (chain B), the GH186 homologs in this study (chains A) and the structure of MiOpgD2 predicted by Alphafold3 are superimposed based on residues in the catalytic centers. The complex structures of XccOgpD, RpOpgD3, CsOpgD2, PeOpgD and RlOpgD and MiOpgD2 are shown in yellow, light green, blue, purple, deep purple and grey, respectively. Subsite numbers are labeled appropriately as needed. Dotted lines without labels show hydrogen bonds with appropriate distances. (a) The schematic diagram of a linear β-1,2-glucan in the catalytic cleft of RpOpgD3. (b) Superposition of α-helix 3 between RpOpgD3 and XccOpgD. G77 is shown as a stick. (c) Superposition of α-helix 3 between RpOpgD3, CsOpgD2, PeOpgD and RlOpgD. W87 of CsOpgD2, R84 of PeOpgD and R72 of RlOpgD are shown as sticks. The latter two residues sterically collide with the glucose moiety at subsite −6 of RpOpgD3. (d) Comparison of substrate recognition residues around subsite −4 of RpOpgD3 between RpOpgD3 (PDB ID: 22PV) and ligand-free predicted structure of MiOpgD2. (e) The schematic diagram of superposition between linear β-1,2-glucans in the catalytic clefts of XccOpgD and CsOpgD2. (f) Superposition between XccOpgD and CsOpgD2 complexes at subsites −6 and −7. (g) Comparison of positions of subsites −13 and −14 between XccOpgD and CsOpgD2.

To understand how DPs of 12 and 13 are distinguished, a binding mode of a linear β-1,2-glucan with MiOpgD2 was predicted using AlphaFold3^10^. The closed conformation of MiOpgD2 was obtained although no plausible binding mode was obtained unfortunately. Thus, the predicted structure without the ligand was superimposed with RpOpgD3 (13). Interestingly, substitution of the four residues interacting with a glucose moiety at subsite −4 and insertion of one residue potentially were observed (Figure 4d).

The four residues are highly conserved almost throughout the GH186 family whereas only a small group in the cluster 1 has the substituted and inserted residues (Supplementary Figure 20).

Substitution of Tyr188, and Asp169 and Arg151 is expected to cause the loss of stacking and hydrogen-bond interactions, respectively, at subsite −4 (Figure 4d). Substitution of Phe to Gln (Phe100 in RpOpgD3 (13)) is thought to be accompanied with the substitution of Arg to Trp (Arg151 in RpOpgD3 (13)) because Trp145 in MiOpgD2 (12) collides with Phe100 in RpOpgD3 (13) potentially (Figure 4d). The inserted residue of E159 in MiOpgD2 (12) clearly collides with a glucose moiety at subsite −5 (Figure 4d). Overall, it is predicted that glucose moieties at subsites −4 and −5 are pushed outside to reduce one glucose moiety in a product. This complicated simultaneous change of residues is also considered as a result of co-evolution.

### Co-determinant for specificity to DPs 16–18

To understand the DP specificities of the product within the range of DPs 16–18 in the cyclic moieties, complex structures of XccOpgD and CsOpgD2 (18) are compared. The substrate binding modes are totally different between them at a glance (Supplementary Figure 21). However, only one glucose moiety in each enzyme is superimposed well at a minus subsite beyond −5. The glucose moieties (at subsite −11 in XccOpgD and −12 in CsOpgD2) are recognized by the second Asp residues in the DDG motif in both enzymes (Figure 3f). Thus, the position of the glucose moiety is the fundamental subsite for the enzymes to produce CβGαs with DPs 16–18. Because one subsite is added in each side of the fundamental subsite in CsOpgD2 (18) compared with XccOpgD, at least two elements are needed to change the specificity.

The one is Trp87 in CsOpgD2 which corresponds to Gln80 in XccOpgD (Figure 4e, f). The Trp residue forms CH-π stacking interaction with a glucose moiety at subsite −7. This interaction clearly changes the orientation of the glucose moiety, resulting in totally different positioning of subsite positions from that of XccOpgD (Figure 4e, f). The other element is N474 and F476 in CsOpgD2, which interact with a moiety at subsite −13 (Figure 4g). It should be noted that potential steric hindrance is not observed between CsOpgD2 and the ligand in XccOpgD. Substitution of the three residues changes DP preference, which is consistent with the fact that CsOpgD2 produced CβGα with DP of 16 as a minor product (Supplementary Figure 6). This structural observation is also consistent with the product preferences of PeOpgD2 (16-17) producing CβGα with DPs of 16 and 17 as main products and production of CβGα containing a cyclic hexadecaose unit by NpOpgD and RpOpgD2. Trp87 in CsOpgD2 are conserved in PeOpgD2 (16-17) while Asn474 and Phe476 are not. The three residues in CsOpgD2 (Trp87, Asn474, and Phe476) are not conserved in NpOpgD nor RpOpgD2 (Supplementary Figure 13).

### Co-determinant for specificity to DPs 21 and 29

A unique feature of substrate binding modes in PeOpgD (21) and RlOpgD (29) is that subsite positions on the minus side from −6 are heading toward completely different directions from those of the other complexes with the D–G or DDG motifs (Figure 5a). Instead, access to the spaces corresponding to subsites −6 and −7 in PeOpgD (21) and RlOpgD (29) are prohibited by protruding loops (Figure 5a).

**Figure 5.**
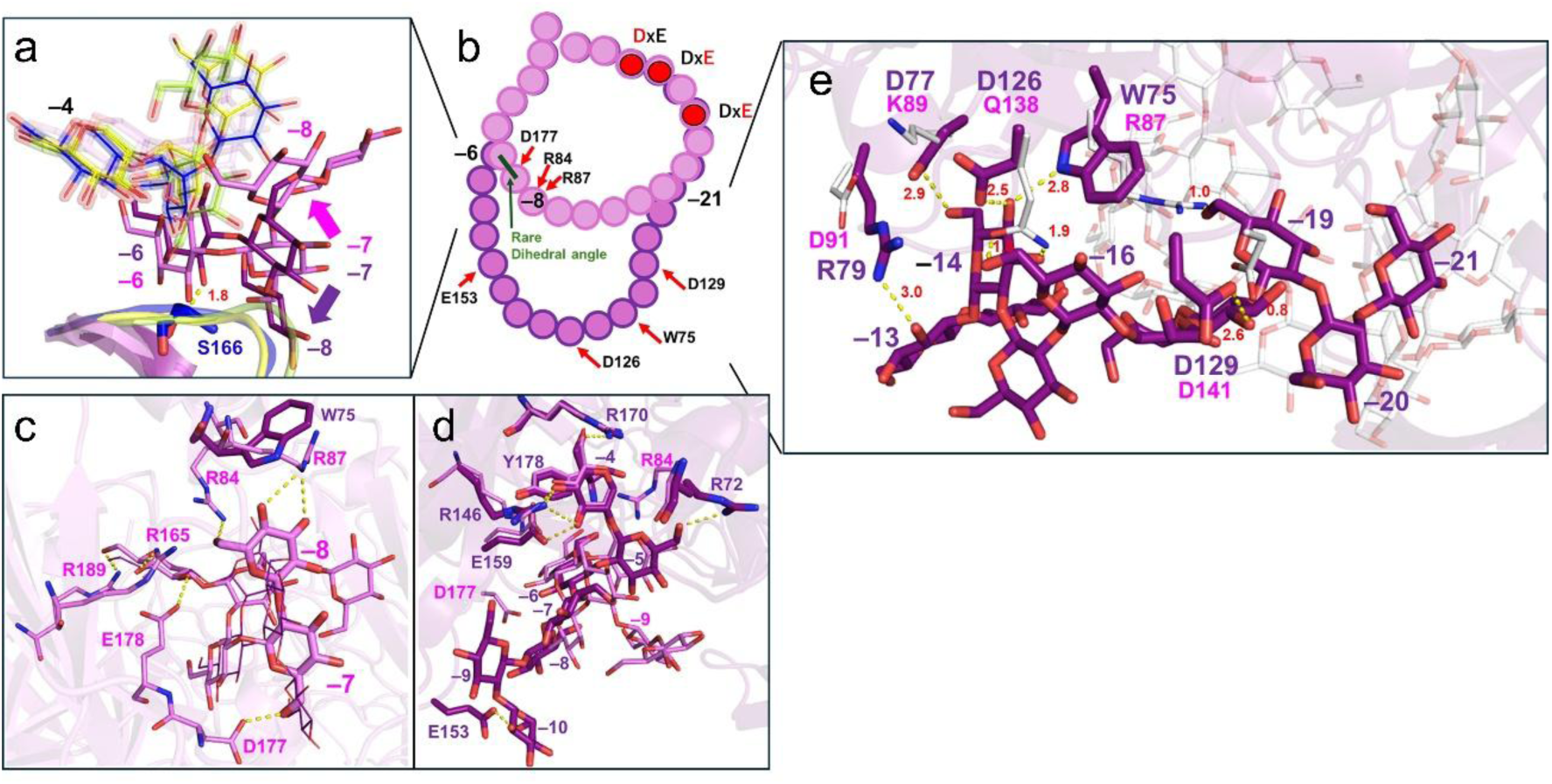
The feature of catalytic clefts and substrate binding mode of DxE-motif GH186 homologs. Complex structures of XccOpgD, RpOpgD3, CsOpgD2, PeOpgD and RlOpgD are distinguished by color; yellow, light green, blue, purple and deep purple, respectively. (a) Substrate binding modes in the GH186 homologs with DxE-motifs around subsite −6. The glucose moieties of RpOpgD3, XccOpgD and CsOpgD2 complexes near subsites −6 in PeOpgD and RlOpgD complexes are shown as lines with semi-transparent sticks. Subsites −4 and −5 in PeOpgD and RlOpgD complexes are shown as semi-transparent sticks. Subsites −6 to −9 of PeOpgD complex and subsites −6 to −8 of RlOpgD are shown as sticks. (b) The schematic diagram of superposition between linear β-1,2-glucans in the catalytic clefts of PeOpgD and RlOpgD complexes. (c) Substrate recognition at subsites −4 to −9 of PeOpgD complex. The glucose moieties at subsites −4 to −9 and recognition residues in PeOpgD complex and W75 in RlOpgD complex are shown as sticks. The glucose moieties at subsites −4 to −9 are shown as lines. (d) Substrate recognition at subsites −4 to −10 of RlOpgD complex. The glucose moieties at subsites −4 to −10 of RlOpgD and recognition residues in RlOpgD complex, and R84 and D177 in PeOpgD are shown as sticks. Subsites −4 to −9 of PeOpgD are shown as lines. (e) Substrate recognition at subsites −13 to −21 of RlOpgD complex. The glucose moieties at subsites −13 to −21 of RlOpgD and recognition residues in RloOpgD complex are shown as sticks. R87, K89, D91, Q138 and D141 in PeOpgD are shown as thin white sticks. The substrate at catalytic cleft of PeOpgD is shown as thin semi-transparent white stick.

A remarkable difference between PeOpgD (21) and RlOpgD (29) in substrate binding mode is that they have a flipped glucose moiety at subsite −7 from each other. Subsite −7 is considered to be an important branching point between them because there is no noticeable recognition residue around the other branching point (subsites −14 in PeOpgD and −22 in RlOpgD). In PeOpgD (21), a glucose moiety at subsite −8 is firmly recognized by Arg 84 and Arg87 (Figure 5b, c, d). It is probably because an orientation of the glucose moiety at subsite −7 is unusual unlike in the case of RlOpgD (29) based on the dihedral angles around glycosidic bonds. Arg87 is a unique residue among the clusters containing henicosaose-producing enzymes (Supplementary Figure 20). Thus, Arg87 is considered to be one of the key factors to produce henicosaose. In RlOpgD (29), this residue is substituted with a Trp residue (Trp75) which is a majority only in cluster 9 (Supplementary Figure 20).

Another residue recognizing subsite −7 (and subsite −6) in PeOpgD (21) is Asp177. This residue is likely to contribute to orientating the glucose moiety at subsite −7 appropriately because this residue does not interact with the corresponding glucose moiety in RlOpgD (29) potentially. This residue is also highly conserved in the same clusters as described for Arg87.

Beyond subsite −8 in PeOpgD (21), Glu357 in the DxE motif strongly recognizes the glucose moieties at subsites −16 and −18. Glu357 is the next residue that newly appears as a highly conserved residue when tracing the subsites in the direction of minus side, which suggests importance of Arg87 and Glu357 (Figure 3g, 5bc). In RlOpgD (29), the binding mode of this region is totally different, which is supported by several recognition residues such as W75, D77, R79, D126 and D129 (Figure 5). W75, D126 and D129 are conserved within most homologs in the cluster 9 (Supplementary Figure 20) and interact with glucose units at subsites −14, −16 and −18 (Figure 5e). This observation suggests that these three residues contribute to branching from the CβGα21 in PeOpgD (21) at subsite −21 in RlOpgD. In addition, the nonacosaose moiety in RlOpgD (29) collides with this region in PeOpgD2 (21) potentially (Figure 5e, Q138, R87, D141), which may also contribute to difference in specificity to DPs of the products. Nevertheless, variation of recognition residues at subsites −13 to −18 in RlOpgD implies further diversity of DPs of CβGαs as products within cluster 9.

### Deletion of *RSp0262* resulted in the loss of CβG13α production and virulence

To confirm that CβG13α is synthesized by the enzyme encoded by *RSp0262* (*RpopgD3*) in *R. pseudosolanacearum*, we generated an in-frame deletion mutant of this gene (Δ*RSp0262*) by double crossover recombination using the pK18mobsacB vector^11^. The deletion of the target gene was confirmed by PCR using genomic DNA as a template. Next, to investigate the production of CβG13α, the cell extracts from Δ*RSp0262* and GMI1000 were prepared and analyzed by LC/MS (Fig. 6a). Consequently, an [M+Na]^+^ ion derived from CβG13α was detected from the wild-type strain GMI1000, but not from Δ*RSp0262*. The complementation of the gene in Δ*RSp0262* using a vector restored CβG13α production. The gene deletion and complementation had no effect on bacterial growth. This indicated that *RSp0262* acts as a biosynthetic gene of CβG13α in *R. pseudosolanaceaerum*.

**Figure 6.**
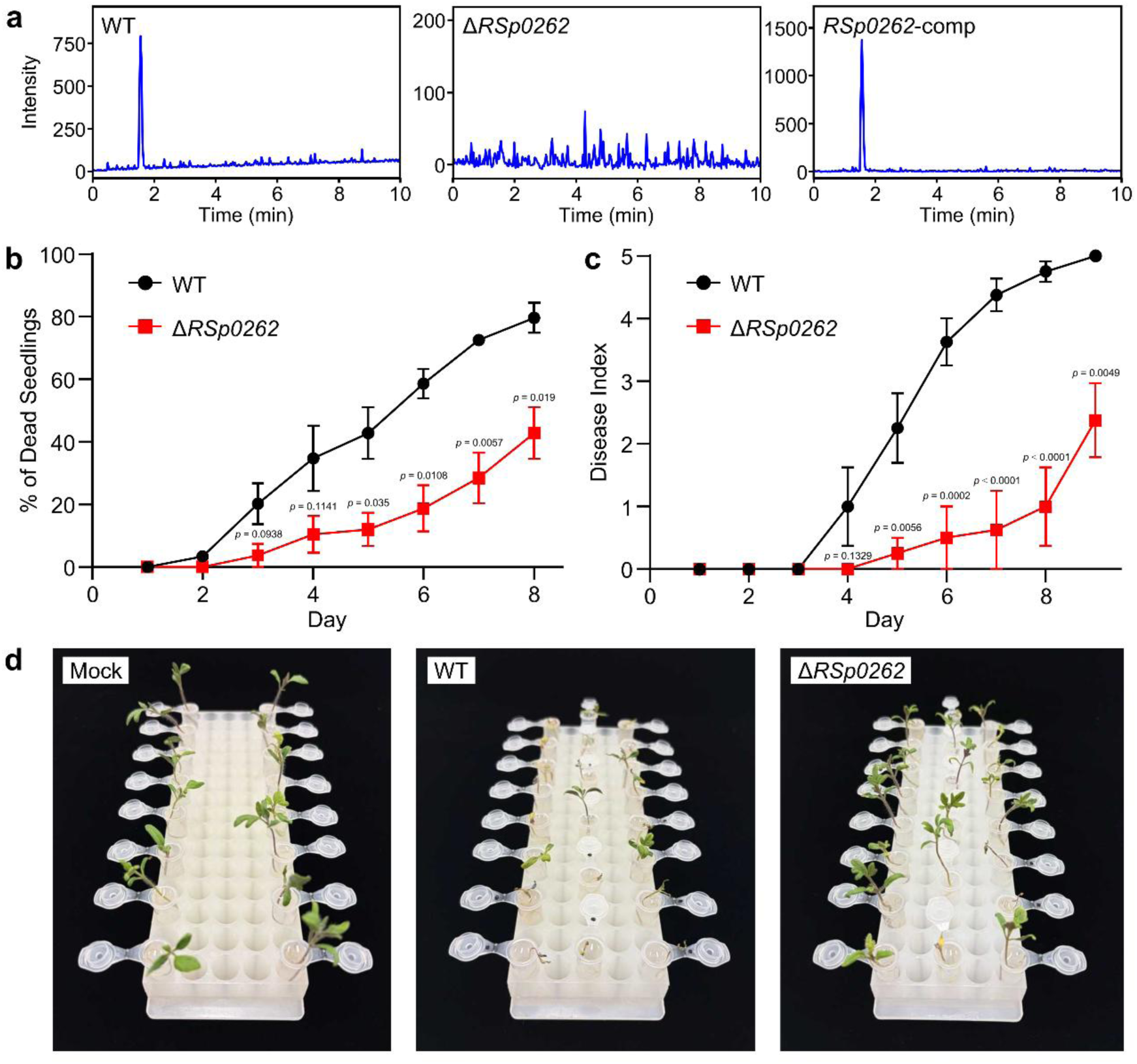
Contribution of CβG13α, produced by the enzyme encoded by *RSp0262*, to *Ralstonia* virulence. (a) LC/MS comparison of CβG13α productivity in wild-type strain, Δ*RSp0262*, and its gene-complemented strain (*RSp0262*-comp). (b) Comparison of virulence between wild-type strain and ΔRSp0262 on tomato seedlings. The *p*-values from the student’s *t*-test are shown. (c) Comparison of virulence between wild-type strain and ΔRSp0262 on tomato plants. The *p*-values from the student’s *t*-test are shown. (d) The tomato seedlings in the virulence assay.

Next, to investigate the contribution of CβG13α to *R. pseudosolanacearum*’s virulence, Δ*RSp0262* was inoculated into tomato seedlings to examine the expression of disease symptoms. This time, the methods of infection via the roots were adopted for tomato plants at different stages^12^. In assays using tomato seedlings, Δ*RSp0262* caused a significantly lower incidence of wilting compared to the wild-type strain (Fig. 6b and d). Furthermore, assays using three-week-old tomato plants similarly showed a significantly reduced degree of wilting symptom compared to the wild-type strain (Fig. 6c). No clear symptoms appeared even after treating tomato seedlings with CβG13α (50–200 μg/mL). These results indicated that the production of CβG13α is important for *R. pseudosolanaceaerum* to express full virulence on plants.

## Discussion

The Xcc CβG16α acted to suppress host defenses both locally and systemically, implying that it may be a key virulence factor in establishing the host–microbe interaction. It has been speculated that the mechanism of CβG16α action differs from those of previously reported plant immune system inhibitors^3^. However, nearly two decades after its activity was first demonstrated, the molecular details remain unclear. In this study, we identified eight additional CβGα synthases, expanding the known repertoire of cyclic glucans among proteobacteria, including *B. diazoefficiens* and *R. pseudosolanacearum*. Moreover, we show that CβG13α is essential for full virulence of *R. pseudosolanacearum* on tomato plants. Notably, treatment with CβG13α alone was not phytotoxic to tomato seedlings, suggesting that this glucan may interact with the plant immune system in a manner analogous to Xcc-derived factors, thereby contributing to infection establishment. Collectively, these findings imply that other CβGα members may also participate in inter-kingdom interactions and influence virulence through similar host–microbe signaling mechanisms.

Furthermore, this study provides fundamentally novel structural insights into the diversity of cyclic glucan-synthesizing enzymes. Cyclic glucan-synthesizing enzymes typically produce cyclic glucans with a certain range of DPs. Although cycloalternan is produced specifically with DP4 by isomaltosyltransferase^13^, cyclodextrin glucanotransferases produce cyclodextrins (DP 6–8) with a preference for a DP, rather than through strict DP specificity^14^. Larger cyclic glucans produced by 4-α-glucanotransferases^15^ as well as cyclic β-1,2-glucan synthases^16,17^, β-1,6- and β-1,3-cyclic glucan synthases^18^ also possess broader ranges of DPs. We previously reported XccOpgD, an enzyme that specifically synthesizes a DP16 cyclic glucan^4^, however, this remained an isolated case. The present study identifying a diverse array of GH186 homologs expanded the discovery of XccOpgD into a comprehensive paradigm for enzymatic synthesis of large cyclic glucans with specific DPs. We demonstrated that such DP specificity is governed by several distinct and systematic structural patterns. GH186 employs a “pinning” mechanism at strategic positions along the linear glucan chain. This approach avoids excessive recognition while effectively defining a size of a cyclic moiety. Particularly notable features are “two pins”: Loop X and the structural environment surrounding subsite –5. Loop X recognizes the outer side of the cyclic glucan slightly distal to the active center; the classification of this region into three distinct sequence patterns dictates the presence or absence of physical constraints, such as steric hindrance, effectively restricting DPs of the products to a narrow range. Furthermore, the local environment around subsite –5 plays a critical role in dictating the trajectory of the glycan chain beyond subsite –6. The binding modes of the glucose units at subsite –5 are classified into three types: standard orientation and conformation, distorted conformation, and flipped orientation, which corresponds precisely to the classification by LoopX. We propose that the synergy between these strategic “pins” and their co-determinants enables the precise synthesis of a specific DP within a high DP range. This work provides a foundational design framework for engineering of enzymes to produce larger cyclic glucans with a specific DP.

Furthermore, by integrating our structural insights into the key residues governing specific DP ranges with Sequence Similarity Network (SSN) analysis, we are now able to predict the product specificity of the vast majority of uncharacterized GH186 homologs. This predictive framework provides significant clues toward deciphering thousands of bacterial phenotypes. For instance, we can now infer that bacteria harboring synthases for CβG13α and/or CβG16α engage in specific plant-microbe interactions, similar to those observed in *Ralstonia* and *Xanthomonas* species. These findings represent a major milestone, paving the way for the systematic identification of host-microbe interactions that could be leveraged for future agricultural applications.

Overall, a wide variety of CβGα synthases gives us a new perspective that CβGαs have various physiological roles. As we partly showed, more plant-microbe interactions related to CβGα are expected for some soil and/or plant-related bacteria. Existence of CβGα synthases from marine bacteria and photosynthetic bacteria and extensive distribution of GH186 homologs in proteobacteria also raises expectation of totally different roles from CβGαs produced by plant-related bacteria. Our finding is a clear milestone for understanding CβGαs and a new horizon for the world of cyclic glycans.

## Methods

### Cloning and purification of recombinant enzymes

Each of the gene encoding the target GH186 proteins were amplified by PCR with the primer pair shown in Supplementary Table 3 using KOD One PCR Master Mix (TOYOBO) and an appropriate genomic DNA as a template. The target genes are as follows: RSP_3187 (GenBank: ABA80803.1) from *C. sphaeroides* (DSM158, type strain), RSc_3057 (GenBank: CAD16766.1) and Rsp0262 (GenBank: CAD17413.1) from *R. pseudosolanacearum* GMI1000, Bll8054 (GenBank: BAC53319.1) from *B. diazoefficiens* USDA 110 (JCM 10833, type strain), JI59_12175 (GenBank: AIT80485.1) from *N. pentaromativorans* (JCM 12182, type strain), RLO149_c030420 (GenBank: AEI94998.1) from *R. litoralis* (JCM 21268, type strain), SO_2107 (GenBank: AAN55154.2) from *S. oneidensis* (JCM 31522, type strain), PESP_a3768 (GenBank: ASM51527.1) and PESP_b0377 (GenBank: ASM51950.1) from *P. espejiana* (JCM 20705, type strain), MishRS11D_32800 (GenBank: BBL76182.1), MishRS11D_34730 (GenBank: BBL76375.1) and MishRS11D_37830 (GenBank: BBL76685.1) from *M. ishizawai* (JCM 18894, type strain). The genomic DNAs of *C. sphaeroides* and *R. pseudosolanacearum* GMI1000 were purchased from the Leibniz Institute DSMZ-German Collection of Microorganisms and Cell Cultures (Braunschweig, Germany) and American Type Culture Collection (VT, USA), respectively. The other genomic DNAs used in this study were purchased from Japan Collection of Microorganisms (Ibaraki, Japan). The forward primers were designed to eliminate *N*-terminal signal sequences predicted by the SignalP6.0 server (https://services.healthtech.dtu.dk/services/SignalP-6.0/)^19^. The amplified gene was inserted between the XhoI and NdeI sites of the pET30a vector amplified as described previously^4^ by the SLiCE method^20^ to add a *C*-terminal His_6_-tag to the target proteins. The constructed plasmids for production of the recombinant Rsp_3187 (CsOpgD2) and MishRS11D_32800 (MiOpgD2) were transformed into *E. coli* BL21 (DE3) cells and the transformants were cultured in 1 L Luria-Bertani medium containing 30 mg/L kanamycin at 37℃ until the absorbance at 600 nm reached 0.6. The constructed plasmids for production of the recombinant Rsp0262 (RpOpgD3), Bll8054 (BdOpgD), SO_2107 (SoOpgD2), PESP_a3768 (PeOpgD), PESP_b0377 (PeOpgD2), MishRS11D_34730 (MiOpgD3) and MishRS11D_37830 (MiOpgD4) were transformed into Rosetta2 (DE3) cells. The constructed plasmids for production of the recombinant RSc3057 (RpOpgD2), JI59_12175 (NpOpgD), RLO149_c030420 (RlOpgD) were transformed into BL21 Codon Plus (DE3). The transformants were cultured in 1 L Luria-Bertani medium containing 30 mg/L kanamycin and 20 mg/mL chloramphenicol at 37℃ until the absorbance at 600 nm reached 0.6. Expression of RpOpgD3, BdOpgD, SoOpgD2, PeOpgD, PeOpgD2, MiOpgD2 was induced by adding isopropyl β -D-1-thiogalactopyranoside to a final concentration of 0.1 mM, and cells were cultured at 15℃ for 24 h. For expression of CsOpgD2, RpOpgD2, MiOpgD3, MiOpgD4 and NpOpgD, induction performed at 20℃ (CsOpgD2), 28℃ (RpOpgD2), 30℃ (MiOpgD3, MiOpgD4), 37℃ (NpOpgD). The cells were centrifuged at 8800 g, 4℃ for 10 min and resuspended in 50 mM Tris-HCl buffer (pH 7.5). The resuspended cells were disrupted with sonication, and centrifuged at 27700 g, 4℃ for 10 min to obtain a cell extract. The cell extract was loaded onto a HisTrap^TM^ FF crude column (5 mL; Cytiva) pre-equilibrated with buffer (50 mM Tris-HCl, 500 mM NaCl, pH 7.5). After washing the column with a buffer containing 50 mM Tris-HCl, 500 mM NaCl and 10 mM imidazole, the target protein was eluted with linear gradient of 0–300 mM imidazole in a buffer containing 50 mM Tris-HCl and 500 mM NaCl (pH 7.5). Vivaspin Turbo 15 10,000 or 30,000 molecular weight cutoff centrifugal filter (Sartorius, Göttingen, Germany), 10,000 molecular weight cutoff centrifugal filter and Amicon Ultra 30,000 molecular weight cutoff centrifugal filter (Merck, NJ, USA) were used to concentrate a portion of the fractionated protein and exchange the buffer to 50 mM Tris-HCl (pH 7.5) buffer containing 50 mM NaCl. Each purified protein migrated as a single band around 50kDa of SDS-PAGE gels, which is consistent with the theoretical molecular masses of the target enzymes (Supplementary Table 4). Concentrations of the purified enzymes were calculated from the absorbance at 280 nm^21^.

### Preparation of β-1,2-glucans

Linear β-1,2-glucans with an average DPs of 121 calculated from their number average molecular weight^22^ were used for TLC analysis, ESI-MS, NMR and investigation of general properties, substrate specificities and specific activity. Linear β-1,2-glucans with an average DPs of 17.7 estimated from their number average molecular weight^22^ were used for crystallization to obtain the Michaelis complexes of RpOpgD3 (D384N), RlOpgD (D373N) and PeOpgD (D392N). Linear β-1,2-glucans with an average DP of 25 estimated from NMR analysis^23^ were used for crystallization to obtain the Michaelis complex of CsOpgD2 (D381N).

### TLC analysis

The GH186 enzymes were incubated with 1% linear β-1,2-glucan (average DPs of 121) under the conditions shown in Supplementary Table 5 until the substrates were fully consumed (sample 1). BtBGL (0.24 mg/mL as a final concentration) was added to each of sample 1 and then each reaction mixture was incubated at 37℃ for sufficient time to remove linear moieties of β-1,2-glucans completely (sample 2). BtBGL and CpSGL (0.24 mg/mL and 0.06 mg/mL as final concentrations, respectively) were added to each of sample 1 and then each reaction mixture was incubated at 37℃ for sufficient time to fully degrade the substrate and the products (sample 3). The reaction mixtures (1 μL) were spotted onto TLC Silica Gel 60 WF_254_S plates (Merck). Each plate was developed once with a solution containing an appropriate concentration of acetonitrile (Supplementary Table 6). The plates were then soaked in a 5% (w/v) sulfuric acid/methanol solution and heated in an oven until the spots were clearly visualized. Linear β-1,2-glucan with average DPs of 121, a mixture of β-1,2-glucooligosaccharides with DPs of 3–7 and cyclic-β-1,2-glucohexadecaose producted by XccOpgD were used as markers^4^.

### ESI-MS

Samples 1–3 (100 μL) were prepared under the same condition as in the case of TLC analysis (Supplementary Table 5). Amberlite MB4 (Organo) was added to all samples to remove ionic compounds. The resultant solutions were diluted 100-fold with a solvent (methanol/water = 1/1, v/v) containing 5 mM ammonium acetate. After filtration, the samples were loaded onto the Sciex X500 R QTOF (Sciex) in the positive mode at a 20 μL /min flow rate.

### NMR

Each of sample 1 (1 mL for RpOpgD3 and MiOpgD2; 2 ml for PeOpgD; 15 ml for CsOpgD2) was prepared under the same condition as in the case of TLC analysis (Supplementary Table 7). BtBGL and Bis-Tris HCl buffer (pH 5.5) (0.24 mg/mL and 35.5 mM as final concentration, respectively) were added to the sample and then the mixture was incubated at 37℃ for the same purpose as the TLC analysis. The BtBGL-resistant reaction products were purified by size-exclusion chromatography using a Toyopearl HW-40F column (∼2 L gel). The sample was eluted with distilled water after injection of the reaction mixtures (∼10 mL). The eluates were fractionated into 30-mL portions, and the fraction containing only a main product of the reaction was lyophilized. One-dimensional ^1^H NMR spectra of products were acquired in D_2_O at 298-333 K using 700 MHz NMR spectrometer Bruker AVANCE III HD (Bruker, MA, USA) or 800 MHz NMR Spectrometer Bruker AVANCE NEO (Bruker, MA, USA) with tBuOH (δ 1.23 ppm for ^1^H) as an internal standard. The H-1 chemical shift at the α-anomer was assigned by referring to the Karplus curve^24^.

### General properties

The optimum pHs of RpOpgD2, RpOpgD3, MiOpgD2, MiOpgD3 and MiOpgD4 were determined by incubating each protein (0.001, 0.005, 0.2, 0.01 and 1 mg/ml as final concentrations, respectively) in various 20 mM buffers (sodium acetate, pH 4.0–5.0; Tris-HCl, pH 6.0–8.0; glycine-NaOH, pH 9.0–10.0) containing 0.25% linear β-1,2-glucan (average DPs of 121) at 30℃ for 10 min. Enzymatic reactions were terminated by heating at 100℃ for 5 min to terminate reactions. In the case of CsOpgD2, BdOpgD, NpOpgD, RlOpgD, SoOpgD2, PeOpgD, PeOpgD2, final concentrations of the proteins (0.05, 2.0, 0.01, 1.0, 0.1, 0.25, and 1.0 mg/ml, respectively) and 20 min for the reaction time were adopted. The reaction was performed at 10℃ for PeOpgD2. Optimum temperatures of RpOpgD2 RpOpgD3, MiOpgD2, MiOpgD3, MiOpgD4 were determined by performing the reactions in 1% linear β-1,2-glucan (average DPs of 121) and 20 mM buffer selected based on analysis of optimal pH at various temperatures (0–70℃) for 10 min. Enzymatic reactions were terminated by heating at 100°C for 5 min to terminate the reaction. In the case of CsOpgD2, BdOpgD NpOpgD, RlOpgD, SoOpgD2, PeOpgD, PeOpgD2, 20 min was adopted for the reaction time. Final concentrations of the proteins in the reaction solutions were 0.001, 0.005, 0.2, 0.01, 1.0, 0.05, 2.0, 0.01, 0.01, 1.0, 0.25, 1.0 mg/ml, respectively. The reaction mixtures were analyzed by TLC analysis in the same way as described in the section of TLC analysis.

### Substrate specificity

CsOpgD2, RpOpgD2, RpOpgD3, BdOpgD, NpOpgD, RlOpgD, SoOpgD2, PeOpgD, PeOpgD2, MiOpgD2, MiOpgD3, MiOpgD4 were incubated containing each substrate (0.4% linear β-1,2-glucan average DPs of 121, 0.4% glucomannan, Neogen, MI, USA; 0.4% carboxymethyl cellulose, Merck; 0.4% soluble starch, FUJIFILM Wako Chemical Corporation, Osaka, Japan; 0.4% carboxymethyl curdlan, Neogen; 0.4% laminarin, Merck; 0.4% lichenan, Neogen; 0.4% barley β-glucan, Neogen; 0.4% tamarind-xyloglucan, Neogen; 0.4% arabinan, Neogen; 0.4% polygalacturonic acid, Neogen; 0.4% arabinogalactan, Neogen) at condition shown in Supplementary Table 8. The reaction patterns were analyzed by TLC using 72% (v/v) acetonitrile for CsOpgD2, PeOpgD2 and 65% (v/v) acetonitrile for RpOpgD2, RpOpgD3, NpOpgD, RlOpgD, SoOpgD2, PeOpgD, MiOpgD2, MiOpgD3, MiOpgD4, as developers.

### Specific activity

Specific activity of CsOpgD2, RpOpgD3 and RlOpgD for linear β-1,2-glucans were determined by performing reactions under condition shown in Supplementary Table 9. The reactions were stopped by heat treatment at 100°C for 5 min. BtBGL (1.2 mg/mL as a final concentration) was added to degrade linear moieties in β-1,2-glucans into glucose. pH was adjusted around pH 5.5 with 20 mM of bis-Tris-HCl (pH 5.5), except for the method for RlOpgD. The reaction products were reduced using a one-fifth volume of 1 M NaBH_4_. The same volume of 1 M acetate as that of the 1M NaBH_4_ solution was added to each sample to neutralize each NaBH_4_-treated solution. The samples were then treated with 0.6 mg/mL of BtBGL and 0.25 mg/mL CpSGL (as final concentrations) at 30°C until the BtBGL-resistant cyclic β-1,2-glucans were degraded into glucose and non-degradable glucopentaose completely. The main residual oligosaccharide was identified to be glucopentaose by ESI-MS. Color development of the reaction mixtures was performed using the GOPOD method^25^ to quantify the concentration of glucose derived from the products released by the final treatment with BtBGL and CpSGL. Molarities of products were determined by a standard curve of glucose. Specific activities of the enzymes (as activities of releasing cyclic glucans) were calculated based on DPs of the main products. One CβGα molecule without side chain is regarded to release (DP−5) glucose molecules by treatment with BtBGL and CpSGL. Each analysis was performed in triplicate.

### Crystallography

As described above, CsOpgD2 (D381N), RpOpgD3 (D384N), RlOpgD (D373N) and PeOpgD (D392N) were purified using a HisTrap^TM^ FF crude 5 mL column. Crystals of the mutant enzymes for linear β-1,2-glucans (average DPs of 17.7) complexes were obtained under the conditions shown in Supplementary Table 10. Crystals of the mutants were soaked in the cryoprotectant solution shown in Supplementary Table 10. The crystals were kept at 100 K in a nitrogen-gas stream during data collection. Data collection was performed at a beamline (BL-5A) in Photon Factory (Tsukuba, Japan). The X-ray diffraction data of crystals for complexes with linear β-1,2-glucans were collected at 1.0 Å and processed with X-ray Detector Software (http://xds.mpimf-heidelberg.mpg.de/)^26^ and the AIMLESS program (http://www.ccp4.ac.uk/). The initial phases of the complex structures were determined by molecular replacement using the structures predicted by AlphaFold Server (https://golgi.sandbox.google.com/)^27^ as model structures. Molecular replacement, auto model building were performed using MOLREP^28^, Buccaneer^29^ and ARP/wARP^30^ programs, respectively (http://www.ccp4.ac.uk/). Automated and manual refinement was performed using REFMAC5^31^ and Coot^32^, respectively. The final data was validated by wwPDB Validation System (https://validate-rcsb-1.wwpdb.org/). The structures were visualized, and the figures were prepared using PyMOL (https://www.pymol.org/).

### Sequence similarity network

Sequence Similarity Network (SSN) was constructed using the sequence BLAST option of the Enzyme Function Initiative – Enzyme Similarity Tool (EFI-EST; https://efi.igb.illinois.edu/efi-est/)^33^. The amino acid sequence of XccOpgD was used as a query. The UniProt BLAST search parameter was set to an E-value of 1, with the maximum number of retrieved sequences limited to 10,000, in order to exclude overly distantly related sequences that would be difficult to interpret the relations of catalytic functions. For SSN edge calculation, the E-value threshold was also set to 1. Based on parameter optimization, an alignment score threshold of 140 was chosen, as this setting resulted in the formation of distinct clusters. To exclude partial sequences as much as possible, the minimum sequence length was set to 201 amino acids. The resulting 95% identity representative node (RepNode) network, composed of 5,184 nodes and 428,369 edges, was visualized and manually edited using Cytoscape version 3.10.3 (https://cytoscape.org/).

### Sequence analysis

All sequence alignment was performed using ClustalW in MEGA11^34^ and visualized using the ESPript 3.0 server (http://espript.ibcp.fr/ESPript/ESPript/)^35^.

### Construction of *RSp0262*-Deletion and -Complemented Mutants

The merged up- and downstream region (400 + 400 bp, containing BamHI- and HindIII-digestion sites) of the RSp0262 gene was synthesized by Eurofins Genomics and cloned into pK18mobsacB to create pΔRSp0262. This plasmid was electroporated into Ter331 competent cells (OD_600_ = 0.6, in 10% glycerol), and kanamycin-resistant (50 μg/mL) recombinants were selected. The recombinants were incubated in SOC medium (1 mL, Sigma-Aldrich) for 6–8 hours. Subsequently, kanamycin-sensitive (50 μg/mL; first selection) and sucrose-resistant (10%; second selection) recombinants were isolated. The deletion of the target region was confirmed by PCR with KOD One PCR Master Mix (Toyobo). The gene complementation was carried out using the modified pDSK519 vector harboring RSp0262. The selection of transformants was performed on agar containing 50 μg/mL kanamycin and checked by PCR.

Detection of CβG13α from *R. pseudosolanacearum* GMI1000, *R. pseudosolanacearum* cells grown in B medium (100 mL) for 5 h were collected by centrifugation at 10,000 × *g* for 5 min and washed with Milli-Q water (20 mL). The cell pellets were suspended in 20% acetone and sonicated to extract CβG13α. The supernatants were collected by centrifugation at 8,000 × *g* for 10 min and subjected to LC/MS analysis. The following LC/MS conditions were used: InertSustain C18 column (150 × 2.1 mm, 3 μm, GL Sciences), eluent: 10%–35% MeCN in 0.1% formic acid in water over 30 min, column oven: 40°C, flow rate: 200 μL/min, injection volume: 5 μL, detection: ESI positive.

### Virulence Assay with Tomato Seedlings

Tomato seeds were presoaked in water for 2 days. Then, the seeds were spread on wet tissue paper in a glass tray and allowed to germinate in a growth chamber maintained at 28°C with a 12-h photoperiod. Water was sprinkled regularly to sustain the germination process for 7 days. Roots of each seedling were dipped in the bacterial inoculum (OD_600_ = 1.0) followed by transfer of the seedling to an empty 1.5-mL microfuge tube. The root-dip-inoculated seedlings transferred to microfuge tubes were exposed to air for 5 min prior to the addition of 1 to 1.5 mL of water to each tube. All of the inoculated seedlings along with controls were transferred to a growth chamber maintained at 28°C with a 12-h photoperiod. Seedlings were analyzed for disease progression from the next day onward till the seventh day postinoculation, and observations were recorded.

### Virulence Assay with Tomato Plants

Tomato plants (cv. Oogata-Fukuju) were grown in pots containing commercial soil (Takii Seed) and watered with five-fold-diluted Hoagland’s solution in a growth room at 25°C under 10000 lx for 16 h per day. The roots of 5-week-old tomato plants were soaked in bacterial suspension at 1.0×10^8^ CFU/mL for 30 min and then washed in running water. The inoculated plants were grown in water-culture pots (Yamato Plastic) with five-fold-diluted Hoagland’s solution in a growth room at 25°C under 10000 lx for 16 h per day. Within each trial, three plants were used. Plants were coded and inspected for wilting symptoms daily after inoculation. Plants were rated on a 0–5 disease index scale: 0, no wilting; 1, 1%–25% wilting; 2, 26%–50% wilting; 3, 51%–75% wilting; 4, 76%–99% wilting; 5, dead.

## Acknowledgement

This work was supported by Photon Factory for X-ray data collection (Proposal No. 2022G523). This work was supported by JSPS KAKENHI Grant Number JP24KJ2021. We express our gratitude to Dr. Shinya Fushinobu for his valuable advice.

## References

1. Bohin, J. Cell-associated glucans of Burkholderia solanacearum and Xanthomonas campestris pv. citri: a new family of periplasmic glucans. J Bacteriol 178, 2263–2271 (1996).

2. Talaga, P. et al. Osmoregulated periplasmic glucans of the free-living photosynthetic bacterium Rhodobacter sphaeroides. Eur. J. Biochem. 269, 2464–2472 (2002).

3. Rigano, L. A. et al. Bacterial cyclic β-(1,2)-glucan acts in systemic suppression of plant immune responses. Plant Cell 19, 2077–2089 (2007).

4. Motouchi, S., Komba, S., Nakai, H. & Nakajima, M. Discovery of Anomer-Inverting Transglycosylase: Cyclic Glucohexadecaose-Producing Enzyme from *Xanthomonas*, a Phytopathogen. J. Am. Chem. Soc. 146, 17738–17746 (2024).

5. Timilsina, S. et al. Xanthomonas diversity, virulence and plant–pathogen interactions. Nat. Rev. Microbiol. 18, 415–427 (2020).

6. Ishiguro, R. et al. Function and structure relationships of a β-1,2-glucooligosaccharide-degrading β-glucosidase. FEBS Lett. 591, 3926–3936 (2017).

7. Abe, K. et al. Biochemical and structural analyses of a bacterial endo-β-1,2-glucanase reveal a new glycoside hydrolase family. J Biol Chem 292, 7487–7506 (2017).

8. Laitinen, T., Rouvinen, J. & Peräkylä, M. MM-PBSA free energy analysis of endo-1,4-xylanase II (XynII)–substrate complexes: binding of the reactive sugar in a skew boat and chair conformation. Org. Biomol. Chem. 1, 3535–3540 (2003).

9. Perić-Hassler, L., Hansen, H. S., Baron, R. & Hünenberger, P. H. Conformational properties of glucose-based disaccharides investigated using molecular dynamics simulations with local elevation umbrella sampling. Carbohydr. Res. 345, 1781–1801 (2010).

10. Abramson, J. et al. Accurate structure prediction of biomolecular interactions with AlphaFold 3. Nature 630, 493–500 (2024).

11. Kai, K. et al. Methyl 3-Hydroxymyristate, a Diffusible Signal Mediating *phc* Quorum Sensing in *Ralstonia solanacearum*. ChemBioChem 16, 2309–2318 (2015).

12. Ujita, Y., Sakata, M., Yoshihara, A., Hikichi, Y. & Kai, K. Signal Production and Response Specificity in the *phc* Quorum Sensing Systems of *Ralstonia solanacearum* Species Complex. ACS Chem. Biol. acschembio.9b00553 (2019) doi:10.1021/acschembio.9b00553.

13. Kim, Y.-K., Kitaoka, M., Hayashi, K., Kim, C.-H. & Côté, G. L. A synergistic reaction mechanism of a cycloalternan-forming enzyme and a d-glucosyltransferase for the production of cycloalternan in Bacillus sp. NRRL B-21195. Carbohydr. Res. 338, 2213–2220 (2003).

14. Terada, Y. et al. Comparative Study of the Cyclization Reactions of Three Bacterial Cyclomaltodextrin Glucanotransferases. Appl. Environ. Microbiol. 67, 1453–1460 (2001).

15. Wang, Y. et al. Exploring the enzymatic landscape of 4-α-glucanotransferases in carbohydrate bioprocessing. Biotechnol. Adv. 86, 108737 (2026).

16. Guidolin, L. S., Ciocchini, A. E., De Iannino, N. I. & Ugalde, R. A. Functional mapping of Brucella abortus cyclic β-1,2-glucan synthase: Identification of the protein domain required for cyclization. J. Bacteriol. 191, 1230–1238 (2009).

17. Tanaka, N. et al. Functional and structural analysis of a cyclization domain in a cyclic β-1,2-glucan synthase. Appl. Microbiol. Biotechnol. 108, 187 (2024).

18. Bontemps-Gallo, S., Bohin, J. P. & Lacroix, J. M. Osmoregulated periplasmic glucans. EcoSal Plus 7, 1–17 (2017).

19. Teufel, F. et al. SignalP 6.0 predicts all five types of signal peptides using protein language models. Nat. Biotechnol. 40, 1023–1025 (2022).

20. Motohashi, K. A simple and efficient seamless DNA cloning method using SLiCE from Escherichia coli laboratory strains and its application to SLiP site-directed mutagenesis. BMC Biotechnol. 15, 47 (2015).

21. Pace, C. N., Vajdos, F., Fee, L., Grimsley, G. & Gray, T. How to measure and predict the molar absorption coefficient of a protein. Protein Sci 4, 2411–2423 (1995).

22. Nakajima, M. et al. Enzymatic control and evaluation of degrees of polymerization of β-(1→2)-glucans. Anal. Biochem. 632, 114366 (2021).

23. Nakajima, M. et al. 1,2-β-oligoglucan phosphorylase from Listeria innocua. PLoS One 9, e92353–e92353 (2014).

24. Minch, M. J. Orientational dependence of vicinal proton-proton NMR coupling constants: The Karplus relationship. Concepts Magn. Reson. 6, 41–56 (1994).

25. Kobayashi, K. et al. Characterization and structural analyses of a novel glycosyltransferase acting on the β-1,2-glucosidic linkages. J Biol Chem 298, 101606 (2022).

26. Kabsch, W. XDS. Acta Crystallogr D Biol Crystallogr D66, 125–132 (2010).

27. Jumper, J. et al. Highly accurate protein structure prediction with AlphaFold. Nature 596, 583–589 (2021).

28. Vagin, A. & Teplyakov, A. Molecular replacement with *MOLREP*. Acta Crystallogr. D Biol. Crystallogr. 66, 22–25 (2010).

29. Cowtan, K. The *Buccaneer* software for automated model building. 1. Tracing protein chains. Acta Crystallogr. D Biol. Crystallogr. 62, 1002–1011 (2006).

30. Langer, G., Cohen, S. X., Lamzin, V. S. & Perrakis, A. Automated macromolecular model building for X-ray crystallography using ARP/wARP version 7. Nat. Protoc. 3, 1171–1179 (2008).

31. Murshudov, G. N., Vagin, A. A. & Dodson, E. J. Refinement of Macromolecular Structures by the Maximum-Likelihood Method. Acta Crystallogr. D Biol. Crystallogr. 53, 240–255 (1997).

32. Emsley, P. & Cowtan, K. *Coot*: model-building tools for molecular graphics. Acta Crystallogr. D Biol. Crystallogr. 60, 2126–2132 (2004).

33. Oberg, N., Zallot, R. & Gerlt, J. A. EFI-EST, EFI-GNT, and EFI-CGFP: Enzyme Function Initiative (EFI) Web Resource for Genomic Enzymology Tools. J. Mol. Biol. 435, 168018 (2023).

34. Tamura, K., Stecher, G. & Kumar, S. MEGA11: Molecular Evolutionary Genetics Analysis Version 11. Mol. Biol. Evol. 38, (2021).

35. Robert, X. & Gouet, P. Deciphering key features in protein structures with the new ENDscript server. Nucleic Acids Res. 42, 320–324 (2014).

