## Supplementary Figures 1-21, Supplementary Tables 1-10, Supplementary Note 1. for "Discovery of a wide variety of α-1,6-cyclized β-1,2-glucan synthases: a new entrance for host-microbe interactions"

Supplementary Figures 1-21
Supplementary Tables 1-10
Supplementary Note 1

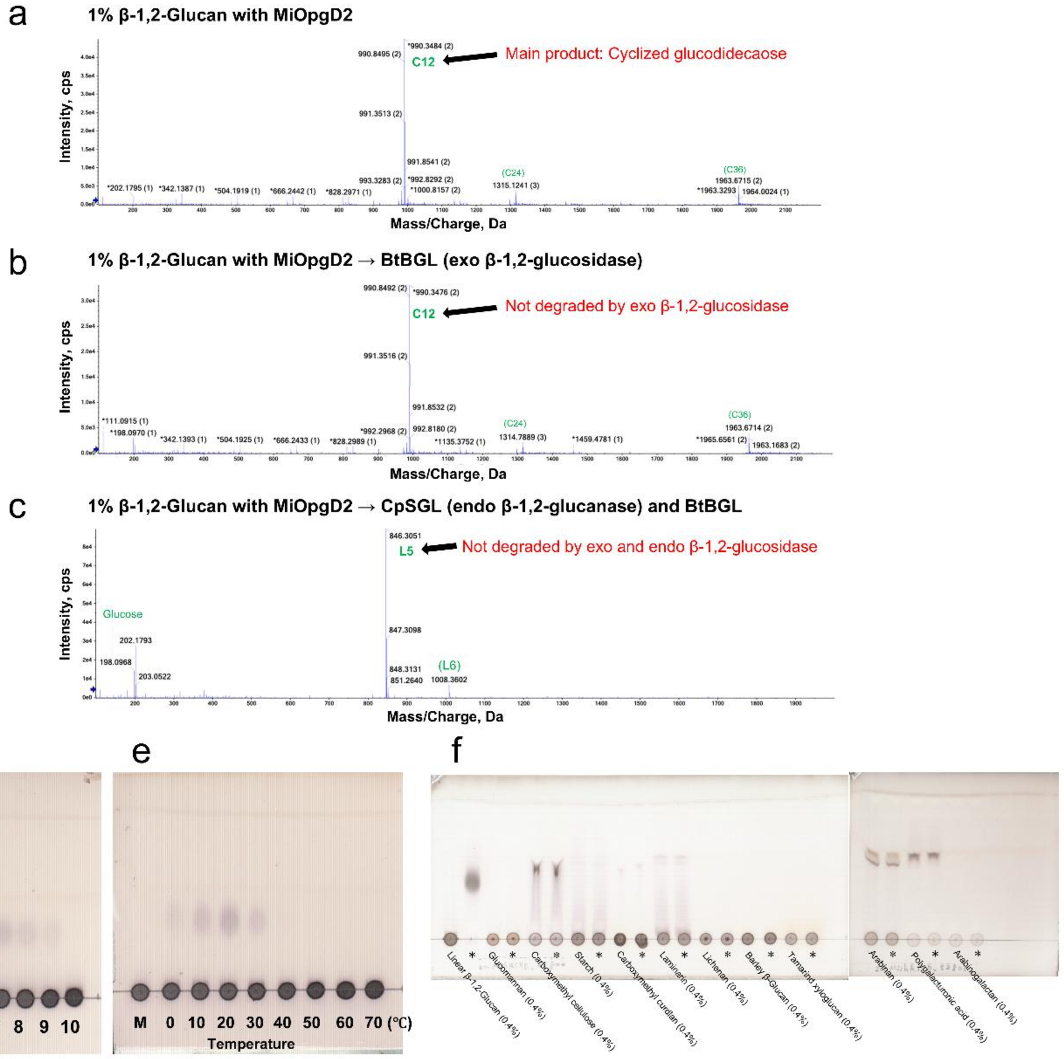

a
1% β-1,2-Glucan with MiOpgD2
Main product: Cyclized glucodidecaose
C12
(C24)
(C36)
Intensity, cps
Mass/Charge, Da
b
1% β-1,2-Glucan with MiOpgD2 → BtBGL (exo β-1,2-glucosidase)
Not degraded by exo β-1,2-glucosidase
C12
(C24)
(C36)
Intensity, cps
Mass/Charge, Da
c
1% β-1,2-Glucan with MiOpgD2 → CpSGL (endo β-1,2-glucanase) and BtBGL
Not degraded by exo and endo β-1,2-glucosidase
L5
(L6)
Glucose
Intensity, cps
Mass/Charge, Da
d
M 4 5 6 7 8 9 10
pH
e
M 0 10 20 30 40 50 60 70 (℃)
Temperature
f
Linear β-1,2-Glucan (0.4%)
Glucomannan (0.4%)
Carboxymethyl cellulose (0.4%)
Starch (0.4%)
Carboxymethyl curdlan (0.4%)
Laminarin (0.4%)
Lichenan (0.4%)
Barley β-Glucan (0.4%)
Tamarind xyloglucan (0.4%)
Arabinan (0.4%)
Polygalacturonic acid (0.4%)
Arabinogalactan (0.4%)

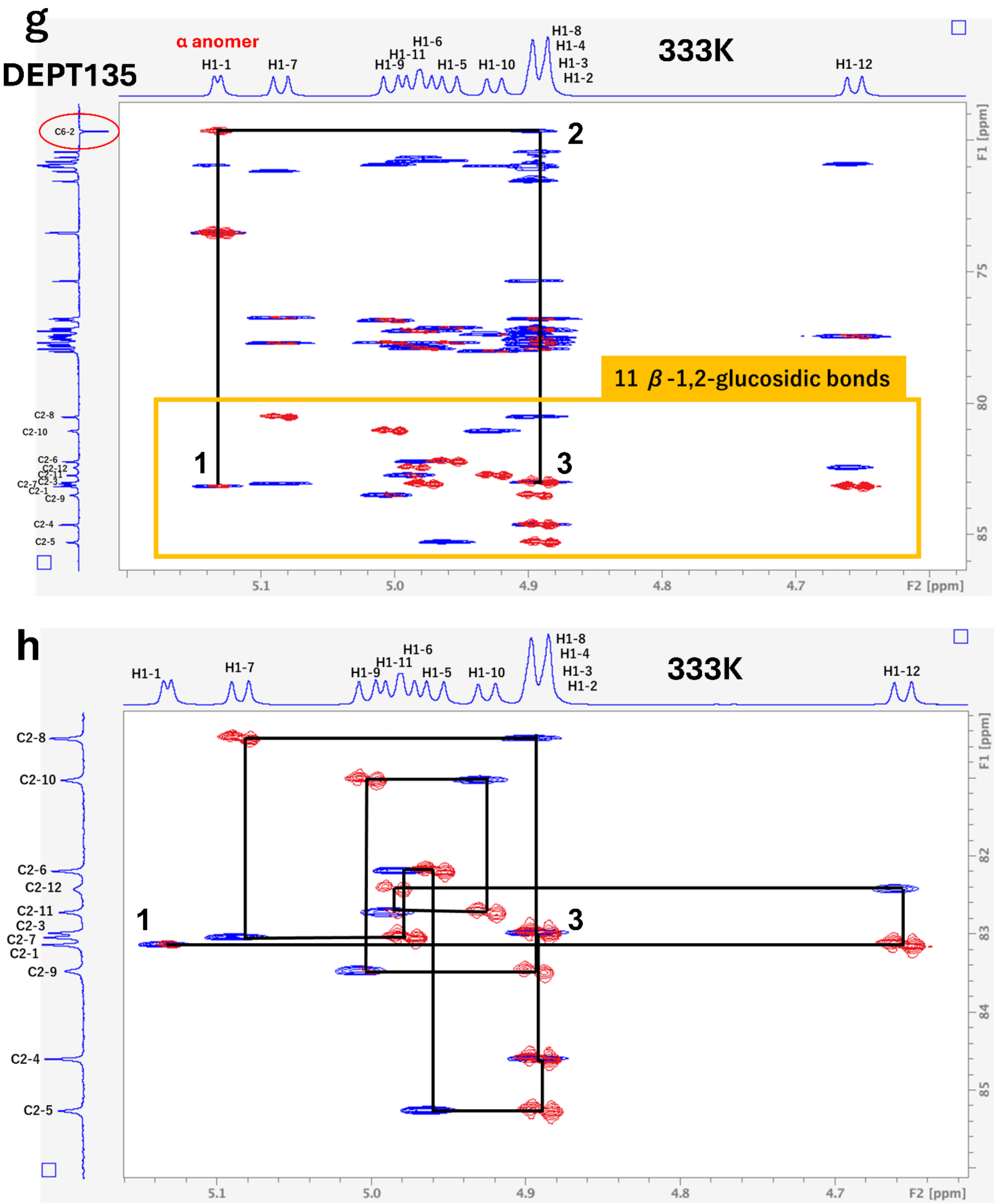


**Supplementary Figure 1. Analysis of main products released by MiOpgD2, optimal condition for catalytic reaction and substrate specificity.**

(a–c) Electrospray ionization-mass spectrometry (ESI-MS) analysis of the reaction products. The peaks are assigned as $[M + nNH_4]^{n+}$, and the arrows indicate the main products. Green letters and numbers represent forms of compounds (cyclic or linear) and DPs of products, respectively. For example, C16 represents cyclized hexadecaose. Peak labels are shown only for signals above the *m/z* threshold indicated by the arrows along the vertical axes. (a) Reaction products released from linear β-1,2-glucan by MiOpgD2. (b) Reaction products after BtBGL treatment. (c) Reaction products after BtBGL and CpSGL treatment. (d) TLC analysis for investigating optimal pH. Lane M, linear β-1,2-glucans (average DP of 121, see Methods for details) as a control; the other lanes, the reaction solutions of MiOpgD2 with linear β-1,2-glucans (average DP of 121) incubated at respective pHs. (e) TLC analysis for investigating optimal temperature. Lane M, linear β-1,2-glucans (average DP of 121) as a control; the other lanes, the reaction solutions of MiOpgD2 with linear β-1,2-glucans (average DP of 121) incubated at respective temperatures. (f) Substrate specificity of MiOpgD2. The reaction was performed at 30°C. Asterisks indicate that the reaction time was 24 h. The other lanes represent a reaction time of 0 h. (g, h) 700 MHz NMR analysis of the purified main product released by MiOpgD2. Blue and red peaks represent HSQC-TOCSY and HMBC data, respectively. The H1 and C2 atoms of α-anomeric Glc moiety are numbered as 1 (i.e. H1-1 and C2-1, respectively). The remaining Glc moieties are numbered sequentially toward the reducing end. (g) The trace of correlations around α-1,6-glucosidic bonds. Black lines trace correlations from H1-3/C2-3 to H1-1/C2-1 in the direction of the non-reducing end. (h) The trace of correlations derived from β-1,2-glucosidic bonds. Black lines trace correlations from H1-1/C2-1 to H1-3/C2-3 in the same direction as (g).

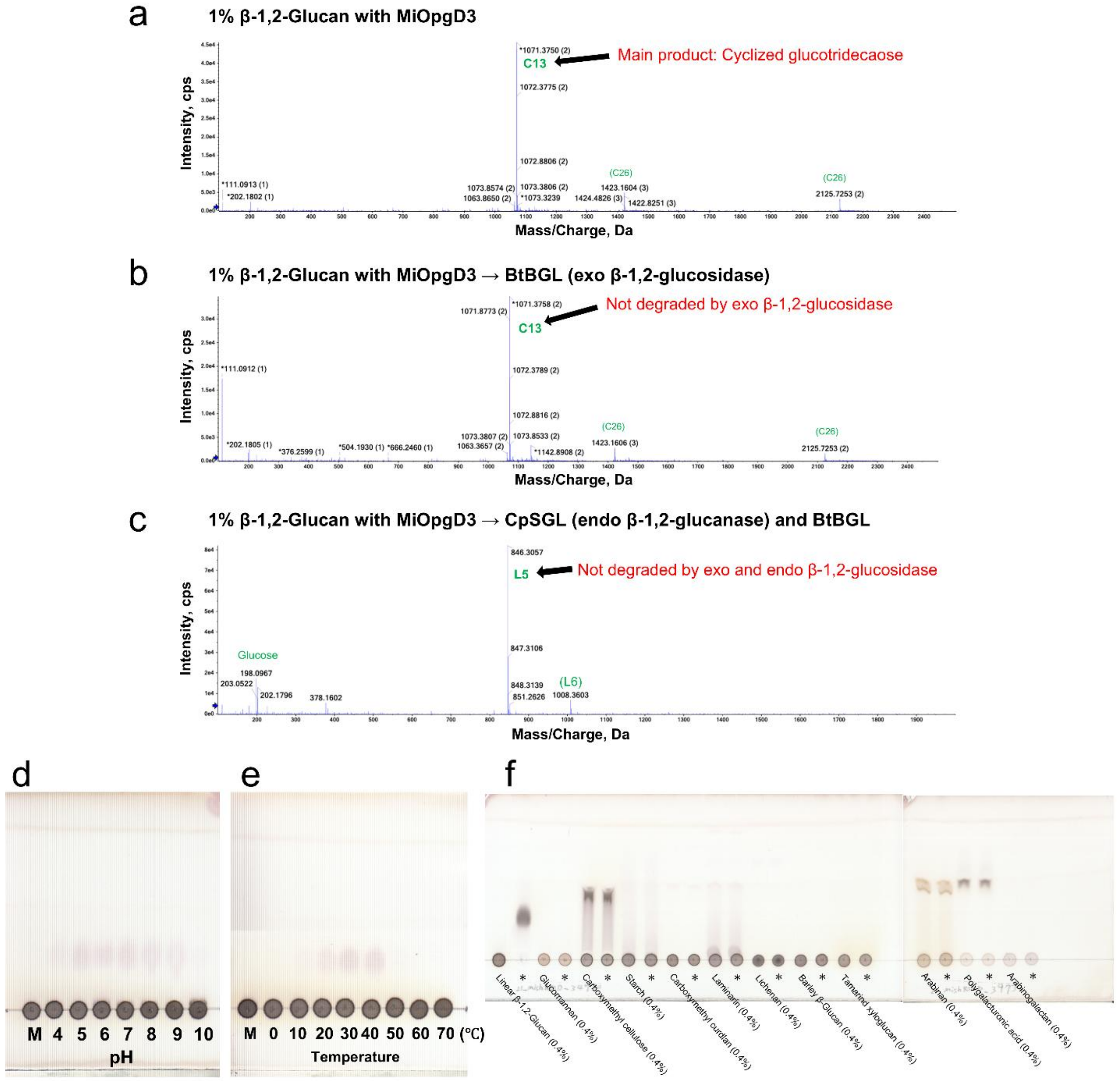


**Supplementary Figure 2. Analysis of main products released by MiOpgD3, optimal condition for catalytic reaction and substrate specificity.** (a–c) ESI-MS analysis of the reaction products. The peaks are assigned as [M + $nNH_4]^{n+}$, and the arrows indicate the main products. The *m/z* values and peak labels are presented in the same manner as in supplementary Fig. 1. (a) Reaction products released from linear β-1,2-glucan by MiOpgD3. (b) Reaction products after BtBGL treatment. (c) Reaction products after BtBGL and CpSGL treatment. (d) TLC analysis for investigating optimal pH. Lane M, linear β-1,2-glucans (average DP of 121) as a control; the other lanes, the reaction solutions of MiOpgD3 with linear β-1,2-glucans (average DP of 121) incubated at respective pHs. (e) TLC analysis for investigating optimal temperature. Lane M, linear β-1,2-glucans (average DP 121) as a control; the other lanes, the reaction solutions of MiOpgD3 with linear β-1,2-glucans (average DP 121) incubated at respective temperatures. (f) Substrate specificity of MiOpgD3. The reaction was performed at 30°C. Asterisks indicate that the reaction time was 24 h. The other lanes represent a reaction time of 0 h.

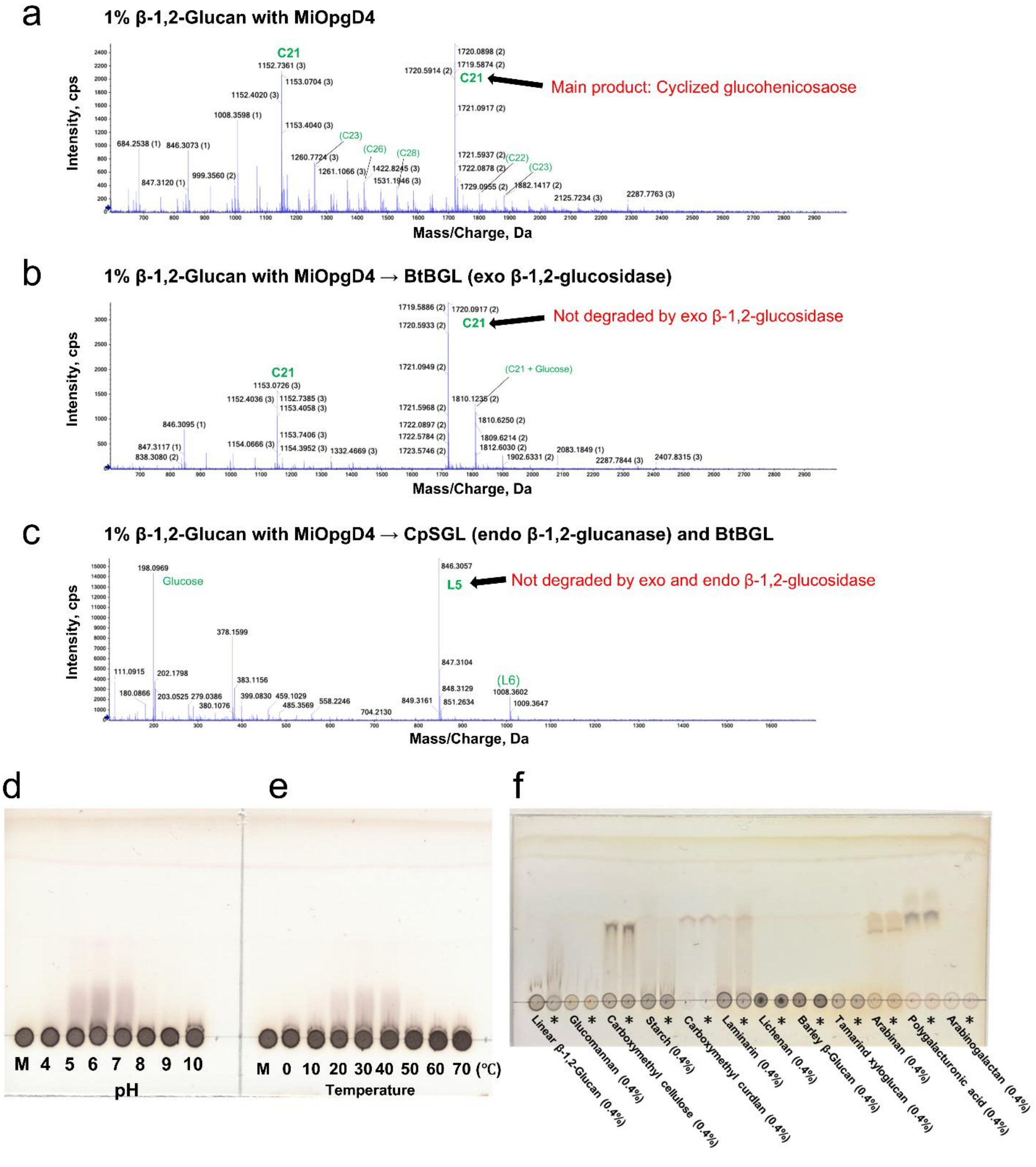


**Supplementary Figure 3. Analysis of main products released by MiOpgD4, optimal condition for catalytic reaction and substrate specificity.** (a–c) ESI-MS analysis of the reaction products. The peaks are assigned as $[M + nNH_4]^{n+}$, and the arrows indicate the main products. The *m/z* values and peak labels are presented in the same manner as in supplementary Fig. 1. (a) Reaction products released from linear β-1,2-glucan by MiOpgD4. (b) Reaction products after BtBGL treatment. (c) Reaction products after BtBGL and CpSGL treatment. (d) TLC analysis for investigating optimal pH. Lane M, linear β-1,2-glucans (average DP 121) as a control; the other lanes, the reaction solutions of MiOpgD4 with linear β-1,2-glucans (average DP 121) incubated at respective pHs. (e) TLC analysis for investigating optimal temperature. Lane M, linear β-1,2-glucans (average DP 121) as a control; the other lanes, the reaction solutions of MiOpgD4 with linear β-1,2-glucans (average DP 121) incubated at respective temperatures. (f) Substrate specificity of MiOpgD4. The reaction was performed at 30°C. Asterisks indicate that the reaction time was 24 h. The other lanes represent a reaction time of 0 h.

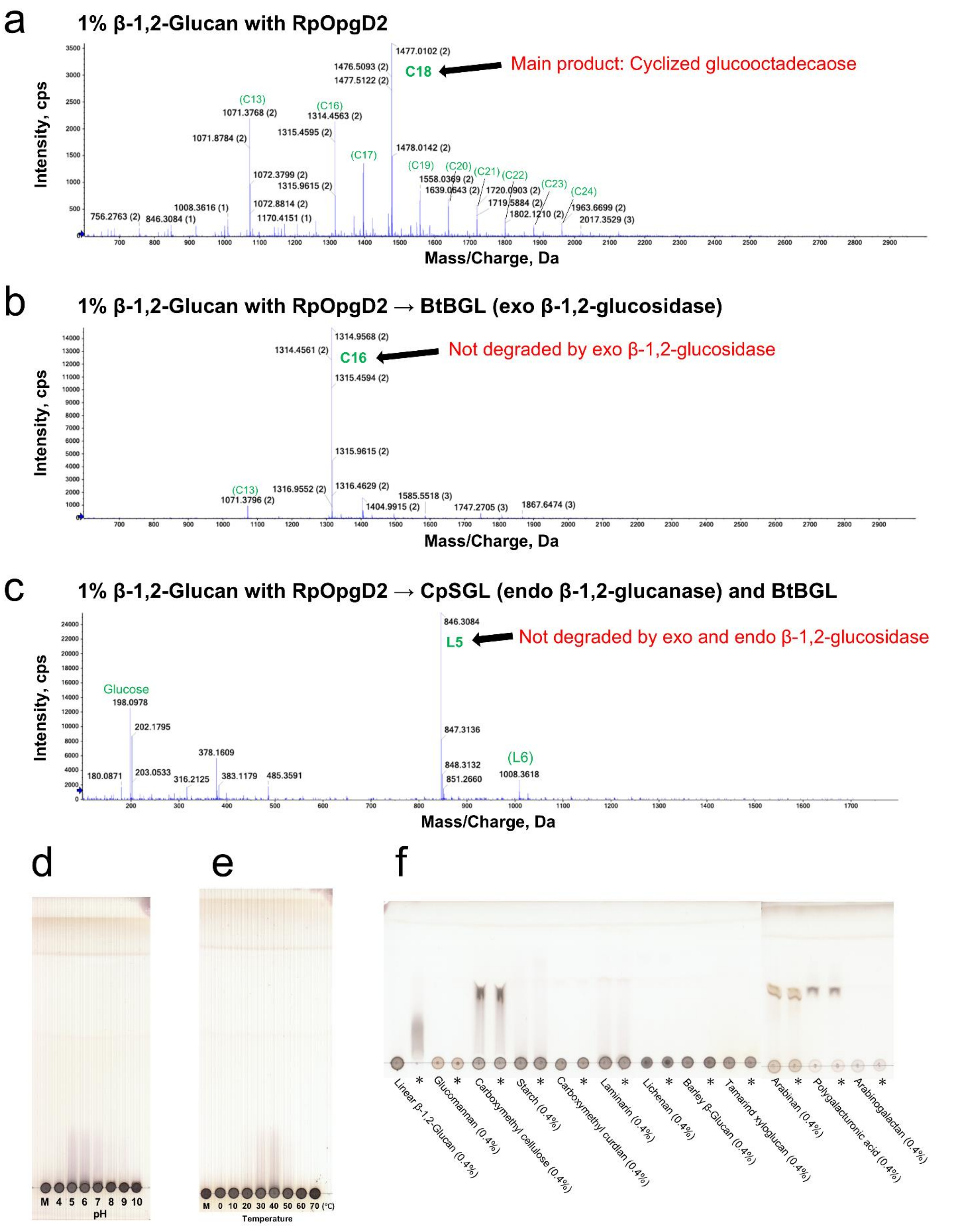


**Supplementary Figure 4. Analysis of main products released by RpOpgD2, optimal condition for catalytic reaction and substrate specificity.** (a–c) ESI-MS analysis of the reaction products. The peaks are assigned as $[M + nNH_4]^{n+}$, and the arrows indicate the main products. The *m/z* values and peak labels are presented in the same manner as in supplementary Fig. 1. (a) Reaction products released from linear β-1,2-glucan by RpOpgD2. (b) Reaction products after BtBGL treatment. (c) Reaction products after BtBGL and CpSGL treatment. (d) TLC analysis for investigating optimal pH. Lane M, linear β-1,2-glucans (average DP 121) as a control; the other lanes, the reaction solutions of RpOpgD2 with linear β-1,2-glucans (average DP 121) incubated at respective pHs. (e) TLC analysis for investigating optimal temperature. Lane M, linear β-1,2-glucans (average DP 121) as a control; the other lanes, the reaction solutions of RpOpgD2 with linear β-1,2-glucans (average DP 121) incubated at respective temperatures. (f) Substrate specificity of RpOpgD2. The reaction was performed at 30°C. Asterisks indicate that the reaction time was 24 h. The other lanes represent a reaction time of 0 h.

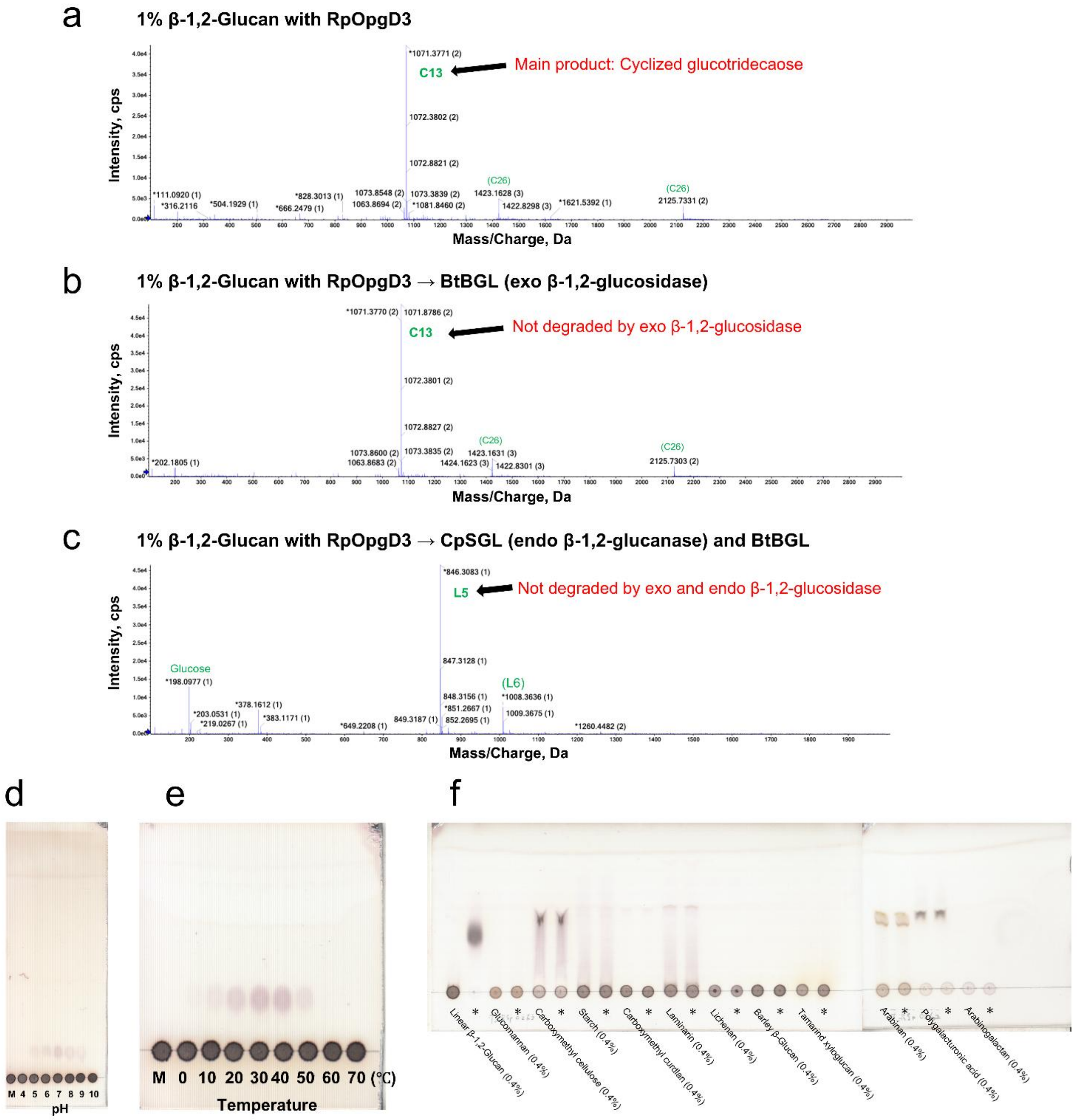
a
1% β-1,2-Glucan with RpOpgD3
Main product: Cyclized glucotridecaose
C13
Intensity, cps
Mass/Charge, Da
b
1% β-1,2-Glucan with RpOpgD3 → BtBGL (exo β-1,2-glucosidase)
Not degraded by exo β-1,2-glucosidase
C13
Intensity, cps
Mass/Charge, Da
c
1% β-1,2-Glucan with RpOpgD3 → CpSGL (endo β-1,2-glucanase) and BtBGL
Not degraded by exo and endo β-1,2-glucosidase
L5
Glucose
(L6)
Intensity, cps
Mass/Charge, Da
d
M 4 5 6 7 8 9 10
pH
e
M 0 10 20 30 40 50 60 70 (℃)
Temperature
f
Linear β-1,2-Glucan (0.4%)
Glucomannan (0.4%)
Carboxymethyl cellulose (0.4%)
Starch (0.4%)
Carboxymethyl curdlan (0.4%)
Laminarin (0.4%)
Lichenan (0.4%)
Barley β-Glucan (0.4%)
Tamarind xyloglucan (0.4%)
Arabinan (0.4%)
Polygalacturonic acid (0.4%)
Arabinogalactan (0.4%)

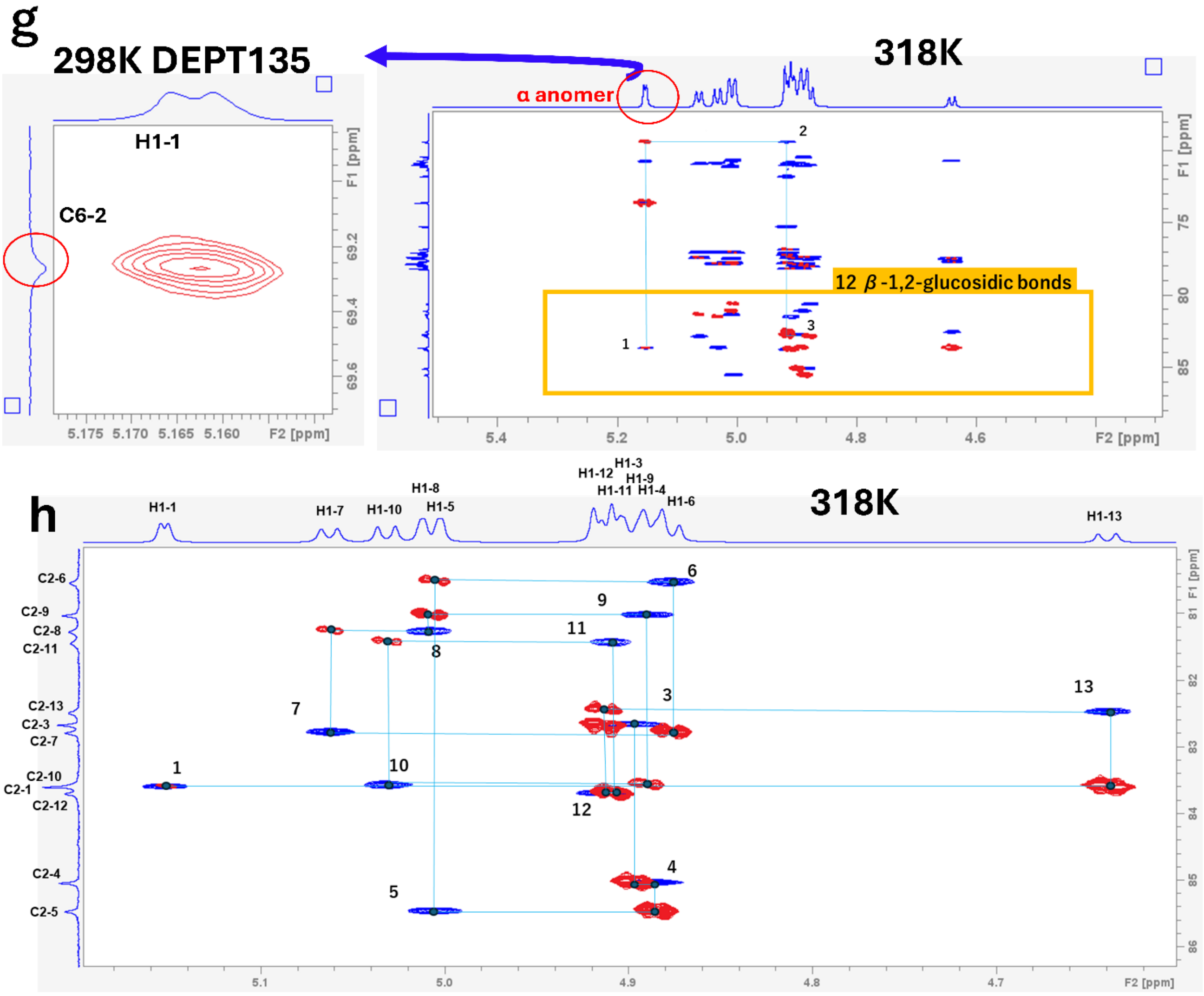


**Supplementary Figure 5. Analysis of main products released by RpOpgD3, optimal condition for catalytic reaction and substrate specificity.** (a–c) ESI-MS analysis of the reaction products. The peaks are assigned as $[M + nNH_4]^{n+}$, and the arrows indicate the main products. The *m/z* values and peak labels are presented in the same manner as in supplementary Fig. 1. (a) Reaction products released from linear β-1,2-glucan by RpOpgD3. (b) Reaction products after BtBGL treatment. (c) Reaction products after BtBGL and CpSGL treatment. (d) TLC analysis for investigating optimal pH. Lane M, linear β-1,2-glucans (average DP 121) as a control; the other lanes, the reaction solutions of RpOpgD3 with linear β-1,2-glucans (average DP 121) incubated at respective pHs. (e) TLC analysis for investigating optimal temperature. Lane M, linear β-1,2-glucans (average DP 121) as a control; the other lanes, the reaction solutions of RpOpgD3 with linear β-1,2-glucans (average DP 121) incubated at respective temperatures. (f) Substrate specificity of RpOpgD3. The reaction was performed at 30°C. Asterisks indicate that the reaction time was 24 h. The other lanes represent a reaction time of 0 h. (g, h) NMR analysis of the purified main product released by RpOpgD3. Blue and red peaks represent HSQC-TOCSY and HMBC data, respectively. The H1 and C2 atoms are numbered in the same manner as in supplementary Fig. 1. (g) The trace of correlations around the α-1,6-glucosidic bond. Cyan lines trace correlations from H1-3/C2-3 to H1-1/C2-1 in the direction of the non-reducing end (right,

NMR analysis at 318 K). A correlation indicating the α-1,6-glucosidic bond was demonstrated by 700 MHz NMR analysis with DEPT135 at 298 K (left). (h) The trace of correlations derived from β-1,2-glucosidic bonds (800 MHz NMR at 318 K). Cyan lines trace correlations from H1-1/C2-1 to H1-3/C2-3 in the same direction as (g).

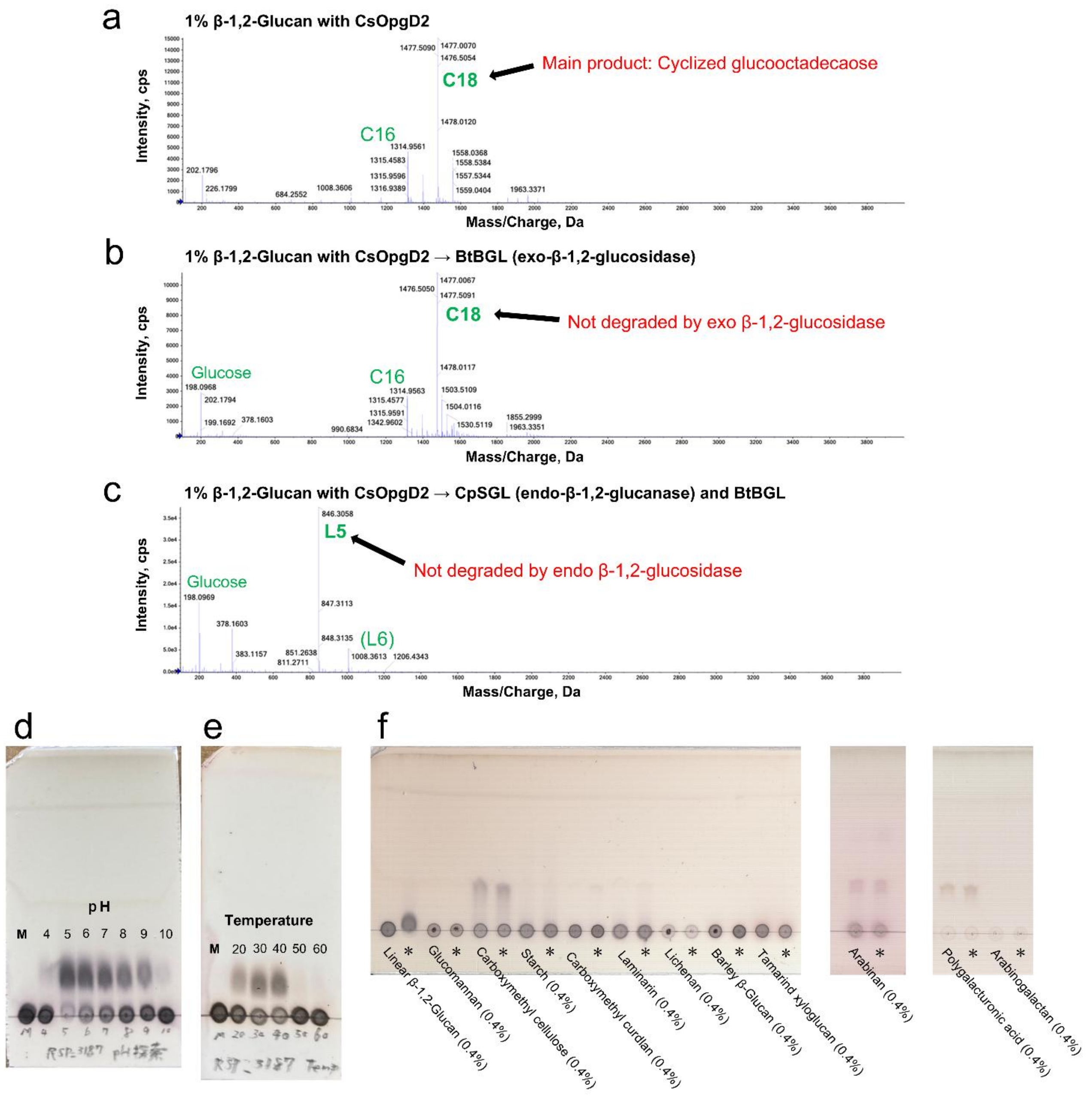

a
1% β-1,2-Glucan with CsOpgD2
Intensity, cps
Mass/Charge, Da
Main product: Cyclized glucooctadecaose
C18
C16
1477.5090
1477.0070
1476.5054
1478.0120
1314.9561
1315.4583
1315.9596
1316.9389
1558.0368
1558.5384
1557.5344
1559.0404
202.1796
226.1799
684.2552
1008.3606
1963.3371
b
1% β-1,2-Glucan with CsOpgD2 → BtBGL (exo-β-1,2-glucosidase)
Intensity, cps
Mass/Charge, Da
Not degraded by exo β-1,2-glucosidase
C18
C16
Glucose
1477.0067
1476.5050
1477.5091
1478.0117
1503.5109
1504.0116
1530.5119
1855.2999
1963.3351
1314.9563
1315.4577
1315.9591
1342.9602
990.6834
198.0968
202.1794
199.1692
378.1603
c
1% β-1,2-Glucan with CsOpgD2 → CpSGL (endo-β-1,2-glucanase) and BtBGL
Intensity, cps
Mass/Charge, Da
L5
Not degraded by endo β-1,2-glucosidase
Glucose
(L6)
846.3058
847.3113
848.3135
851.2638
811.2711
1008.3613
1206.4343
198.0969
378.1603
383.1157
d
pH
M 4 5 6 7 8 9 10
e
Temperature
M 20 30 40 50 60
f
Linear β-1,2-Glucan (0.4%)
Glucomannan (0.4%)
Carboxymethyl cellulose (0.4%)
Starch (0.4%)
Carboxymethyl curdlan (0.4%)
Laminarin (0.4%)
Lichenan (0.4%)
Barley β-Glucan (0.4%)
Tamarind xyloglucan (0.4%)
Arabinan (0.4%)
Polygalacturonic acid (0.4%)
Arabinogalactan (0.4%)

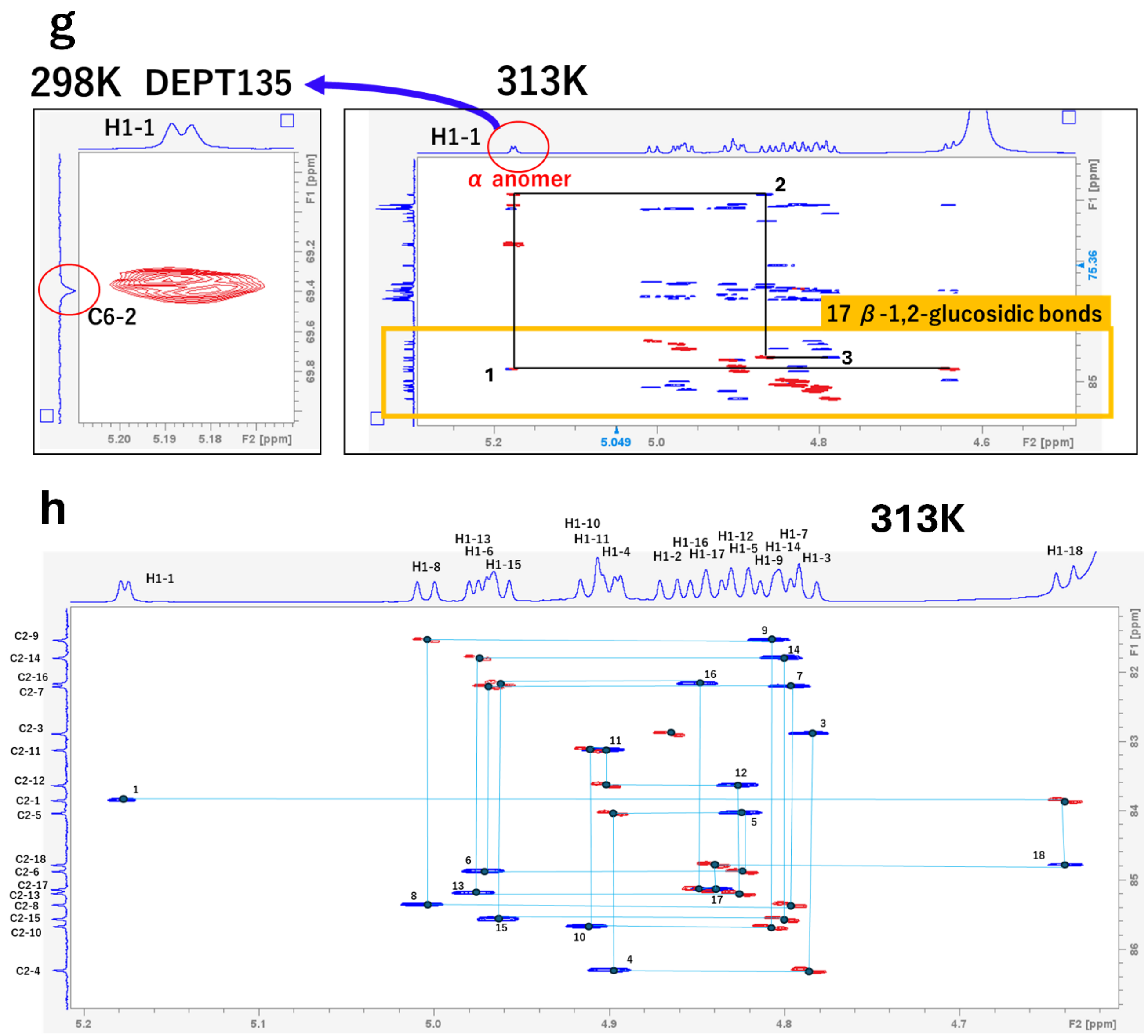


**Supplementary Figure 6. Analysis of main products released by CsOpgD2, optimal condition for catalytic reaction and substrate specificity.**

(a–c) ESI-MS analysis of the reaction products. The peaks are assigned as $[M + nNH_4]^{n+}$, and the arrows indicate the main products. The *m/z* values and peak labels are presented in the same manner as in supplementary Fig. 1. (a) Reaction products released from linear β-1,2-glucan by CsOpgD2. (b) Reaction products after BtBGL treatment. (c) Reaction products after BtBGL and CpSGL treatment. (d) TLC analysis for investigating optimal pH. Lane M, linear β-1,2-glucans (average DP 121) as a control; the other lanes, the reaction solutions of CsOpgD2 with linear β-1,2-glucans (average DP 121) incubated at respective pHs. (e) TLC analysis for investigating optimal temperature. Lane M, linear β-1,2-glucans (average DP 121) as a control; the other lanes, the reaction solutions of CsOpgD2 with linear β-1,2-glucans (average DP 121) incubated at respective temperatures. (f) Substrate specificity of CsOpgD2. The reaction was performed at 30°C. Asterisks indicate that the reaction time was 24 h. The other lanes represent a reaction time of 0 h. (g, h) 800 MHz NMR analysis of the purified main product released by CsOpgD2. Blue and red peaks represent HSQC-TOCSY and HMBC data, respectively. The H1 and C2 atoms are numbered in the same manner as in supplementary Fig. 1. (g) The trace of correlations around the α-1,6-glucosidic bond. Black lines trace from H1-3/C2-3 to H1-18/C2-1 in the direction of the non-reducing end (right, NMR analysis at 313 K). A correlation indicating the α-1,6-glucosidic bond was demonstrated by NMR analysis with DEPT135 at 298 K (left). (h) The trace of correlations derived from β-1,2-glucosidic bonds (NMR analysis at 313 K). Cyan lines trace correlations from H1-1/C2-1 to H1-3/C2-3 in the same direction as (g).

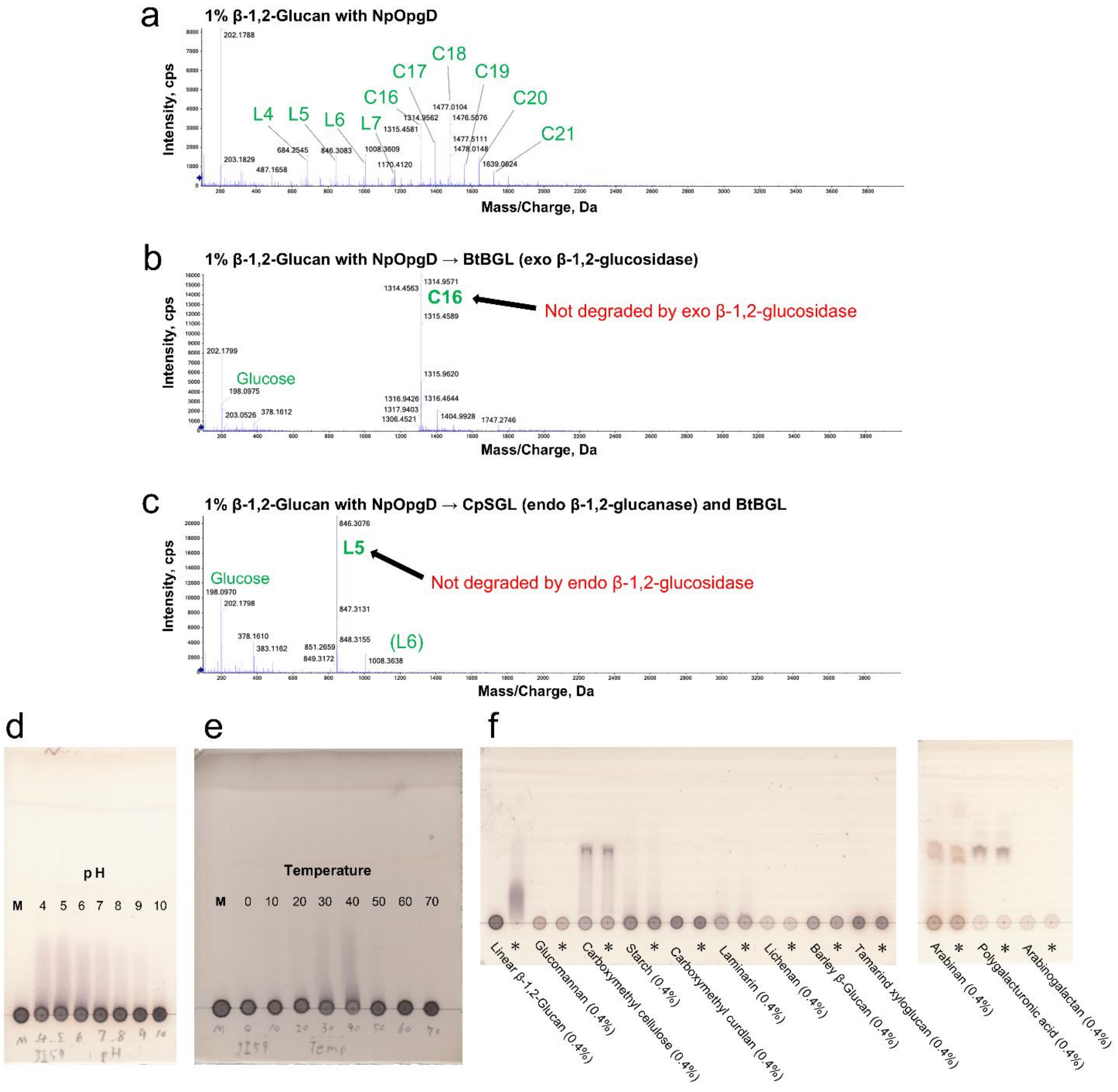


**Supplementary Figure 7. Identification of main products released by NpOpgD, optimal condition for catalytic reaction and substrate specificity.**

(a) ESI-MS analysis of the reaction products. The peaks are assigned as $[M + nNH_4]^{n+}$, and the arrows indicate the main products. The *m/z* values and peak labels are presented in the same manner as in supplementary Fig. 1. (a) Reaction products released from linear β-1,2-glucan by NpOpgD. (b) Reaction products after BtBGL treatment. (c) Reaction products after BtBGL and CpSGL treatment. (d) TLC analysis for investigating optimal pH. Lane M, linear β-1,2-glucans (average DP 121) as a control; the other lanes, the reaction solutions of NpOpgD with linear β-1,2-glucans (average DP 121) incubated at respective pHs. (e) TLC analysis for investigating optimal temperature. Lane M, linear β-1,2-glucans (average DP 121) as a control; the other lanes, the reaction solutions of NpOpgD with linear β-1,2-glucans (average DP 121) incubated at respective temperatures. (f) Substrate specificity of NpOpgD. The reaction was performed at 30°C. Asterisks indicate that the reaction time was 24 h. The other lanes represent a reaction time of 0 h.

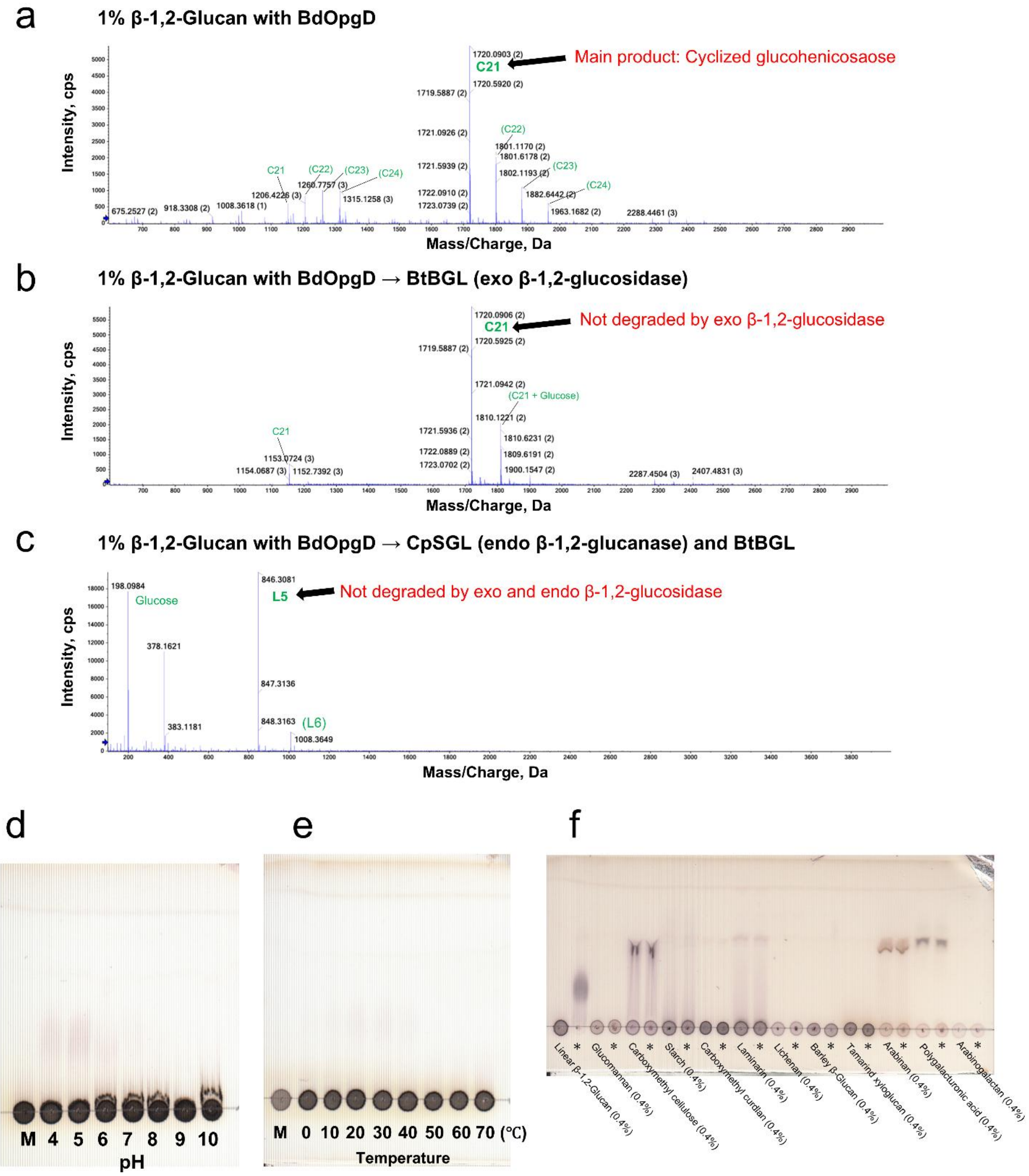


**Supplementary Figure 8. Analysis of main products released by BdOpgD, optimal condition for catalytic reaction and substrate specificity.**

(a–c) ESI-MS analysis of the reaction products. The peaks are assigned as $[M + nNH_4]^{n+}$, and the arrows indicate the main products. The *m/z* values and peak labels are presented in the same manner as in supplementary Fig. 1. (a) Reaction products released from linear β-1,2-glucan by BdOpgD. (b) Reaction products after BtBGL treatment. (c) Reaction products after BtBGL and CpSGL treatment. (d) TLC analysis for investigating optimal pH. Lane M, linear β-1,2-glucans (average DP 121) as a control; the other lanes, the reaction solutions of BdOpgD with linear β-1,2-glucans (average DP 121) incubated at respective pHs. (e) TLC analysis for investigating optimal temperature. Lane

M, linear β-1,2-glucans (average DP 121) as a control; the other lanes, the reaction solutions of BdOpgD with linear β-1,2-glucans (average DP 121) incubated at respective temperatures. (f) Substrate specificity of BdOpgD. The reaction was performed at 30°C. Asterisks indicate that the reaction time was 24 h. The other lanes represent a reaction time of 0 h.

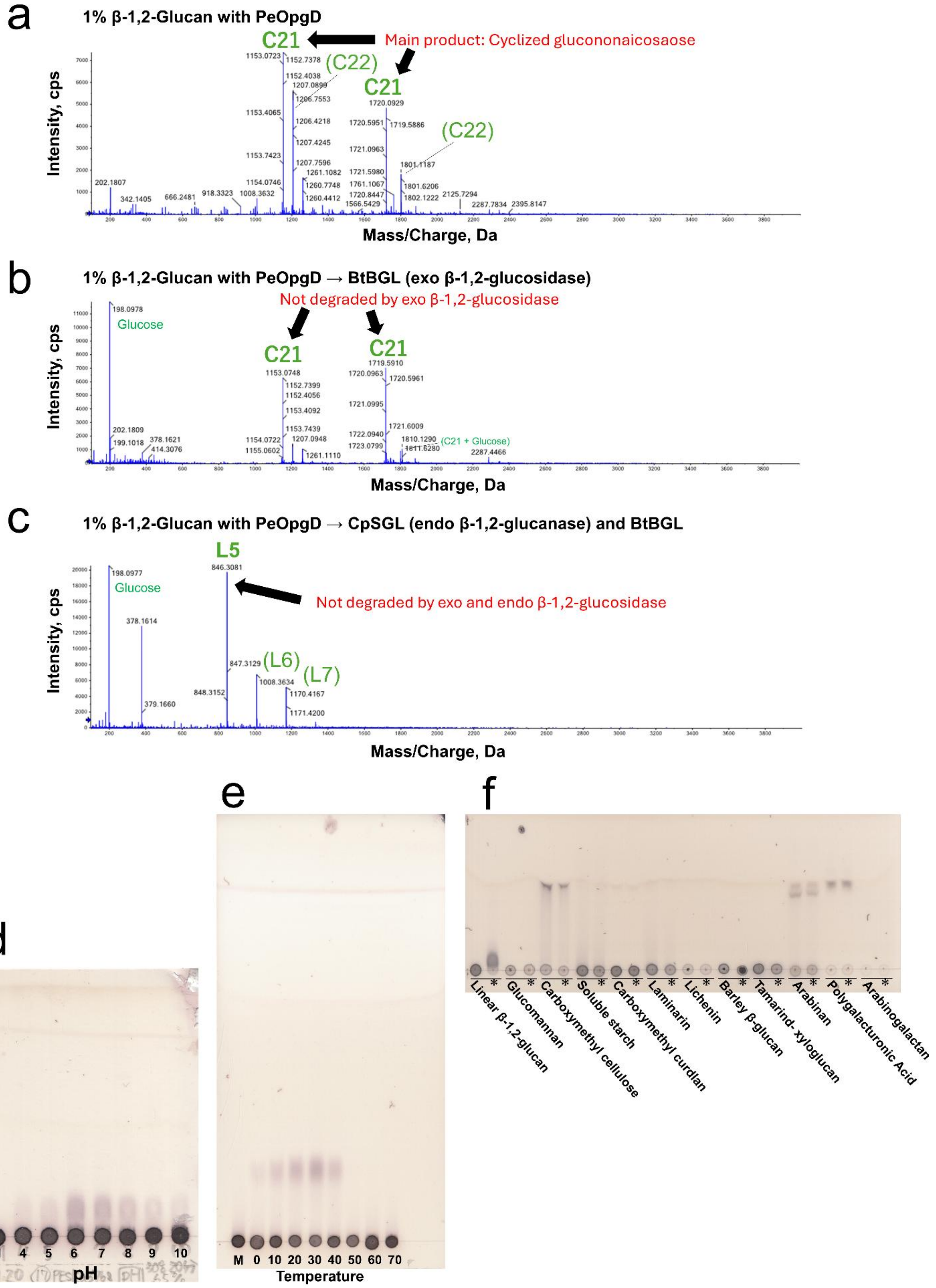
a
1% β-1,2-Glucan with PeOpgD
C21
Main product: Cyclized gluconononaicosaose
(C22)
C21
(C22)
Intensity, cps
Mass/Charge, Da
b
1% β-1,2-Glucan with PeOpgD → BtBGL (exo β-1,2-glucosidase)
Not degraded by exo β-1,2-glucosidase
Glucose
C21
C21
(C21 + Glucose)
Intensity, cps
Mass/Charge, Da
c
1% β-1,2-Glucan with PeOpgD → CpSGL (endo β-1,2-glucanase) and BtBGL
L5
Glucose
Not degraded by exo and endo β-1,2-glucosidase
(L6)
(L7)
Intensity, cps
Mass/Charge, Da
d
M 4 5 6 7 8 9 10
pH
e
M 0 10 20 30 40 50 60 70
Temperature
f
Linear β-1,2-glucan
Glucomannan
Carboxymethyl cellulose
Soluble starch
Carboxymethyl curdlan
Laminarin
Lichenin
Barley β-glucan
Tamarind- xyloglucan
Arabinan
Polygalacturonic Acid
Arabinogalactan

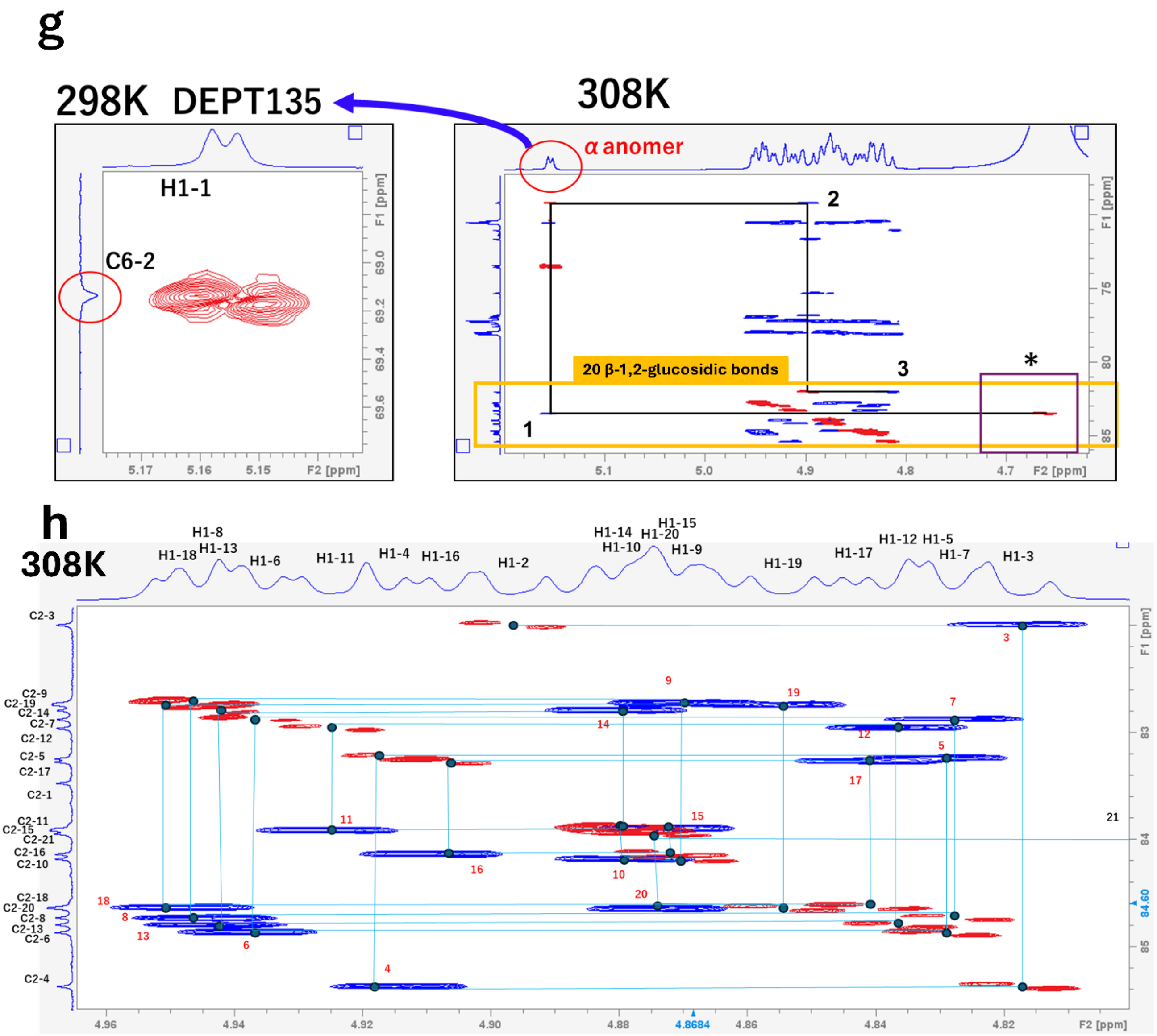


**Supplementary Figure 9. Analysis of main products released by PeOpgD, optimal condition for catalytic reaction and substrate specificity.**

(a–c) ESI-MS analysis of the reaction products. The peaks are assigned as $[M + nNH_4]^{n+}$, and the arrows indicate the main products. The *m/z* values and peak labels are presented in the same manner as in supplementary Fig. 1. (a) Reaction products released from linear β-1,2-glucan by PeOpgD. (b) Reaction products after BtBGL treatment. (c) Reaction products after BtBGL and CpSGL treatment. (d) TLC analysis for investigating optimal pH. Lane M, linear β-1,2-glucans (average DP 121) as a control; the other lanes, the reaction solutions of PeOpgD with linear β-1,2-glucans (average DP 121) incubated at respective pHs. (e) TLC analysis for investigating optimal temperature. Lane M, linear β-1,2-glucans (average DP 121) as a control; the other lanes, the reaction solutions of PeOpgD with linear β-1,2-glucans (average DP 121) incubated at respective temperatures. (f) Substrate specificity of PeOpgD. The reaction was performed at 30°C. Asterisks indicate that the reaction time was 24 h. The other lanes represent a reaction time of 0 h. (g) H1 proton NMR analysis of the main product of Experiment 1 by PeOpgD. (g, h) NMR analysis of the purified main product released by PeOpgD. Blue and red peaks represent HSQC-TOCSY and HMBC data, respectively. The H1 and C2 atoms are numbered in the same manner as in supplementary Fig. 1. (g) The trace of correlations around the α-1,6-glucosidic bond. Black lines trace from H1-3/C2-3 to H1-21/C2-1 in the direction of the non-reducing end (right, NMR analysis at 308 K). A correlation indicating the α-1,6-glucosidic bond was demonstrated by NMR analysis with DEPT135 298 K (left). Astarisk indicates the predicted position of the HSQC peak at H1-21/C2-21. The positon is invisble due to the presence of the peak of $D_2O$ [The presences of 20 HSQC peaks of H1-X/C2-X which can be sequentially traced with HMBC analysis were indicated in 333K NMR (Data not shown)]. (h) The trace of correlations derived from β-1,2-glucosidic bonds (NMR analysis at 308 K). Cyan lines trace from H1-20/C2-21 to H1-2/C2-3 in the same direction as (g).

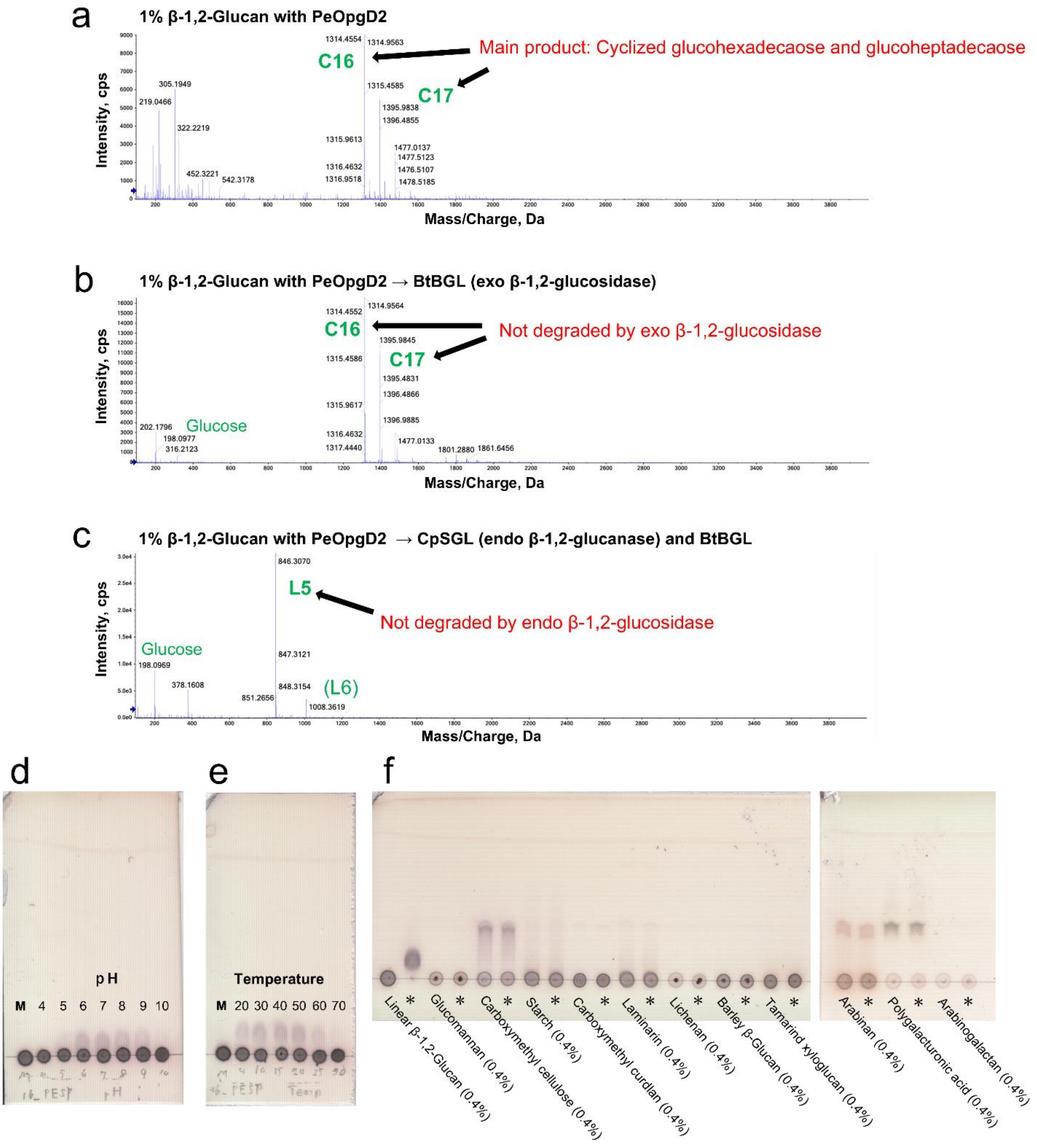


**Supplementary Figure 10. Analysis of main products released by PeOpgD2, optimal condition for catalytic reaction and substrate specificity.**

(a–c) ESI-MS analysis of the reaction products. The peaks are assigned as $[M + nNH_4]^{n+}$, and the arrows indicate the main products. The *m/z* values and peak labels are presented in the same manner as in supplementary Fig. 1. (a) Reaction products released from linear β-1,2-glucan by PeOpgD2. (b) Reaction products after BtBGL treatment. (c) Reaction products after BtBGL and CpSGL treatment. (d) TLC analysis for investigating optimal pH. Lane M, linear β-1,2-glucans (average DP 121) as a control; the other lanes, the reaction solutions of PeOpgD with linear β-1,2-glucans (average DP 121) incubated at respective pHs. (e) TLC analysis for investigating optimal temperature. Lane

M, linear β-1,2-glucans (average DP 121) as a control; the other lanes, the reaction solutions of PeOpgD2 with linear β-1,2-glucans (average DP 121) incubated at respective temperatures. (f) Substrate specificity of PeOpgD2. The reaction was performed at 30°C. Asterisks indicate that the reaction time was 24 h. The other lanes represent a reaction time of 0 h.

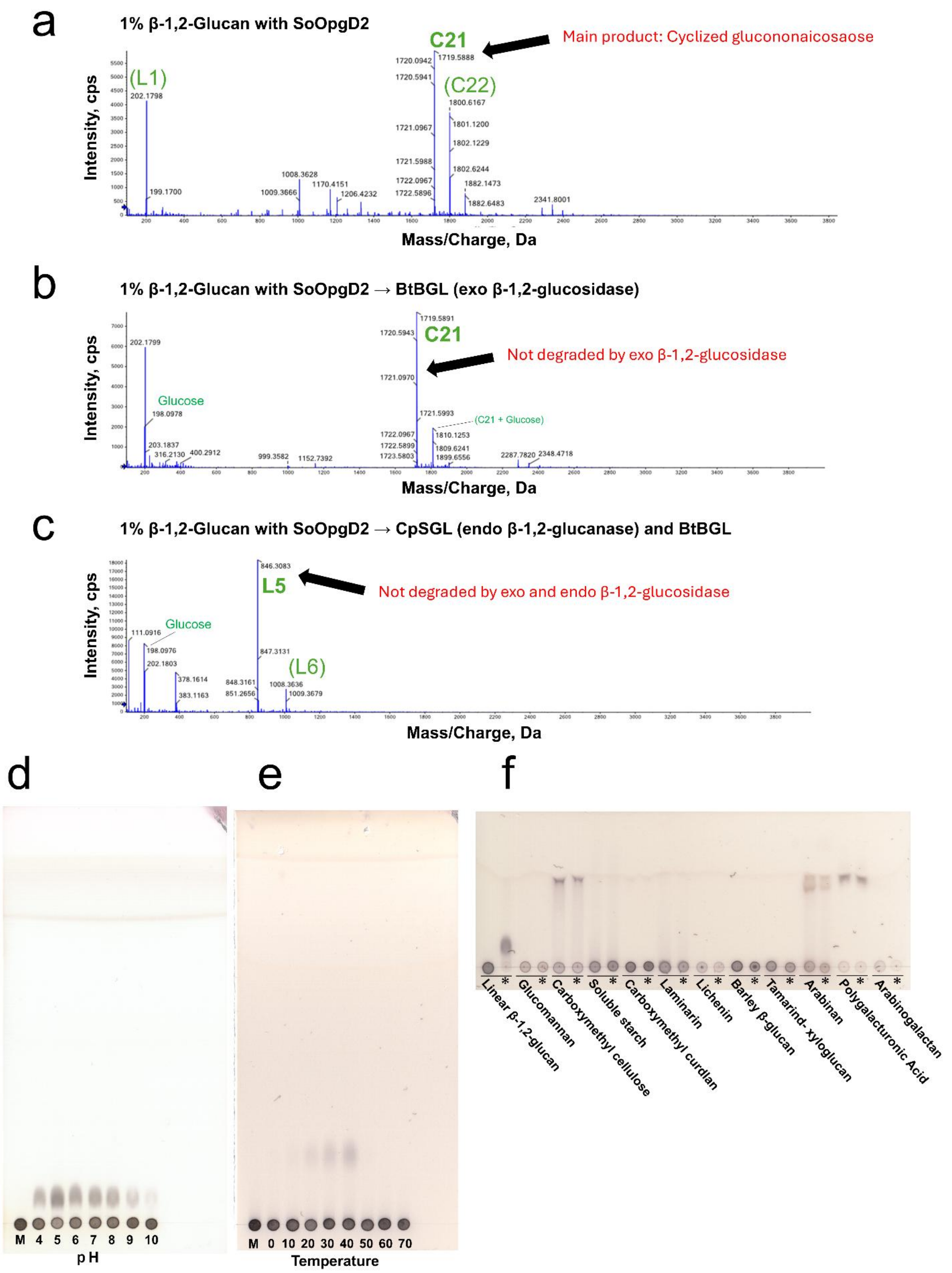


**Supplementary Figure 11. Analysis of main products released by SoOpgD2, optimal condition for catalytic reaction and substrate specificity.**

(a–c) ESI-MS analysis of the reaction products. The peaks are assigned as $[M + nNH_4]^{n+}$, and the arrows indicate the main products. The *m/z* values and peak labels are presented in the same manner as in supplementary Fig. 1. (a) Reaction products released from linear β-1,2-glucan by SoOpgD2. (b) Reaction products after BtBGL treatment. (c) Reaction products after BtBGL and CpSGL treatment. (d) TLC analysis for investigating optimal pH. Lane M, linear β-1,2-glucans (average DP 121) as a control; the other lanes, the reaction solutions of SoOpgD2 with linear β-1,2-glucans (average DP 121) incubated at respective pHs. (e) TLC analysis for investigating optimal temperature. Lane M, linear β-1,2-glucans (average DP 121) as a control; the other lanes, the reaction solutions of SoOpgD2 with linear β-1,2-glucans (average DP 121) incubated at respective temperatures. (f) Substrate specificity of SoOpgD2. The reaction was performed at 30°C. Asterisks indicate that the reaction time was 24 h. The other lanes represent a reaction time of 0 h.

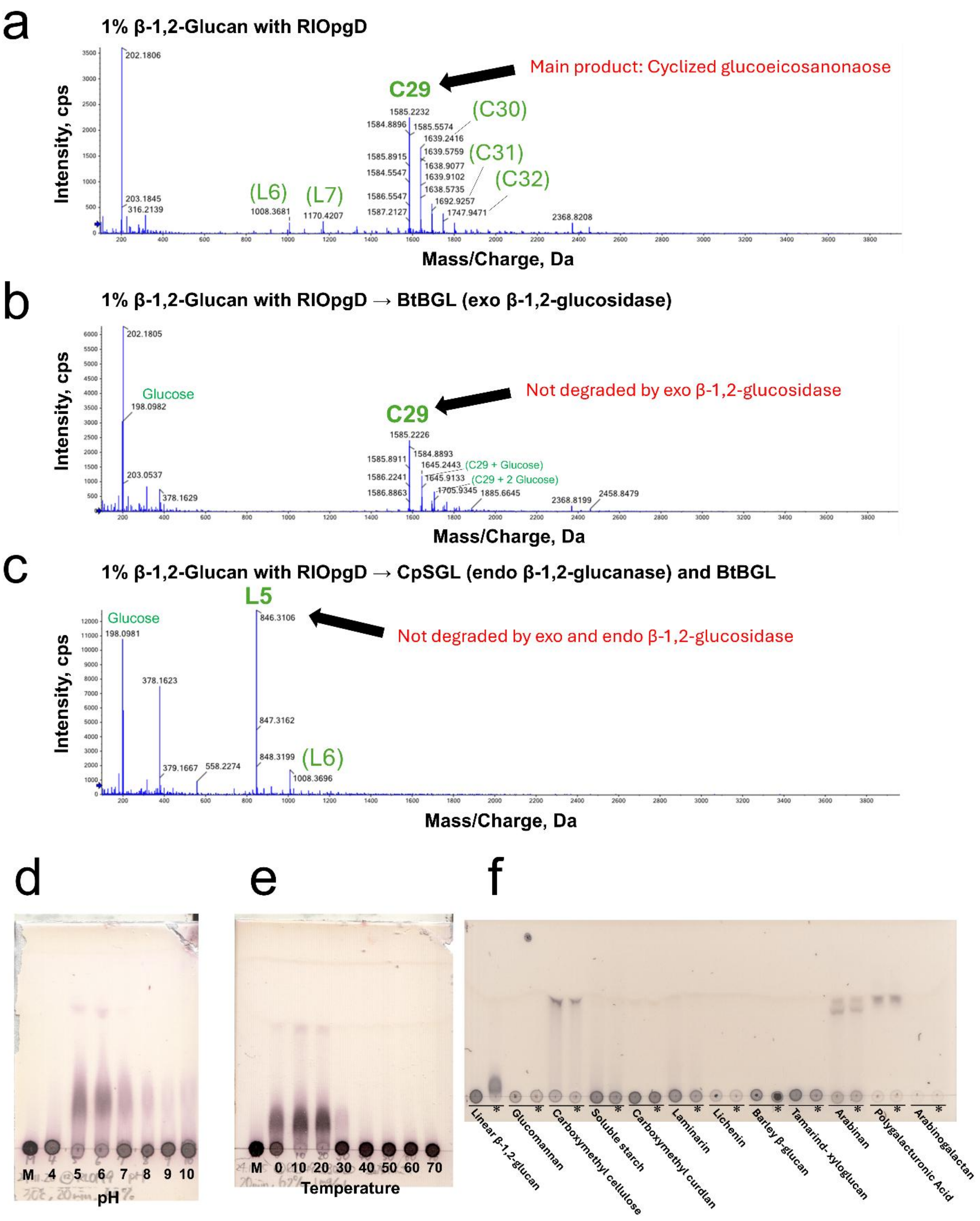


**Supplementary Figure 12. Analysis of main products released by RlOpgD, optimal condition for catalytic reaction and substrate specificity.**

(a–c) ESI-MS analysis of the reaction products. The peaks are assigned as $[M + nNH_4]^{n+}$, and the arrows indicate the main products. The *m/z* values and peak labels are presented in the same manner as in supplementary Fig. 1. (a) Reaction products released from linear β-1,2-glucan by RlOpgD. (b) Reaction products after BtBGL treatment. (c)

Reaction products after BtBGL and CpSGL treatment. (d) TLC analysis for investigating optimal pH. Lane M, linear β-1,2-glucans (average DP 121) as a control; the other lanes, the reaction solutions of RlOpgD with linear β-1,2-glucans (average DP 121) incubated at respective pHs. (e) TLC analysis for investigating optimal temperature. Lane M, linear β-1,2-glucans (average DP 121) as a control; the other lanes, the reaction solutions of RlOpgD with linear β-1,2-glucans (average DP 121) incubated at respective temperatures. (f) Substrate specificity of RlOpgD. The reaction was performed at 30°C. Asterisks indicate that the reaction time was 24 h. The other lanes represent a reaction time of 0 h.

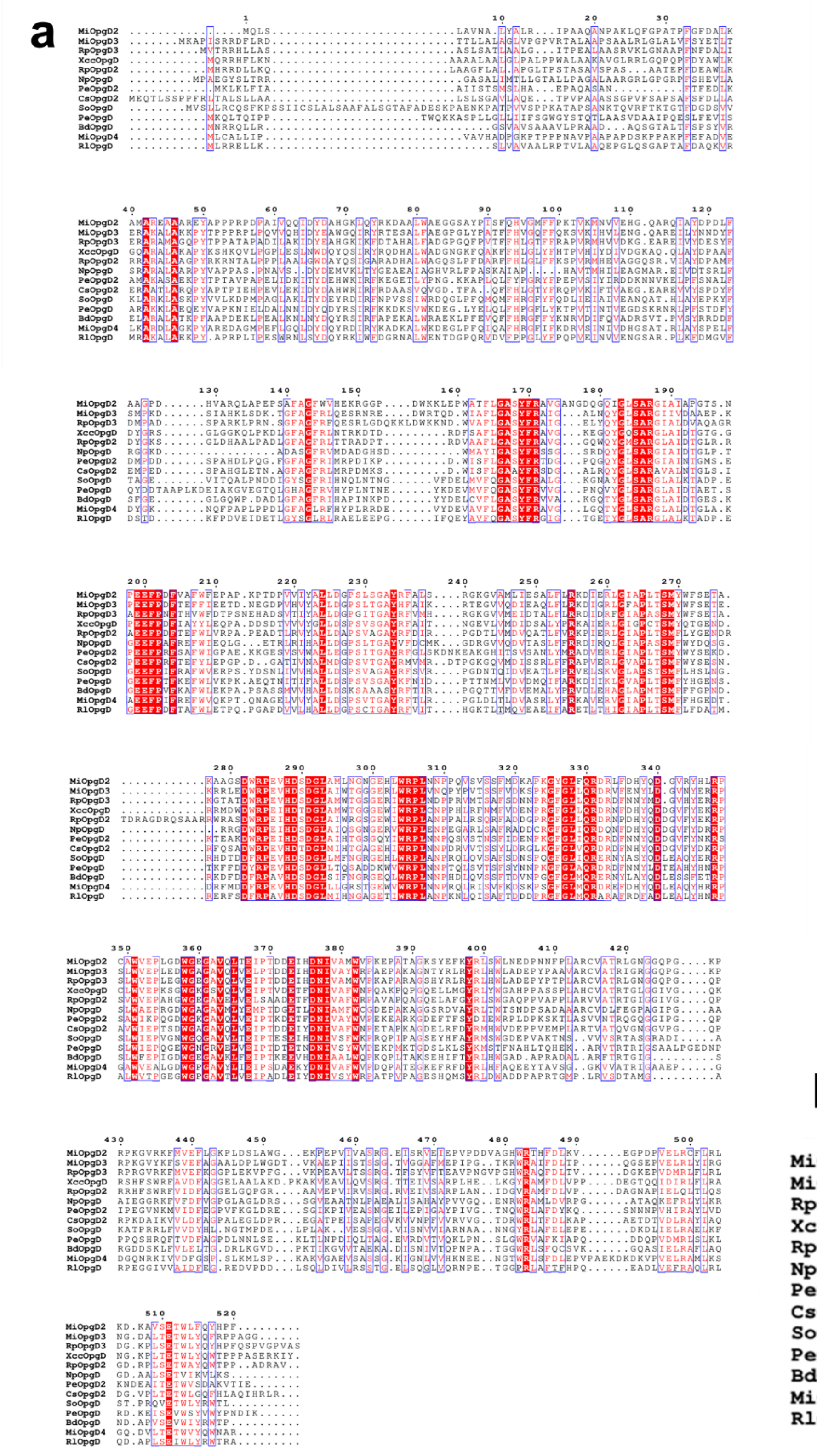

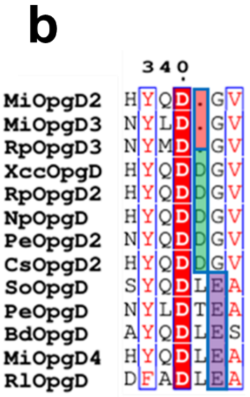

**Supplementary Figure 13. Amino acid sequence aliment among the 12 homologs in this study/ and XccOpgD.** (a) Overall aliment. 12 GH186 homologs in this study and XccOpgD are aligned by ClustalW in MEGA11[1] and visualized by Espript3[2]. (b) Aliment around Loop X. The sequence around loop X is shown for each cluster.

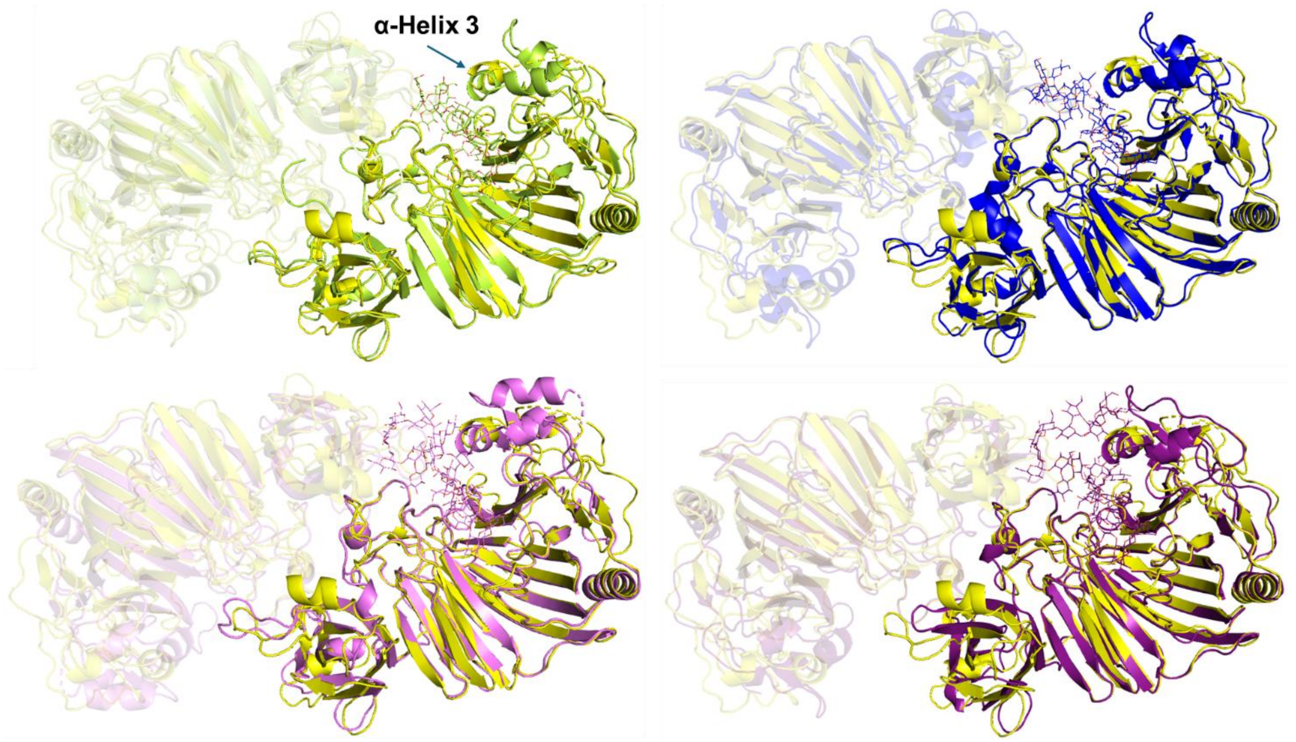


**Supplementary Figure 14. Superposition between XccOpgD (chain B) and Chain A of RpOpgD3, CsOpgD2, PeOpgD and RlOpgD.**

Complex structures of XccOpgD, RpOpgD3, CsOpgD2, PeOpgD and RlOpgD are distinguished by color; yellow, light green, blue, purple and deep purple, respectively. Chain A of XccOpgD and chains B of RpOpgD3, CsOpgD2, PeOpgD and RlOpgD are shown semi-transparently. Substrates in catalytic clefts of chain A are shown as lines. The other substrates are omitted.

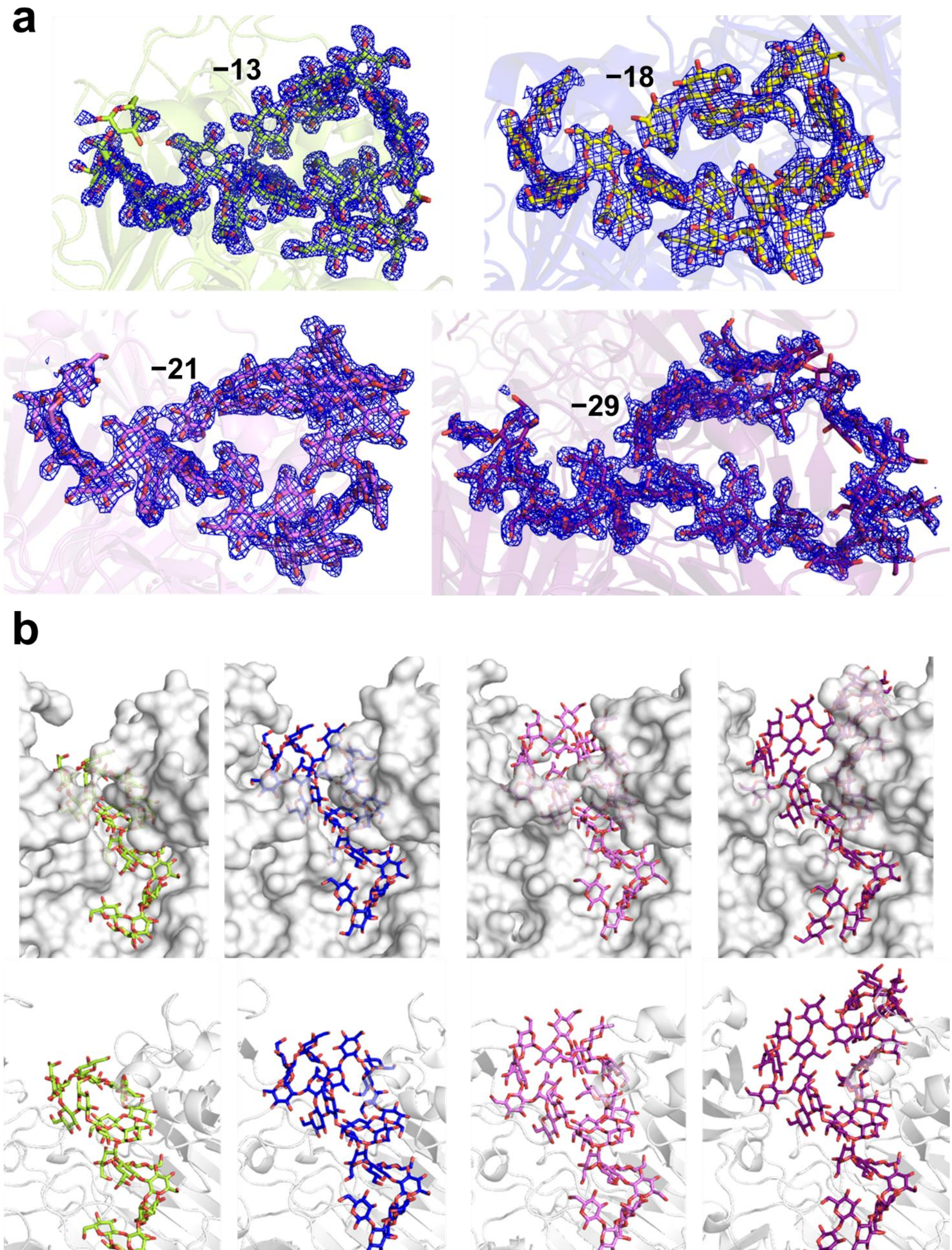
a
−13
−18
−21
−29
b

**Supplementary Figure 15. Overall substrate binding modes.**

(a) $F_o$-$F_c$ omit maps of the substrate at the catalytic cleft of each homolog. The substrates at the catalytic centers of RpOpgD3, CsOpgD2, PeOpgD and RlOpgD are shown as light green, yellow, purple and deep purple sticks, respectively. $F_o$-$F_c$ omit maps of the substrates in RpOpgD3, CsOpgD2, PeOpgD and RlOpgD are shown as blue meshes at 2.5, 2.0, 2.5 and 2.0 σ contour levels, respectively.

(b) Overall substrate binding modes of RpOpgD3, CsOpgD2, PeOpgD and RlOpgD. Each homolog is shown as semi-transparent cartoon. The substrates at the catalytic centers of RpOpgD3, CsOpgD2, PeOpgD and RlOpgD are shown as light green, blue, purple and deep purple sticks, respectively.

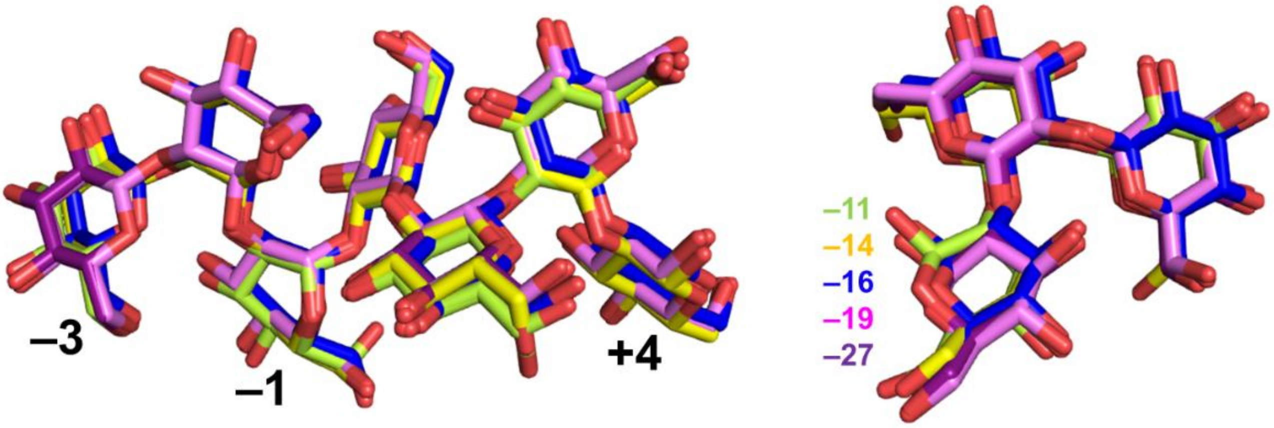


**Supplementary Figure 16**. **Common subsites among RpOpgD3, XccOpgD, CsOpgD2, PeOpgD and RlOpgD.** The Michaelis complexes of XccOpgD (chain B) and the GH186 homologs in this study (chains A) are superimposed based on the positions of the catalytic centers. The glucose moieties at subsites −3 to +4 and the three glucose moieties from the non-reducing ends of RpOpgD3, CsOpgD2, PeOpgD and RlOpgD are shown as light green, blue, purple and deep purple sticks, respectively. (a) Subsites −3 to +4. (b) The three glucose moieties from the non-reducing ends.

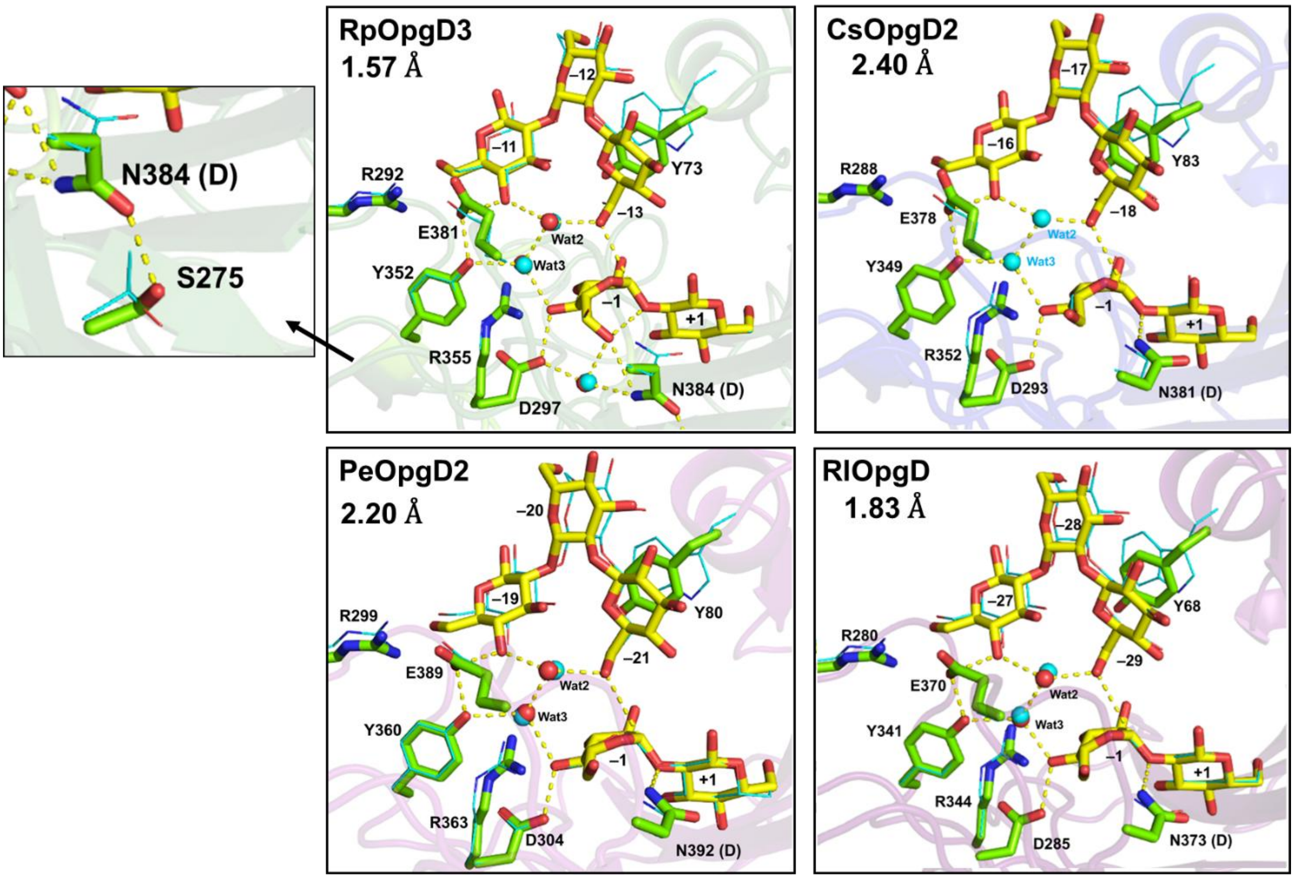


**Supplementary Figure 17. Catalytic centers of four Michaelis complexes of the GH186 enzymes.**

Hydrogen bonds with appropriate distances are shown as yellow dotted lines. Residues and substrates are shown as green and yellow sticks, respectively. Each of the four structures is superimposed with the Michaelis complex of XccOpgD (PDB ID: 8X18). XccOpgD complex is shown as cyan lines and water molecules in the complex are shown as cyan spheres. Parentheses represent the original residues in the wild-type enzymes.

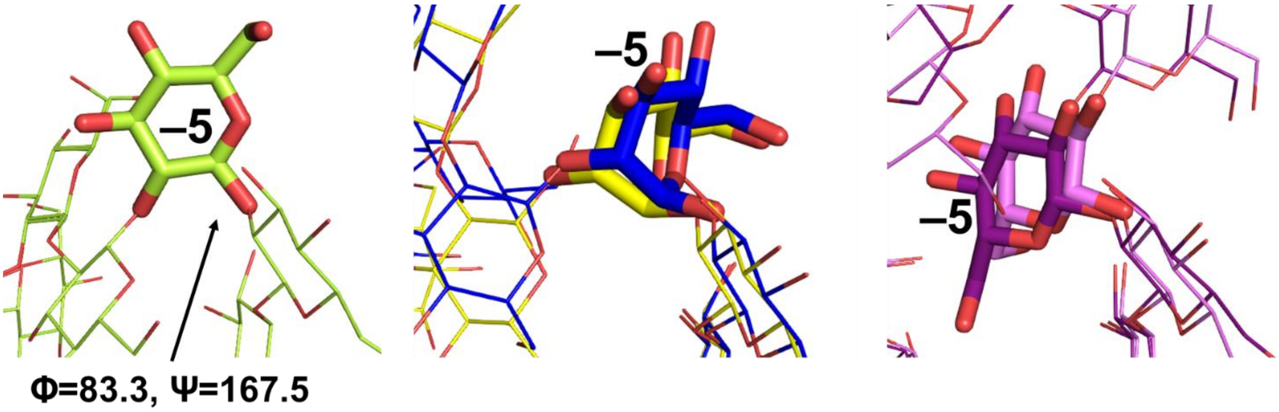


**Supplementary Figure 18. Differences in conformations at subsites −5 of four GH186 enzymes.**

The Michaelis complexes of XccOpgD (chain B) and GH186 homologs in this study (chains A) are superimposed based on the positions of the catalytic centers. Each glucose moiety at subsite −5 is shown as a stick and the other glucose moieties are shown as lines. Complex structures of XccOgpD, RpOpgD3, CsOpgD2, PeOpgD and RlOpgD are distinguished by color; yellow, light green, blue, purple and deep purple, respectively. (a) Subsite −5 of RpOpgD3. (b) Subsites −5 of XccOpgD and CsOpgD2. (c) Subsites −5 of PeOpgD and RlOpgD.

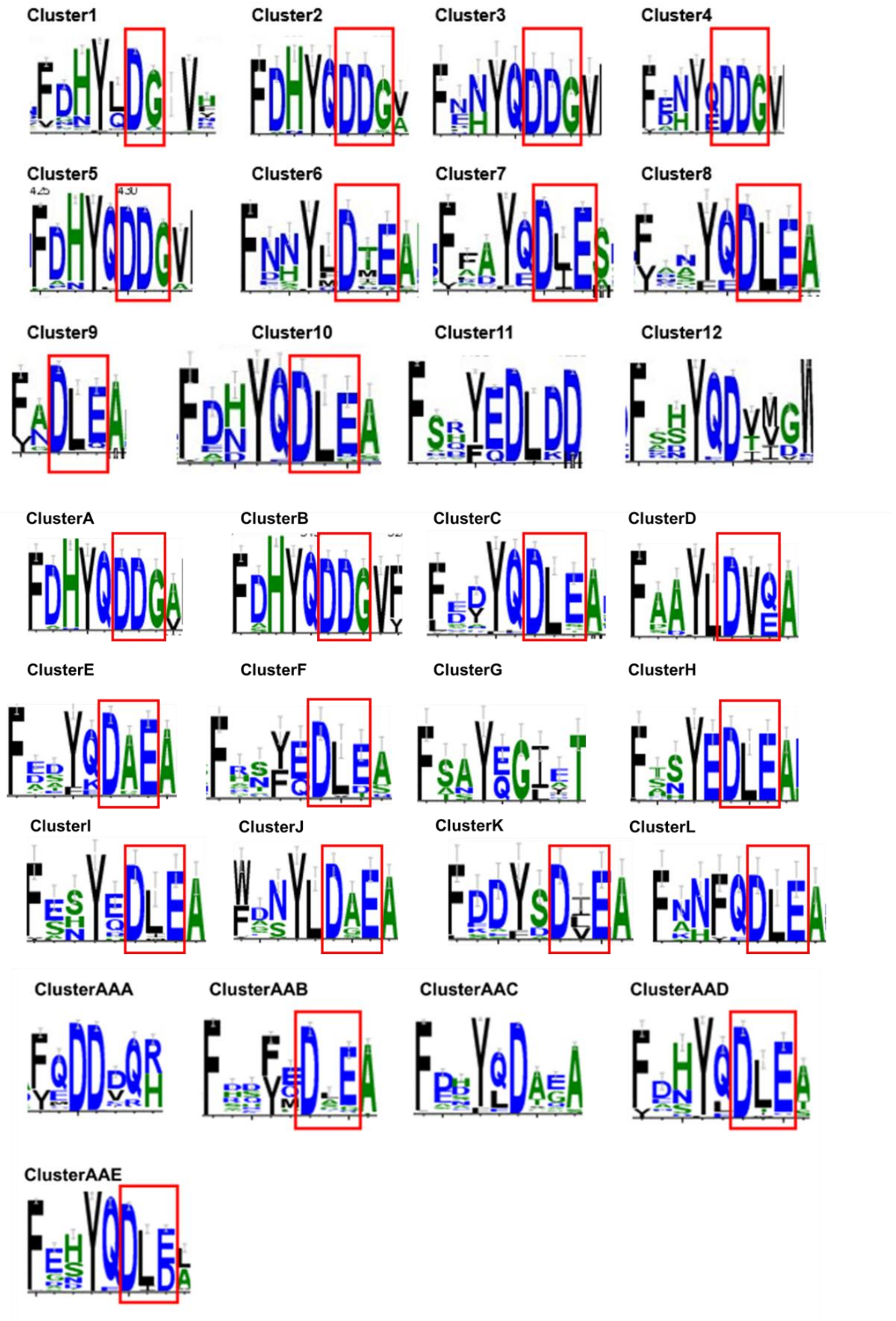


**Supplementary Figure 19. Sequence logo analyses at Loop X.**

Each cluster on SSN analysis is aligned by ClustalW in MEGA11[1] and visualized by WebLogo3 server (https://weblogo.threeplusone.com/create.cgi). The sequence logos around loop X is shown for each cluster. Red boxes indicate the typical motifs of Loop X (D-G, DDG and DxE motifs).

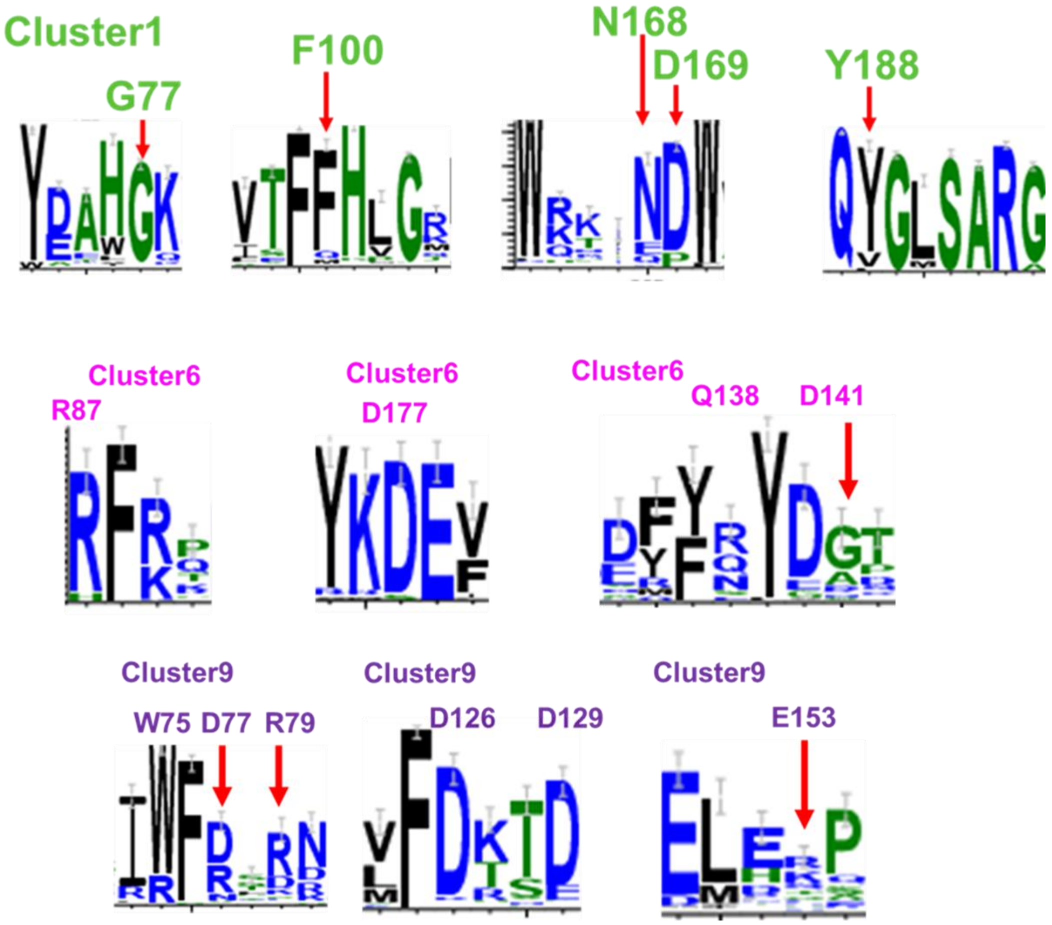


**Supplementary Figure 20. Sequence logo analyses for substrate recognition residues.**

Each cluster on SSN analysis is aligned by ClustalW in MEGA11[1] and visualized by WebLogo3 server (https://weblogo.threeplusone.com/create.cgi). The sequence logos are prepared in the same way as Supplementary figure 19. The sequence logos around the notable substrate recognition residues are shown for clusters 1, 6 and 9.

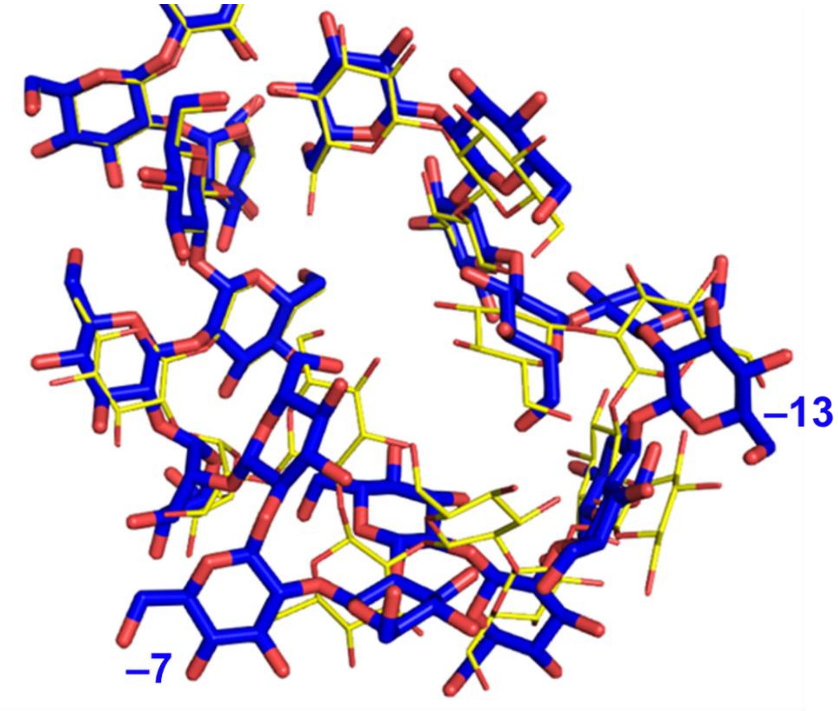


**Supplementary Figure 21. Structural differences in the configuration at minus subsites between XccOpgD and CsOpgD2.**

The glucose moieties at minus subsites of XccOpgD (thin yellow stick) and CsOpgD2 (blue stick) are superimposed. The glucose moieties at subsites −7 and −13 are indicated by letter.

**Table S1. Chemical shifts of the α-1,6-cyclized β-1,2-glucooligosaccharides.**

**a. H-1 proton chemical shifts of the cyclic hexadecaose produced by Experiment 1 of RpOpgD2**

800 MHz NMR ($D_2O$ with internal standard tBuOH $\delta$ = 1.23, 298 K)

| Proton | Chemical shift of each glucose residue (ppm) | | | | | | | | | | | | | | | |
|---|---|---|---|---|---|---|---|---|---|---|---|---|---|---|---|---|
| | 1 | 2 | 3 | 4 | 5 | 6 | 7 | 8 | 9 | 10 | 11 | 12 | 13 | 14 | 15 | 16 |
| H-1 (ppm) | 5.160 | 4.901 | 4.796 | 4.968 | 4.872 | 4.991 | 4.820 | 5.035 | 4.843 | 4.913 | 4.991 | 4.801 | 5.010 | 4.839 | 4.920 | 4.651 |
| $J_{1,2}$ (Hz) | 3.68 | 7.51 | 8.15 | 7.11 | 7.91 | 7.91 | 7.43 | 7.83 | 7.35 | 7.75 | 7.91 | 7.91 | 7.99 | 7.99 | 7.83 | 7.99 |

**b. H-1 proton chemical shifts and C-2, C-6 carbon chemical shifts of the cyclic didecaose produced by Experiment 1 of MiOpgD2**

700 MHz NMR ($D_2O$ with internal standard tBuOH δ = 1.23, 333 K)

| Proton | Chemical shift of each glucose moiety (ppm) | | | | | | | | | | | |
|---|---|---|---|---|---|---|---|---|---|---|---|---|
| | 1 | 2 | 3 | 4 | 5 | 6 | 7 | 8 | 9 | 10 | 11 | 12 |
| H-1 (ppm) | 5.132 | 4.891 | 4.892 | 4.899 | 4.958 | 4.976 | 5.084 | 4.899 | 5.002 | 4.924 | 4.994 | 4.665 |
| $J_{1,2}$ (Hz) | 3.43 | 7.81 | 7.81 | 7.81 | 7.79 | 7.15 | 7.74 | 7.81 | 7.69 | 7.79 | 7.28 | 7.78 |
| C-2 (ppm) | * | - | * | * | * | * | * | * | * | * | * | * |
| C-6 (ppm) | | 69.61 | | | | | | | | | | |

*: 80.48-85.26ppm

**c. H-1 proton chemical shifts and C-2, C-6 carbon chemical shifts of the cyclic tridecaose produced by Experiment 1 of RpOpgD3**

800 MHz NMR (D2O with internal standard tBuOH δ = 1.23, 318 K)

| Proton | Chemical shift of each glucose moiety (ppm) | | | | | | | | | | | | |
|---|---|---|---|---|---|---|---|---|---|---|---|---|---|
| | 1 | 2 | 3 | 4 | 5 | 6 | 7 | 8 | 9 | 10 | 11 | 12 | 13 |
| H-1 (ppm) | 5.163 | 4.924 | 4.901 | 5.048 | 5.008 | 4.889 | 5.027 | 4.877 | 5.081 | 4.901 | 4.912 | 4.924 | 4.644 |
| $J_{1,2}$ (Hz) | 3.22 | 8.19 | 7.42 | 7.70 | 7.70 | 9.48 | 7.77 | 8.05 | 6.99 | 7.42 | 7.98 | 8.19 | 7.76 |
| C-2 (ppm) | * | | * | * | * | * | * | * | * | * | * | * | * |
| C-6 (ppm) | | 69.27 | | | | | | | | | | | |

*: 80.27-85.56 ppm

**d. H-1 proton chemical shifts and C2, C6 carbon chemical shifts of the cyclic octadecaose produced by Experiment 1 of CsOpgD2**

800 MHz NMR ($D_2O$ with internal standard tBuOH $\delta$ = 1.23, 313 K)

| Proton | Chemical shift of each glucose moiety (ppm) | | | | | | | | | | | | | | | | | |
|---|---|---|---|---|---|---|---|---|---|---|---|---|---|---|---|---|---|---|
| | 1 | 2 | 3 | 4 | 5 | 6 | 7 | 8 | 9 | 10 | 11 | 12 | 13 | 14 | 15 | 16 | 17 | 18 |
| H-1 (ppm) | 5.176 | 4.866 | 4.786 | 4.898 | 4.825 | 4.970 | 4.797 | 5.004 | 4.809 | 4.911 | 4.902 | 4.825 | 4.975 | 4.801 | 4.962 | 4.849 | 4.840 | 4.640 |
| $J_{1,2}$ (Hz) | 3.94 | 7.89 | 8.12 | 8.03 | 7.89 | 7.76 | 8.17 | 8.02 | 7.77 | 8.31 | 8.00 | 7.89 | 8.29 | 7.60 | 8.02 | 7.98 | 7.98 | 7.77 |
| C-6 (ppm) | | 69.42 | | | | | | | | | | | | | | | | |
| C-2 (ppm) | * | - | * | * | * | * | * | * | * | * | * | * | * | * | * | * | * | * |

*: 81.53-86.29 ppm

**e. H-1 proton chemical shifts and C2, C6 carbon chemical shifts of the cyclic henicosaose produced by Experiment 1 of PeOpgD**

800 MHz NMR ($D_2O$ with internal standard tBuOH $\delta$ = 1.23, 308 K)

| Proton | Chemical shift of each glucose moiety (ppm) | | | | | | | | | | | | | | | |
|---|---|---|---|---|---|---|---|---|---|---|---|---|---|---|---|---|
| | 1 | 2 | 3 | 4 | 5 | 6 | 7 | 8 | 9 | 10 | 11 | 12 | 13 | 14 | 15 | 16 |
| H-1 (ppm) | 5.153 | 4.897 | 4.818 | 4.915 | 4.830 | 4.937 | 4.827 | 4.945 | 4.870 | 4.879 | 4.924 | 4.837 | 4.944 | 4.879 | 4.871 | 4.908 |
| $J_{1,2}$ (Hz) | 3.42 | 7.80 | 7.74 | 8.29 | 8.25 | 7.50 | 7.39 | 7.96 | 7.26 | 8.07 | 8.05 | 8.04 | 7.79 | 7.22 | 7.23 | 7.94 |
| C-6 (ppm) | | 69.17 | | | | | | | | | | | | | | |
| C-2 (ppm) | * | | * | * | * | * | * | * | * | :* | * | * | * | * | * | * |

| Proton | Chemical shift of each glucose moiety (ppm) | | | | |
|---|---|---|---|---|---|
| | 17 | 18 | 19 | 20 | 21 |
| H-1 (ppm) | 4.840 | 4.947 | 4.855 | 4.874 | *’ |
| $J_{1,2}$ (Hz) | 8.24 | 7.92 | 7.96 | 7.51 | *’ |
| C-6 (ppm) | | | | | |
| C-2 (ppm) | * | * | * | * | * |

*: 82.01-85.34 ppm

*’: The peak of H1-21 is not observed due to the overlapping with the peak of $D_2O$.

**Supplementary Table 2. Crystallographic data collection and refinement statistics**

| Data set | RpOpgD3-β-1,2-glucan (D384N mutant) | CsOpgD2-β-1,2-glucan (D381N mutant) | PeOpgD-β-1,2-glucan (D392N mutant) | RlOpgD-β-1,2-glucan (D373N mutant) |
|---|---|---|---|---|
| **Data collection** | | | | |
| Beamline | KEK BL-5A | KEK BL-5A | KEK BL-5A | KEK BL-5A |
| Space group | *C* 1 2 1 | *P* $4_3$ $2_1$ 2 | *I* 2 2 2 | *P* 2 $2_1$ $2_1$ |
| Unit cell parameters (Å, °) | *a* = 151.50<br>*b* = 54.48<br>*c* = 79.62<br>α, γ = 90<br>β= 97.59 | a = 78.50<br>b = 78.50<br>c = 431.30<br>α, β, γ = 90 | a = 89.00<br>b = 117.59<br>c = 119.58<br>α, β, γ = 90 | a = 47.57<br>b = 110.70<br>c = 207.05<br>α, β, γ = 90 |
| Resolution (Å)[a] | 43.87-1.57 (1.60-1.57) | 46,68-2.40 (2.47-2.40) | 45.73-2.20 (2.27-2.20) | 48.81-1.83 (1.86-1.83) |
| Total reflections[a] | 284733 (13139) | 338268 (28220) | 213104 (18680) | 618183 (31547) |
| Unique reflections[a] | 88087 (4272) | 51173 (4452) | 32177 (2763) | 97564 (4725) |
| Completeness (%)[a] | 98.1 (97.0) | 95.2 (96.8) | 100 (100) | 100 (100) |
| Multiplicity[a] | 3.2 (3.1) | 6.6 (6.3) | 6.6 (6.8) | 6.3 (6.7) |
| Mean $I/\sigma(I)$[a] | 14.2 (2.2) | 19.0 (2.7) | 9.8 (2.3) | 18.6 (2.2) |
| $R_{merge}$ (%)[a] | 4.6 (47.2) | 8.2 (82.7) | 23.0 (91.6) | 7.4 (76.3) |
| $R_{pim}$ (%)[a] | 5.6 (57.1) | 3.4 (34.6) | 9.6 (37.8) | 3.2 (31.9) |
| $CC_{1/2}$[a] | (0.843) | (0.927) | (0.615) | (0.826) |
| **Refinement** | | | | |
| Resolution (Å) | 43.87-1.57 | 46.72-2.40 | 45.73-2.20 | 46.93-1.83 |
| No. of reflections | 88048 | 50861 | 32176 | 97183 |
| No. of atoms | 4946 | 8760 | 4591 | 9225 |
| No. of water molecules | 386 | 54 | 268 | 564 |
| $R_{work}/R_{free}$ | 0.166/0.186 | 0.229/0.276 | 0.177/0.222 | 0.187/0.230 |
| No. of asymmetric units | 1 | 2 | 1 | 2 |
| r.m.s.d. from ideal values | | | | |
| Bond lengths (Å) | 0.011 | 0.011 | 0.010 | 0.014 |
| Bond angles (°) | 1.649 | 1.488 | 1.709 | 1.507 |
| Average *B*-factors (Å$^2$) | 22.0 | 37.0 | 21.0 | 29.0 |
| Ramachandran plot (%) | | | | |
| Favored | 97 | 97 | 96 | 96 |
| Allowed | 3 | 3 | 4 | 4 |
| Outlier | 0 | 0 | 0 | 0 |
| **PDB entry** | 22PV | 22PX | 22PZ | 22QA |

[a]Values in parentheses represent the highest resolution shell.

Supplementary Table 3. Primer pairs for recombinant enzymes

| | Forward primer | Reverse primer |
|---|---|---|
| CsOpgD2 WT | actttaagaaggagatatacatatgcaggaaacccccgtccccgcg | agtggtggtggtggtggtgctcgaggcgcaggcgatggatctgggccag |
| CsOpgD2 D381N | atctacaataatatcgtggccttctgg | tggctgctgctctagatgttattatatg |
| RpOpgD2 WT | actttaagaaggagatatacatatggccagccgcgtcaagctcggcaat | agtggtggtggtggtggtgctcgaggctcgccaccggccccaccggcga |
| RpOpgD3 WT | actttaagaaggagatatacatatggccagccgcgtcaagctcggcaat | agtggtggtggtggtggtgctcgaggctcgccaccggccccaccggcga |
| RpOpgD3 D384N | atccacaataacatcgtcgccatgtgg | tggctgctactctaggtgttattgtag |
| BdOpgD WT | actttaagaaggagatatacatatgcagtccggaacggccctcacgttc | agtggtggtggtggtggtgctcgaggggcgtccatcgatagatccagac |
| NpOpgD WT | actttaagaaggagatatacatatggcgcgcggcaggcttggtcc | agtggtggtggtggtggtgctcgagtgacttcagcacctttatcacagt |
| RlOpgD WT | actttaagaaggagatatacatatggcgcaggaacccgggctgcaatcg | agtggtggtggtggtggtgctcgagtgacttcagcacctttatcacagt |
| RlOpgD2 D373N | atctacaataacatcgtgtcttattgg | tgactagacctttagatgttattgtag |
| SoOpgD2 WT | actttaagaaggagatatacatatggcgcaggaacccgggctgcaatcg | agtggtggtggtggtggtgctcgagtgacttcagcacctttatcacagt |
| PeOpgD WT | actttaagaaggagatatacatatggctagcgttgatgccgctattcca | agtggtggtggtggtggtgctcgagcttgatgtcatttgggtaccaaac |
| PeOpgD D392N | actaacaataatattgtgagttactgg | tgtctatgactttgattgttattataa |
| PeOpgD2 WT | actttaagaaggagatatacatatgagtgccaattttacatttgaagac | agtggtggtggtggtggtgctcgagcttgatgtcatttgggtaccaaac |
| MiOpgD2 WT | actttaagaaggagatatacatatggatcccggcaagcccacgcccccg | agtggtggtggtggtggtgctcgagtcgggcgttccactggtaaaccca |
| MiOpgD3 WT | actttaagaaggagatatacatatggccccgagcgccgccctgcgcctg | agtggtggtggtggtggtgctcgaggcctccggcgggaggccggaactg |
| MiOpgD4 WT | actttaagaaggagatatacatatggatcccggcaagcccacgcccccg | agtggtggtggtggtggtgctcgagtcgggcgttccactggtaaaccca |

All primer pairs are represented from 5′ to 3′.

The forward primers for wild-type were designed to eliminate the N-terminal signal sequence predicted by the signalP6.0 server (https://services.healthtech.dtu.dk/services/SignalP-6.0/)[3].

The positions of the mutations are underlined.

Supplementary Table 4. Molecular masses and extinction coefficients of enzymes

| | Molecular mass (Da) | Extinction coefficient ($mol^{-1}cm^{-1}$) |
|---|---|---|
| CsOpgD2 WT | 57176.521 | 84340 |
| CsOpgD2 D381N | 57175.537 | 84340 |
| RpOpgD2 WT | 57032.54 | 112870 |
| RpOpgD3 WT | 58298.15 | 98445 |
| RpOpgD3 D384N | 58297.166 | 98445 |
| BdOpgD WT | 54933.177 | 83100 |
| NpOpgD WT | 52109.757 | 76360 |
| RlOpgD WT | 56435.599 | 81485 |
| RlOpgD2 D373N | 55369.509 | 81485 |
| SoOpgD2 WT | 58471.048 | 77810 |
| PeOpgD WT | 58368.586 | 86290 |
| PeOpgD D392N | 58368.602 | 86290 |
| PeOpgD2 WT | 56089.12 | 101300 |
| MiOpgD2 WT | 56795.668 | 105225 |
| MiOpgD3 WT | 57328.917 | 109905 |
| MiOpgD4 WT | 58320.984 | 84800 |

Molecular masses and extinction coefficients of purified enzymes were calculated using UV absorbance at 280 nm[4].

SupplementaryTable 5. Reaction conditions for TLC and ESI-MS analyses

| Enzyme | Concentration of linear β-1,2-glucan (%) | Protein concentration (mg/ml) | Buffer | pH | Temperature (°C) |
|---|---|---|---|---|---|
| CsOpgD2 | 1% (w/v) | 0.05 | 5mM Tris-HCl | 7 | 37 |
| RpOpgD2 | 1% (w/v) | 0.119 | 5mM Tris-HCl | 7.5 | 30 |
| RpOpgD3 | 1% (w/v) | 0.131 | 5mM Tris-HCl | 7.5 | 37 |
| BdOpgD | 1% (w/v) | 1.152 | 5mM Tris-HCl | 7.5 | 30 |
| NpOpgD | 1% (w/v) | 0.1 | 50mM NaOAc | 5 | 37 |
| RlOpgD | 1% (w/v) | 0.64 | 50mM Bis-Tris | 6 | 37 |
| SoOpgD2 | 1% (w/v) | 0.62 | 50mM Bis-Tris | 6 | 37 |
| PeOpgD | 1% (w/v) | 1.0654 | 50mM Bis-Tris | 6 | 30 |
| PeOpgD2 | 1% (w/v) | 1 | 5mM Tris-HCl | 7 | 10 |
| MiOpgD2 | 1% (w/v) | 1.096 | 5mM Tris-HCl | 7.5 | 37 |
| MiOpgD3 | 1% (w/v) | 0.522 | 5mM NaOAc | 4 | 37 |
| MiOpgD4 | 1% (w/v) | 0.458 | 5mM Tris-HCl | 7 | 30 |

Supplementary Table 6. Acetonitrile concentrations for TLC analysis and general properties

| Enzyme | For TLC | | For optimum pH | | For optimum temperature | |
|---|---|---|---|---|---|---|
| | Acetonitrile concentration (%) | Number of developments | Acetonitrile concentration (%) | Number of developments | Acetonitrile concentration (%) | Number of developments |
| CsOpgD2 | 72% (v/v) | 1 | 72% (v/v) | 1 | 72% (v/v) | 1 |
| RpOpgD2 | 75% (v/v) | 1 | 65% (v/v) | 1 | 65% (v/v) | 1 |
| RpOpgD3 | 75% (v/v) | 1 | 70% (v/v) | 1 | 70% (v/v) | 3 |
| BdOpgD | 75% (v/v) | 1 | 60% (v/v) | 1 | 65% (v/v) | 1 |
| NpOpgD | 65% (v/v) | 1 | 65% (v/v) | 1 | 65% (v/v) | 1 |
| RlOpgD | 60% (v/v) | 1 | 60% (v/v) | 1 | 62% (v/v) | 1 |
| SoOpgD2 | 65% (v/v) | 1 | 65% (v/v) | 1 | 65% (v/v) | 1 |
| PeOpgD | 72% (v/v) | 1 | 65% (v/v) | 1 | 65% (v/v) | 1 |
| PeOpgD2 | 67% (v/v) | 1 | 72% (v/v) | 1 | 72% (v/v) | 1 |
| MiOpgD2 | 72% (v/v) | 1 | 65% (v/v) | 1 | 65% (v/v) | 1 |
| MiOpgD3 | 72% (v/v) | 1 | 65% (v/v) | 1 | 65% (v/v) | 1 |
| MiOpgD4 | 65% (v/v) | 1 | 65% (v/v) | 1 | 65% (v/v) | 1 |

Supplementary Table 7. Reaction conditions for NMR

| Enzyme | Linear β-1,2-glucan (%) | Protein concentration (mg/ml) | Buffer concentration | pH | Temperature (°C) |
|---|---|---|---|---|---|
| CsOpgD2 | 3% (w/v) | 0.1 | 50mM Tris-HCl | 7 | 37 |
| RpOpgD2 | 5% (w/v) | 0.1 | 50mM Tris-HCl | 7 | 30 |
| RpOPgD3 | 5% (w/v) | 0.1 | 50mM Tris-HCl | 7 | 30 |
| PeOpgD | 5% (w/v) | 1 | 50mM Bis-Tris HCl | 6 | 30 |
| MiOpgD2 | 5% (w/v) | 0.1 | 50mM Tris-HCl | 7 | 30 |

Supplementary Table 8. Reaction conditions for Substrate specificity analyses

| Enzyme | Protein concentration (mg/ml) | Buffer concentration | pH | Temperature (℃) |
|---|---|---|---|---|
| CsOpgD2 | 0.045 | 50mM NaOAc | 5 | 37 |
| RpOpgD2 | 0.001 | 50mM NaOAc | 5 | 40 |
| RpOpgD3 | 0.005 | 50mM Tris-HCl | 7 | 40 |
| BdOpgD | 2 | 50mM NaOAc | 5 | 20 |
| NpOpgD | 0.1 | 50mM NaOAc | 5 | 37 |
| RlOpgD | 0.5 | 50mM NaOAc | 5 | 20 |
| SoOpgD2 | 1 | 50mM NaOAc | 5 | 40 |
| PeOpgD | 0.4 | 50mM Bis-Tris | 6 | 30 |
| PeOpgD2 | 1 | 50mM Tris-HCl | 7 | 10 |
| MiOpgD2 | 0.2 | 50mM Tris-HCl | 7 | 20 |
| MiOpgD3 | 0.01 | 50mM NaOAc | 5 | 30 |
| MiOpgD4 | 1 | 50mM Bis-Tris | 6 | 30 |

### Supplementary Table 9. Reaction condition of specific activity analysis

| Enzyme | Linear β-1,2-glucan (%) | Protein concentration (mg/ml) | Buffer concentration | Temperature (°C) | Time (min) |
|---|---|---|---|---|---|
| CsOpgD2 | 0.39% (w/v) | 0.25 | 20mM NaOAc (pH 5.0) | 40 | 10 |
| RpOpgD3 | 0.39% (w/v) | 0.04 | 20mM Tris-HCl (pH 7.0) | 40 | 10 |
| RlOpgD | 0.39% (w/v) | 0.25 | 20mM NaOAc (pH5.0) | 20 | 30 |

Supplementary Table 10. Conditions of crystalization

| Enzyme | Reservoir and drop | Temperature (°C) | Cryoprotectant supplemented with reservoir solution |
|---|---|---|---|
| CsOpgD2 D381N | 0.2M Potassium sodium tartrate tetrahydrate,<br>0.1M Bis-Tris propane pH 6.5,<br>20% (w/v) PEG 3350<br>[1% (w/v) linear β-1,2-glucan (average DP25)] | 20°C | 30% (w/v) glycerol<br>20% (w/v) linear β-1,2-glucan (average DP25) |
| RpOpgD3 D384N | 0.1M MIB buffer pH 8.0,<br>25% (w/v) PEG 1500<br>[1% (w/v) linear β-1,2-glucan (average DP17.7)] | 20°C | 25% (v/v) PEG 400<br>1% (w/v) linear β-1,2-glucan (average DP17.7) |
| RlOpgD D373N | 0.2 M Calcium chloride dihydrate<br>0.1M Tris-HCl pH 8.0,<br>20% v/v PEG 6000<br>[1% (w/v) linear β-1,2-glucan (average DP 17.7)] | 20°C | 25% (w/v) PEG400<br>3% (w/v) linear β-1,2-glucan (average DP 17.7) |
| PeOpgD D392N | 0.1 M Magnesium chloride,<br>0.05M Bis-Tris propane pH 6.5,<br>14% (w/v) PEG 3350<br>[1% (w/v) linear β-1,2-glucan (average DP17.7)] | 20°C | 25% (v/v) PEG 400<br>1% (w/v) linear β-1,2-glucan (average DP17.7) |

Crystals were obtained by mixing enzyme solution (7 mg/ml) and reservoir solution with the ratio of 1:1 and substrates. MIB buffer was prepared by mixing sodium malonate, imidazole, and boric acid with the molar ratio of 2:3:3.

Square brackets represent they were supplemented only in drops.

**Supplementary Note 1. Catalytic mechanism of GH186 transglycosylases**

Catalytic centers of four Michaelis complexes were compared with that of XccOpgD[5]. The reaction mechanism of XccOpgD (16) is suggested that D379 is a typical general acid, and D300 is a general base that activates 6-hydroxy group of the glucose moiety at subsite −16 (non-reducing end) via two water molecules and 4-hydroxy group of the glucose moiety at subsite −1. i.e., the cyclization of XccOpgD is catalyzed by anomer-inverting transglycosylation. In particular, the sequestration of the Grotthuss proton relay pathway at general base side by the glucose moieties at subsite −14 (the third glucose moiety from the non-reducing end) and −16 is essential for catalytic efficiency and specificity for transglycosylation activity[5]. These structural features (general acid and base, water molecules, $^{1}S_{3}$ conformation at subsite −1 and the subsites for sequestering the Grotthuss pathway) important for anomer-inverting transglycosylation were found to be conserved in two Michaelis complexes [PeOpgD (21) and RlOpgD (29)] (Supplementary Figure 17), strongly indicating that these two homologs synthesize CβGα by same mechanism with XccOpgD. In the case of Michaelis complex of CsOpgD2 (18), the electron density of water molecules at Grotthuss proton relay pathway are very poor due to rather low resolution (2.40 Å) (Supplementary Figure 17). Nevertheless, CsOpgD2 are suggested to adopt the same reaction mechanism as that of XccOpgD, considering that the features described above other than the water molecules are conserved (Supplementary Figure 17). RpOpgD3 also has almost the same features as that of XccOpgD except that the conformation of general acid (D384N) is different from that of XccOpgD (D379N) (Supplementary Figure 17). N384 does not reach the appropriate position for lateral *syn* protonation[6]. This direction is due to the interaction between N384 and S275. Amino acid sequence alignment indicates that this S275 is rarely conserved in GH186 (Supplementary Figure 13). Most of GH186 proteins have Thr at this position, implying that this direction of general acid in RpOpgD3 is very rare case which is adopted only for Ser type homologs. If the position of general acid is productive one, 6-hydroxy group of glucose moiety at subsite −1 looks vital for indirectly providing a proton to the oxygen atom of scissile glycosidic bond. D297 could be a candidate for a general acid because a proton transfer pathway via a water molecule and a 6-hydroxy group of the glucose moiety at subsite −1 is structurally observed although further analysis is needed to elucidate this pathway.

**References**


1. Tamura, K., Stecher, G. & Kumar, S. MEGA11: Molecular Evolutionary Genetics Analysis Version 11. *Mol. Biol. Evol.* **38**, (2021).
2. Robert, X. & Gouet, P. Deciphering key features in protein structures with the new ENDscript server. *Nucleic Acids Res.* **42**, 320–324 (2014).
3. Teufel, F. *et al.* SignalP 6.0 predicts all five types of signal peptides using protein language models. *Nat. Biotechnol.* **40**, 1023–1025 (2022).
4. Pace, C. N., Vajdos, F., Fee, L., Grimsley, G. & Gray, T. How to measure and predict the molar absorption coefficient of a protein. *Protein Sci* **4**, 2411–2423 (1995).
5. Motouchi, S., Komba, S., Nakai, H. & Nakajima, M. Discovery of Anomer-Inverting Transglycosylase: Cyclic Glucohexadecaose-Producing Enzyme from *Xanthomonas* , a Phytopathogen. *J. Am. Chem. Soc.* **146**, 17738–17746 (2024).
6. Nerinckx, W., Desmet, T., Piens, K. & Claeyssens, M. An elaboration on the syn-anti proton donor concept of glycoside hydrolases: Electrostatic stabilisation of the transition state as a general strategy. *FEBS Lett.* **579**, 302–312 (2005).
